# Anatomic position determines oncogenic specificity in melanoma

**DOI:** 10.1101/2020.11.14.383083

**Authors:** Joshua M. Weiss, Miranda V. Hunter, Nelly M. Cruz, Arianna Baggiolini, Mohita Tagore, Yilun Ma, Sandra Misale, Michelangelo Marasco, Theresa Simon-Vermot, Nathaniel R. Campbell, Felicity Newell, James S. Wilmott, Peter A. Johansson, John F. Thompson, Georgina V. Long, John V. Pearson, Graham J. Mann, Richard A. Scolyer, Nicola Waddell, Emily D. Montal, Ting-Hsiang Huang, Philip Jonsson, Mark T.A. Donoghue, Christopher C. Harris, Barry S. Taylor, Tianhao Xu, Ronan Chaligné, Pavel V. Shliaha, Ronald Hendrickson, Achim A. Jungbluth, Cecilia Lezcano, Richard Koche, Lorenz Studer, Charlotte E. Ariyan, David B. Solit, Jedd D. Wolchok, Taha Merghoub, Neal Rosen, Nicholas K. Hayward, Richard M. White

## Abstract

Oncogenic alterations to DNA are not transforming in all cellular contexts^1,2^. This may be due to pre-existing transcriptional programs in the cell of origin. Here, we define anatomic position as a major determinant of why cells respond to specific oncogenes. Cutaneous melanoma arises throughout the body, whereas the acral subtype arises on the palms of the hands, soles of the feet, or under the nails^3^. We sequenced the DNA of cutaneous and acral melanomas from a large cohort of human patients and found a specific enrichment for BRAF mutations in cutaneous melanoma but CRKL amplifications in acral melanoma. We modeled these changes in transgenic zebrafish models and found that CRKL-driven tumors predominantly formed in the fins of the fish. The fins are the evolutionary precursors to tetrapod limbs, indicating that melanocytes in these acral locations may be uniquely susceptible to CRKL. RNA profiling of these fin/limb melanocytes, compared to body melanocytes, revealed a positional identity gene program typified by posterior HOX13 genes. This positional gene program synergized with CRKL to drive tumors at acral sites. Abrogation of this CRKL-driven program eliminated the anatomic specificity of acral melanoma. These data suggest that the anatomic position of the cell of origin endows it with a unique transcriptional state that makes it susceptible to only certain oncogenic insults.

## Introduction

During development, cells express lineage-specific gene programs as well as position-specific gene programs. This coordinated transcriptional state ensures that cells pattern appropriately and fulfill the unique role they have at a given anatomic location^4^. Transcriptional programs unique to a particular anatomic site are referred to as positional identity^5^. Whether the transcriptional programs that mediate positional identity determines the response to oncogenes is unknown. Across many cancer types, the anatomic location of the tumor is associated with distinct molecular and clinical subtypes^6–10^. This is exemplified most strikingly by melanoma, a cancer categorized by its anatomic origin at cutaneous sites (skin of face, chest, back), acral sites (glabrous skin of the palms and soles of the hands and feet^3, 9^), mucosal sites (i.e. mouth, rectum, vagina), or within the uveal tract (the eye). Even within the skin, melanocytes residing in anatomic substructures (i.e. follicular/interfollicular or scale/interscale regions) can have markedly different susceptibilities to transformation^11, 12^. Highlighting these important clinical differences, acral melanoma, compared to other forms of cutaneous melanoma, has a unique genetic profile^6, 7, 13–15^, a lower response rate to both targeted and immunotherapy, and a worse overall survival^9, 16–18^. Using data from human patients and transgenic zebrafish models, we identify genetic drivers unique to this anatomically restricted type of melanoma and find that the positional identity gene program in the cell-of-origin determines the competence to respond to those oncogenes.

## Results

### Acral driver genes lead to anatomically distinct tumors in transgenic zebrafish

To discern the genetic differences between acral and cutaneous melanoma, we performed targeted DNA sequencing of 100 acral and 839 cutaneous melanoma patients using the MSK-IMPACT platform^19–22^, a focused sequencing panel of 468 clinically relevant genes (Fig. 1a). We calculated the log2-fold ratio of genetic alterations in acral versus cutaneous melanoma to generate an “acral enrichment score” (Fig. 1a). Consistent with previous reports^6, 7^, we found BRAF mutations as the most common coding mutation in cutaneous melanoma (Extended Data Fig. 1a). In contrast, we found that acral melanoma has a significantly higher frequency of copy number alterations (CNAs), a significantly lower frequency of activating BRAF mutations, and a lower overall mutational burden (Fig. 1a-b, Extended Data Fig. 1a-b), which has been observed in other acral melanoma genomic analyses^6, 7, 13–15^. To better define which of the frequently amplified genes were likely acral-specific drivers, we also performed RNA-sequencing (RNA-seq) on an independent set of 61 acral and 53 cutaneous melanoma patient samples. Amplification of CRKL and GAB2 were amongst the top acral enriched genes identified by a combined analysis of DNA and RNA-sequencing (Fig. 1b-c, Extended Data Fig. 1c). These were of particular interest to us since CRKL and GAB2 are both oncogenic signal amplifier proteins that together form a complex with receptor tyrosine kinases (RTKs) to amplify downstream MAPK and PI3K signaling pathways through the recruitment of signaling mediators, such as SOS1and the p85 subunit of PI3K (Extended Data Fig. 1d)^23^. They have previously been implicated in driving other cancer types^24–27^, but limited investigation has been performed in melanoma^28, 29^. Other frequently altered genes included NF1 and TERT (Fig. 1a-b, Extended Data Fig. 1a-b), which significantly co-occur with alterations in CRKL and GAB2 across cancer (Extended Data Fig. 1e). This led to the hypothesis that alterations in CRKL, GAB2, NF1, and TERT may synergize to specifically drive acral compared to cutaneous melanoma (Extended Data Fig. 1d). As an example, we identified an acral melanoma patient at Memorial Sloan Kettering Cancer Center (MSKCC) who had amplification of CRKL and GAB2, deletion of NF1, and an activating promoter mutation in TERT in both a primary tumor and metastasis (Extended Data Fig. 1f-h).

**Fig. 1:**
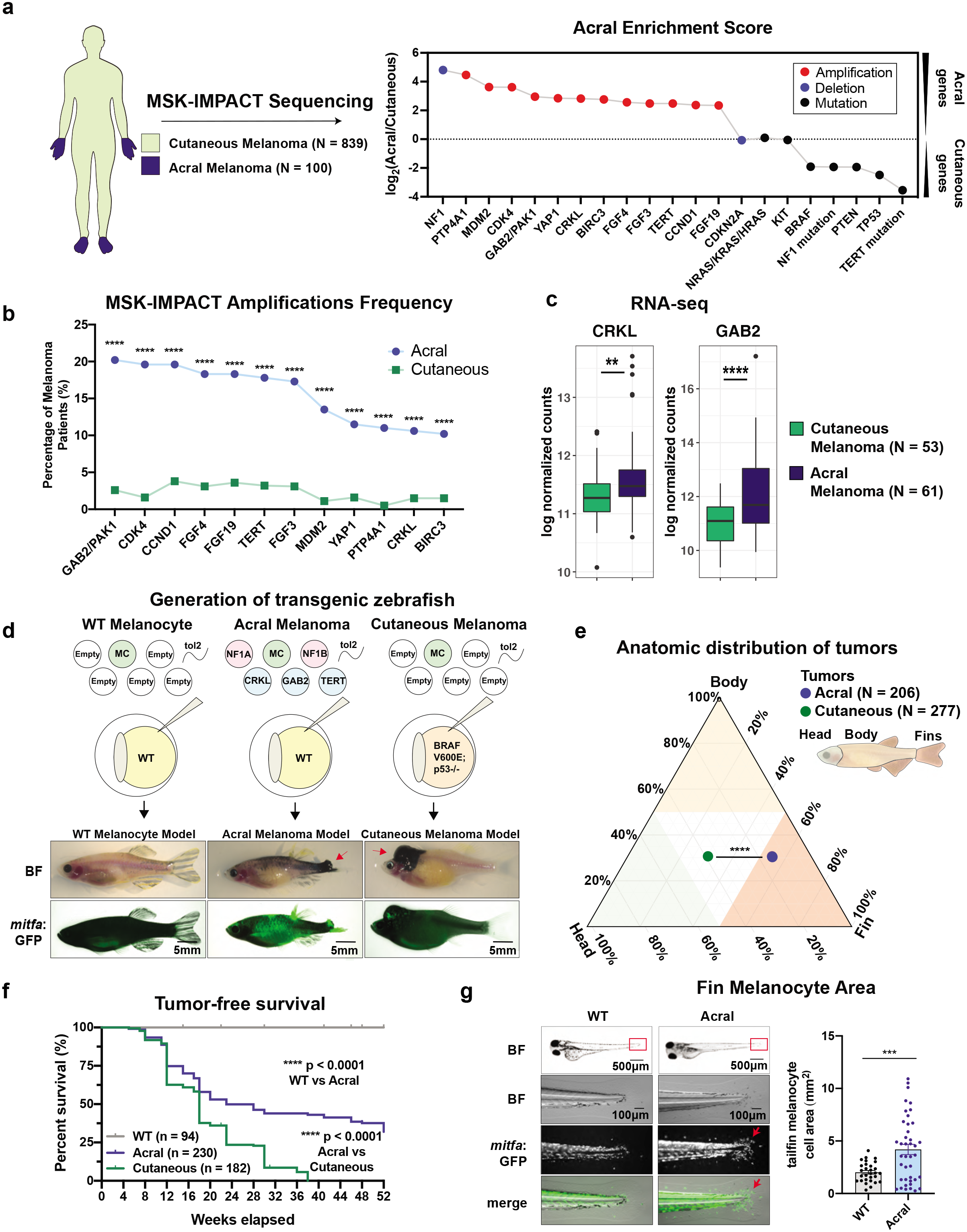
Acral versus cutaneous melanoma driver genes lead to anatomically distinct tumors in transgenic zebrafish. (a-b) MSK-IMPACT targeted sequencing of 100 acral and 839 cutaneous melanoma patients. (a) The acral enrichment score was calculated by dividing the frequency a gene is altered in acral melanoma by the frequency it is altered in cutaneous melanoma. Positive scores indicate genes enriched in acral melanoma and negative scores indicate genes enriched in cutaneous melanoma. (b) Frequency of gene amplifications compared by melanoma subtype using Fisher’s exact test. (c) RNA sequencing on a separate cohort of 61 acral and 53 cutaneous melanoma patients shows significantly higher expression of CRKL and GAB2 in acral versus cutaneous melanoma. P-values calculated using DESeq2. (d) Transgenic zebrafish models. The WT melanocyte model expresses no oncogenic drivers. The acral melanoma model has overexpression of CRKL, GAB2, TERT and knockout of zebrafish orthologues of NF1, *nf1a* and *nf1b.* The cutaneous melanoma model expresses BRAF^V600E^ in a p53^-/-^ genetic background. All transgenes are driven by the melanocyte-specific promoter *mitfa,* and melanocytes are marked with *mitfa*:GFP in all models. See Extended Data Fig.2a for more details. (e) Ternary diagram portraying the percentage of tumors arising in the head, body, or fins between the acral and cutaneous melanoma models. Anatomic distribution was compared using a Chi-squared test. See Extended Data Figure 2c for histogram representation. See Supplemental Table 3 for a full list of fish and tumor numbers across all replicates and experimental conditions. (f) Tumor-free survival of WT melanocytes (n = 94), acral (n = 230), cutaneous (n = 182) transgenic zebrafish models. P-values generated by log-rank Mantel-Cox test. (g) WT melanocyte vs acral melanoma model compared for melanocyte area in tailfin at 3-days post-fertilization. Data represents n=29 WT fish and n=41 acral fish pooled from four biological replicates. Each point represents the tailfin melanocyte area of a different animal. P-values generated by a two-sided student’s t-test. Error bars = SEM. * p-value < 0.05, ** p-value < 0.01, *** p-value < 0.001, **** p-value < 0.0001.

Since there are few widely available animal models of acral melanoma^30^, we used a rapid transgenic zebrafish system to model these potential drivers *in vivo*. Using the melanocyte-specific *mitfa* promoter, we created mosaic (F0) transgenic animals expressing putative acral melanoma drivers (CRKL, GAB2, TERT and NF1 loss) and then compared this to a previously developed cutaneous melanoma model (BRAF^V600E^;p53-/-) or to wildtype melanocytes (Fig. 1d., Extended Data Fig. 2a, Extended Data Fig. 3). We developed a rigorous set of criteria to detect tumors in both acral and cutaneous melanoma models by imaging (Supplemental Table 1) and confirmed the validity of these criteria with histology. Melanocytes in all models were fluorescently labeled with *mitfa*:GFP (Fig. 1d, Extended Data Fig. 2a-b, Extended Data Fig. 4), and darkly pigmented cells with dendritic morphology on H&E, positive immunohistochemistry (IHC) staining for GFP, RNA expression of *tyrp1a*, and immunofluorescence (IF) for *sox10* indicate that tumors generated by both models are of the melanocyte lineage (Extended Data Fig. 2b, Extended Data Fig. 3d, Extended Data Fig. 4). We validated the expression and knockout of the transgenes using a combination of DNA genotyping, CRISPR-seq, qPCR, RNA-seq, Western blot, and IHC (Extended Data Fig. 3, Extended Data Fig. 4). While wildtype melanocytes never formed melanomas, we found that 68% of the animals expressing the acral genes resulted in melanoma and 100% of the animals expressing the cutaneous genes resulted in melanoma by 1 year (Fig. 1f).

Extensive research across many fields ranging from paleontology to genetics has demonstrated an evolutionary link between the morphogenesis of fish fins and tetrapod limbs, which include the acral sites of the hands and feet^31–34^. Although fins and limbs are structurally distinct, the genes involved in fin and limb development are well conserved^32, 34^. This led us to ask whether the acral versus cutaneous driver genes would yield differences in anatomic distribution of the tumors. We monitored the fish as they developed tumors over the course of a year and calculated the relative proportion of tumors arising on the head, body, and fins. To visually aid our ability to determine the effect of genetic drivers on the anatomic distribution of tumors, we utilized the ternary plot, which compresses 3-dimensional categorical data into 2-dimensional space (Fig. 1e, Extended Data Fig. 2c)^35^. This revealed that the acral model developed a significantly higher proportion of fin tumors than the cutaneous model (53% versus 30%, p < 0.0001) (Fig. 1e, Extended Data Fig. 2c, Supplemental Table 3).

Because mosaic (F0) transgenics are more variable than stable germline transgenics, we also created stable “WT” (*mitfa*:GFP), acral melanoma (*mitfa:*CRKL;GAB2;TERT;NF1-/-), and cutaneous melanoma (*mitfa*:BRAF^V600E^;p53-/-) lines and assessed their phenotypes in the same way (Extended Data Fig. 5). In the acral stable lines, the preponderance of fin tumors was even more striking. Whereas in the mosaic acral fish 53% of tumors (n=110/206) arose on the fins, we found that 84.2% (n=96/114) of tumors from the acral stable line were fin tumors (Extended Data Fig. 5c, Supplemental Table 3). In contrast, only 39.7% (n=77/194) of tumors arose on the fins in the cutaneous stable line (Extended Data Fig. 5c, Supplemental Table 3). The anatomic position of the tumors did not shift over the 1-year observation period, suggesting that this phenotype is established early in tumorigenesis. At 3 days post-fertilization, melanocytes in the acral melanoma stable line demonstrated a greater total melanocyte area in the tailfin than melanocytes in the wild-type stable line (Fig. 1g). By 6 weeks post-fertilization the acral melanoma model developed fin hyperpigmentation and a significantly greater expansion of the melanocyte population in the fins than in the body (Extended Data Fig. 6).

### CRKL drives melanoma to acral sites

To determine more specifically which of the acral driver genes were sufficient for fin positioning, we expressed each one individually (Fig. 2a). CRKL was the only genetic driver sufficient to form tumors without any additional genetic alterations (Fig. 2b-c). Strikingly, 62% of CRKL-alone tumors arose in the fins (Fig. 2d-e, Supplemental Table 3), indicating that fin melanocytes were more efficiently transformed by CRKL than melanocytes at other locations. Tumor initiation in CRKL-only tumors was slower compared to all 4 genes together, suggesting that GAB2 and TERT overexpression with NF1 loss accelerated disease progression (Fig. 2c). We then investigated whether any of the 4 putative genetic drivers were necessary for tumor progression by removing CRKL, GAB2, TERT, or NF1 loss from the full 4-gene acral model (Fig. 2f). While tumors still formed after withdrawal of GAB2, TERT, or NF1 loss, tumors never formed after withdrawal of CRKL (Fig. 2g-h), indicating that CRKL is both necessary and sufficient for tumor formation in the acral melanoma model. Withdrawal of GAB2, TERT, or NF1 loss from the acral model still demonstrated fin specificity (Fig. 2i, Supplemental Table 3), which suggests that CRKL is solely responsible for the preponderance of acral fin tumors.

**Fig. 2:**
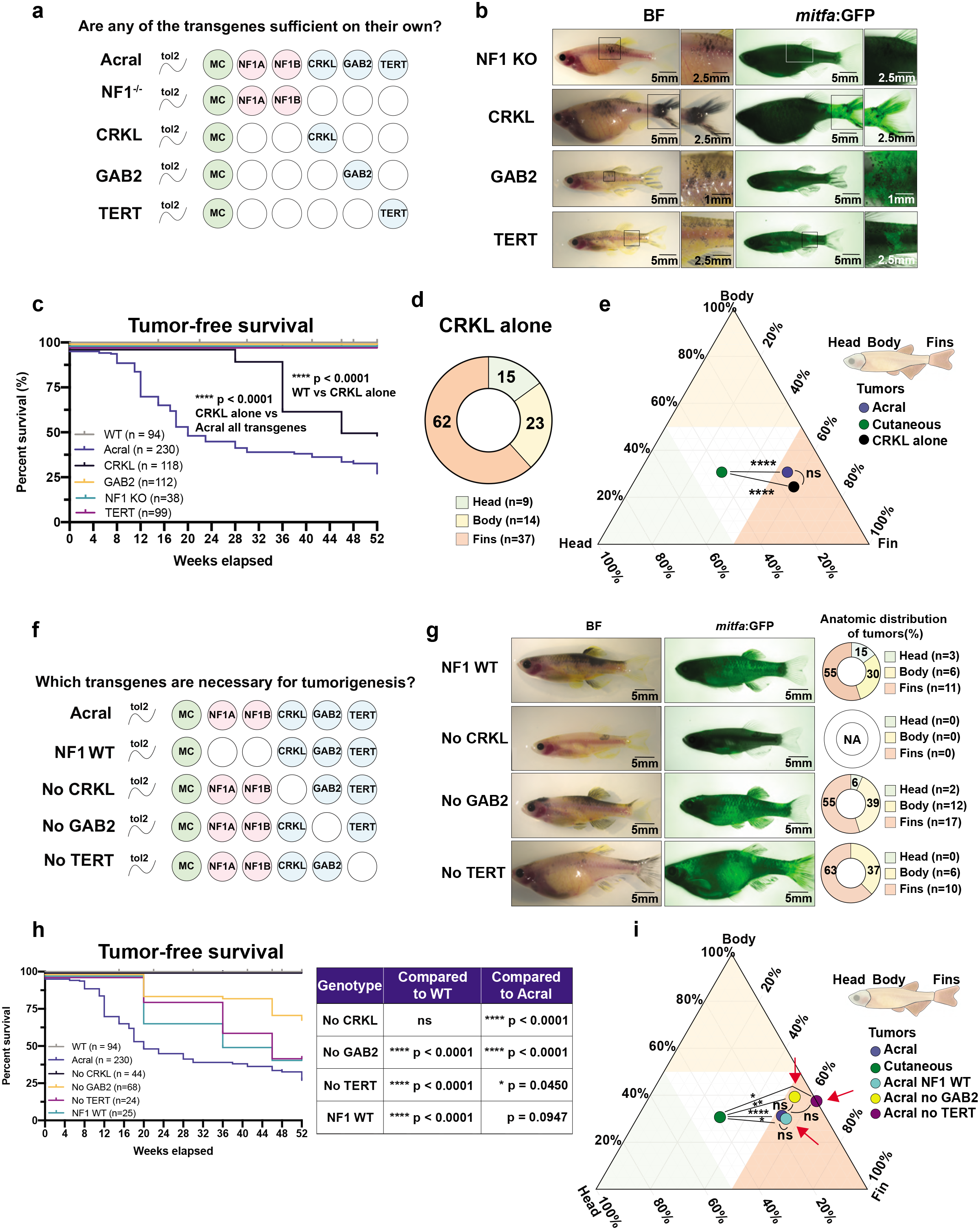
CRKL drives melanoma to acral sites. (a) The 4 genes (CRKL, GAB2, TERT, NF1 KO) used to drive the acral melanoma model were introduced separately to assess sufficiency in tumorigenesis. (b) Representative images of zebrafish for each genotype. (c) Tumor-free survival of indicated genotypes. Number of fish for each genotype indicated in figure key. P-values generated by log-rank Mantel-Cox test. (d) Pie chart showing the percentage of CRKL-alone melanomas arising at each anatomic location. The number of tumors is listed under the pie chart. (e) Ternary diagram portraying the percentage of tumors forming in the head, body, or fins of indicated genotypes. A Chi-squared test was performed to compare the anatomic distribution between the different transgenic models. See Supplemental Table 3 for a full list of fish and tumor numbers across all replicates and experimental conditions. (f) Each one of the 4 acral drive genes was removed to assess which driver genes were necessary for tumorigenesis. (g) Representative images of zebrafish for each genotype. Right panel shows pie charts with the anatomic distribution of tumors. Number of tumors are indicated. (h) Tumor-free survival of indicated genotypes. Number of fish for each genotype indicated in figure key. P-values generated by log-rank Mantel-Cox test. (i) Ternary diagram comparing the indicated genotypes. Chi-squared test was used to compare anatomic distribution between genotypes. See Supplemental Table 3 for a full list of fish and tumor numbers across all replicates and experimental conditions. * p-value < 0.05, ** p-value < 0.01, *** p-value < 0.001, **** p-value < 0.0001.

To ensure these results were not an artifact of mosaic F0 transgenesis, we also created a stable line for CRKL alone (*mitfa:*CRKL) (Extended Data Fig. 5a). Similar to what we observed in the acral stable line, the enrichment of fin tumors was even greater in the CRKL stable line. While 62% (n=37/60) of tumors arose on the fins in the mosaic CRKL fish, we found that 75.8% (n=25/33) of tumors from CRKL-only stable transgenic fish were fin tumors (Extended Data Fig. 5b-c, Supplemental Table 3). Together, these data demonstrate that CRKL is a key driver of acral melanoma and that fin melanocytes are more vulnerable to CRKL than melanocytes at other locations.

### Positional identity gene programs determine the response to CRKL

These data suggest that fin melanocytes, analogous to human hand/feet melanocytes, are uniquely sensitive to CRKL. We hypothesized that this susceptibility was due to intrinsic differences in the gene program of these melanocytes, and that this gene program was due to anatomic positioning. To address this, we performed RNA-sequencing of body versus fin melanocytes. We did this in both the acral melanoma model as well as in wild-type melanocytes. GFP+ melanocytes and GFP-microenvironment cells were isolated from adult fish using a combination of dissection and FACS (Fig. 3a). As expected, the GFP+ melanocyte population (compared to the GFP-population representing the microenvironment) showed a marked enrichment for melanocyte markers, such as *mitfa*, *pmela*, *tyr*, *tyrp1a*, *dct*, *sox10*, and *mlpha*, as well as the relevant transgenes (GFP, CRKL, etc.), confirming the successful isolation of melanocytes from both body skin and fins (Fig. 3e, Extended Data Fig. 7). We performed unsupervised hierarchical clustering on all of the samples. Surprisingly, this showed that the samples clustered first by cell lineage (melanocyte or not), second by anatomic location (fin vs body), and third by genotype (CRKL vs WT) (Fig. 3b). Along the same lines Principal Component Analysis (PCA) showed clustering by cell lineage on PC1 and by anatomic position on PC2 (Extended Data Fig. 8a). This data suggests that anatomic location contributes a greater amount of transcriptional variation than the genotype and can shape the response to oncogenes, such as CRKL.

**Fig. 3:**
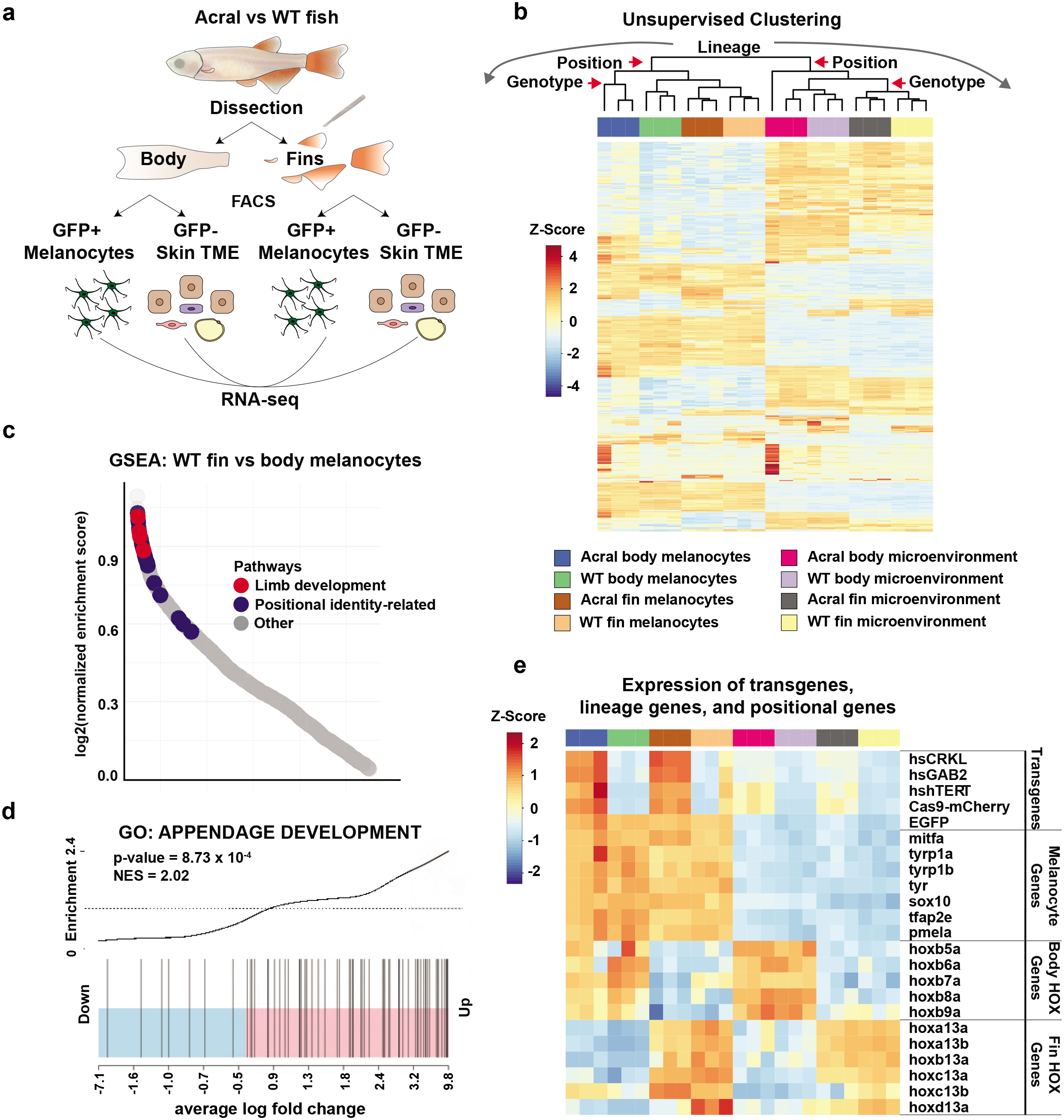
Positional identity gene programs determine the response to CRKL. (a) Schematic illustrating the fin versus body RNA-seq experiment. Body skin and fins were isolated from WT melanocyte and acral melanoma fish by dissection and then FACS sorted for GFP+ melanocytes and GFP-microenvironmental cells. (b) Unsupervised clustering using the 500 most variable genes across all samples, which clustered by cell lineage, anatomic location, and then genotype. (c) Waterfall plot representation of GSEA pathway analysis of WT fin vs body melanocytes showing the top 2500 enriched pathways and highlighting pathways related to limb development and positional identity. Limb development pathways were identified using the search terms “limb” or “appendage.” Positional-identity related pathways were identified using the search terms “morphogenesis,”, “pattern”, and “regionalization”. Only pathways with FDR < 0.05 are highlighted. The specific pathways highlighted in the plot can be found in Extended Data Fig. 8b and Supplemental Table 4. (d) GSEA barcode plot showing enrichment of genes in the GO: Appendage Development pathway. NES and FDR are indicated. (e) Heatmap representing expression of transgenes, melanocyte markers, and HOX genes across all samples.

We performed pathway analysis comparing fin versus body melanocytes, which found that 11 out of 30 of the top enriched pathways in fin melanocytes were related to limb development or anatomic position (Fig. 3c-d, Extended Data Fig. 8b, Supplemental Table 4). This indicated that the differences in the transcriptional programs of fin versus body melanocytes were largely associated with anatomic position gene programs known as positional identity. More specifically, comparing fin melanocytes to body melanocytes, we found a significant upregulation of limb-specific HOX genes in the fin melanocytes (*hoxa13a*, *hoxa13b*, *hoxb13a*, *hoxc13a*, *hoxc13b*, *hoxd13a*) whereas the body melanocytes upregulated axial-specific HOX genes (*hoxb5a*, *hoxb6a*, *hoxb7a*, *hoxb8a*, *hoxb9a*) (Fig. 3e, Extended Data Fig. 8c-d). Because FACS sorting followed by bulk RNA-seq could result in the presence of contaminating microenvironmental cells, we validated our bulk RNA-seq by performing single-cell RNA-sequencing (scRNA-seq) (Extended Data Fig. 9a-b). Similar to what we saw by bulk sequencing, there was a significant upregulation of posterior HOX13 genes such as *hoxc13b* in fin versus body melanocytes (Extended Data Fig. 9c). To avoid single-cell dropouts, we also summed the counts for all posterior HOX13 genes and similarly found a significant enrichment in fin versus body melanocytes (Extended Data Fig. 9d). scRNA-seq of the fin vs body demonstrated significantly greater expression of HOX13 genes in other cell types as well, which included fibroblasts, keratinocytes, and xanthophores (Extended Data Fig. 9e). Posterior HOX13 genes are master regulators of limb development^32^, whereas the more anterior HOXB5-9 regulate many segments of axial body development^36^. Unsupervised clustering based on expression of all detected HOX genes from bulk RNA-seq clustered samples first by anatomic location, demonstrating that positional identities of fin versus body melanocytes are associated with a HOX code (Extended Data Fig. 10).

### Human acral versus cutaneous melanoma has a positional identity gene program

We next wished to understand if these anatomic position gene programs were conserved in human acral versus cutaneous melanomas. We performed RNA-seq on human acral versus cutaneous melanoma and analyzed this by GSEA pathway analysis (Fig. 4a-b). Similar to the zebrafish models, 6 out of 30 of the top pathways in acral melanoma were related to limb development or positional identity (Fig. 4b, Extended Data Fig. 11a-b, Supplemental Table 5). Similar to what we observed in zebrafish, human acral melanoma had significantly greater expression of HOXA13, HOXB13, HOXD13 as well as other key homeobox transcription factors that regulate limb development, such as TBX4, DLX4, and HAND2 (Fig. 4c-d)^37–39^. PCA of all samples based on just HOX gene expression shows clustering by melanoma subtype (Extended Data Fig. 11c). While the majority of melanoma samples expressed HOX genes in a pattern consistent with a melanoma HOX code, 14 out of 62 acral melanomas and 16 out of 53 cutaneous melanomas did not follow this pattern, suggesting that some melanomas may lose or acquire new positional identities over the course of tumorigenesis (Extended Data Fig. 11c). To validate HOX13 expression on the protein level, we used an acral melanoma tissue microarray (TMA) developed at Sloan Kettering with n=32 patient samples, and then performed IHC for HOXB13 and CRKL (Fig. 4e, Extended Data Fig. 12a-c). Analysis by a dermatopathologist revealed that 90% were positive for HOXB13 and 53.3% positive for CRKL (Fig. 4e, Extended Data Fig. 12a-b). Of note acral tumors staining positively for HOX13 included both primary (n=8/9) and metastatic (n=19/21) samples, indicating that positional identity is likely retained at distant metastatic sites (Fig. 4e, Extended Data Fig. 12a-b). Because many genes other than HOX genes are differentially expressed between acral and cutaneous melanoma, we also performed HOMER motif enrichment analysis on the RNA-seq data^40–42^ to identify which transcription factors regulate the broader set of differentially expressed genes. A motif for HOXB13 was one of the top motifs indicating that HOX genes may regulate many of the transcriptional differences between the two subtypes (Fig. 4f).

**Fig. 4:**
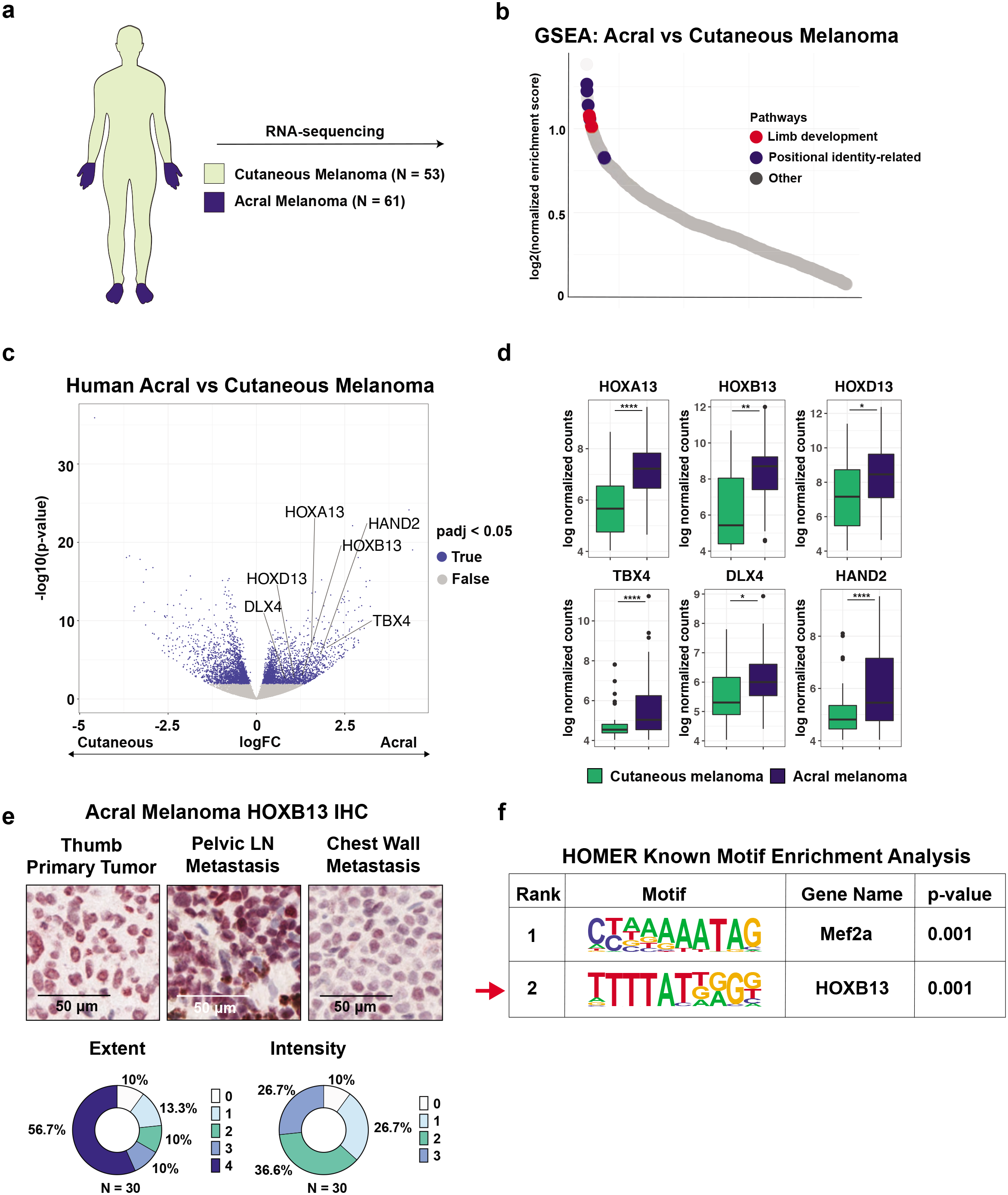
Human acral versus cutaneous melanoma has a positional identity gene program. (a) RNA-seq performed on 61 acral and 53 cutaneous melanoma human patient samples. (b) Waterfall plot representation of GSEA pathway analysis of human acral vs cutaneous melanoma showing the top 2000 enriched pathways and highlighting pathways related to limb development and positional identity. GSEA analysis is controlled for disease stage. Limb development pathways were identified using the search terms “limb” or “appendage.” Positional-identity related pathways were identified using the search terms “morphogenesis,”, “pattern”, and “regionalization.” Only pathways with FDR < 0.05 are highlighted. The specific pathways highlighted in the plot can be found in Extended Data Fig. 11a and Supplemental Table 5. (c) Volcano plot showing differentially expressed genes between acral and cutaneous melanoma samples. Genes with FDR-adjusted p-value < 0.05 indicated in blue. P-values calculated using DESeq2. (d) Boxplots showing differences in gene expression between acral melanoma and cutaneous melanoma samples. P-values adjusted for FDR = 0.05 indicated on the plot. Error bars = SEM. P-values calculated using DESeq2. (e) Acral melanoma tissue microarray underwent IHC for HOXB13. Representative images of 3 samples are shown. N=30 samples were independently scored by a dermatopathologist for extent and intensity, which are represented by pie charts. (f) Genes differentially expressed between acral and cutaneous melanoma samples underwent HOMER known motif enrichment analysis for predicted transcription factor binding motifs. * p-value < 0.05, ** p-value < 0.01, *** p-value < 0.001, **** p-value < 0.0001.

### Limb-specific HOX13 genes regulate IGF signaling in acral melanocytes

This data suggested that oncogenic drivers such as CRKL might synergize with transcriptional targets of limb-specific HOX13 genes. To identify the genes regulated by HOX13 in the limbs, we analyzed HOX13 ChIP-seq data from developing mouse limb bud tissue^43^ (Fig. 5a). We used Cistrome-GO^44^ to perform pathway analysis for genes with promoters or enhancers occupied by HOX13 and found that IGF/insulin signaling, a key regulator of limb development and regeneration, was amongst the topmost dysregulated pathways^45–48^ (Fig. 5b-c, Extended Data Fig. 13a-b). We observed significant binding of HOXA13 and HOXD13 at multiple sites along the promoters of both IGF1 and IGF2 indicating that HOX13 genes regulate IGF (Extended Data Fig. 13c-d). We also found significant H3K27 acetylation peaks at HOX13 binding sites along the promoters of IGF1 and IGF2, indicating that HOX13 likely is associated with IGF1 and IGF2 expression (Extended Data Fig. 13c-d). Consistent with these data, we found that one of the top enriched pathways in fin versus body melanocytes was regulation of multicellular organism growth, where fin melanocytes have significantly elevated expression of *igf1*, *igf2a*, and *igf2b* compared to the body melanocytes (Fig. 5d, Extended Data Fig. 13e-f).

**Fig. 5:**
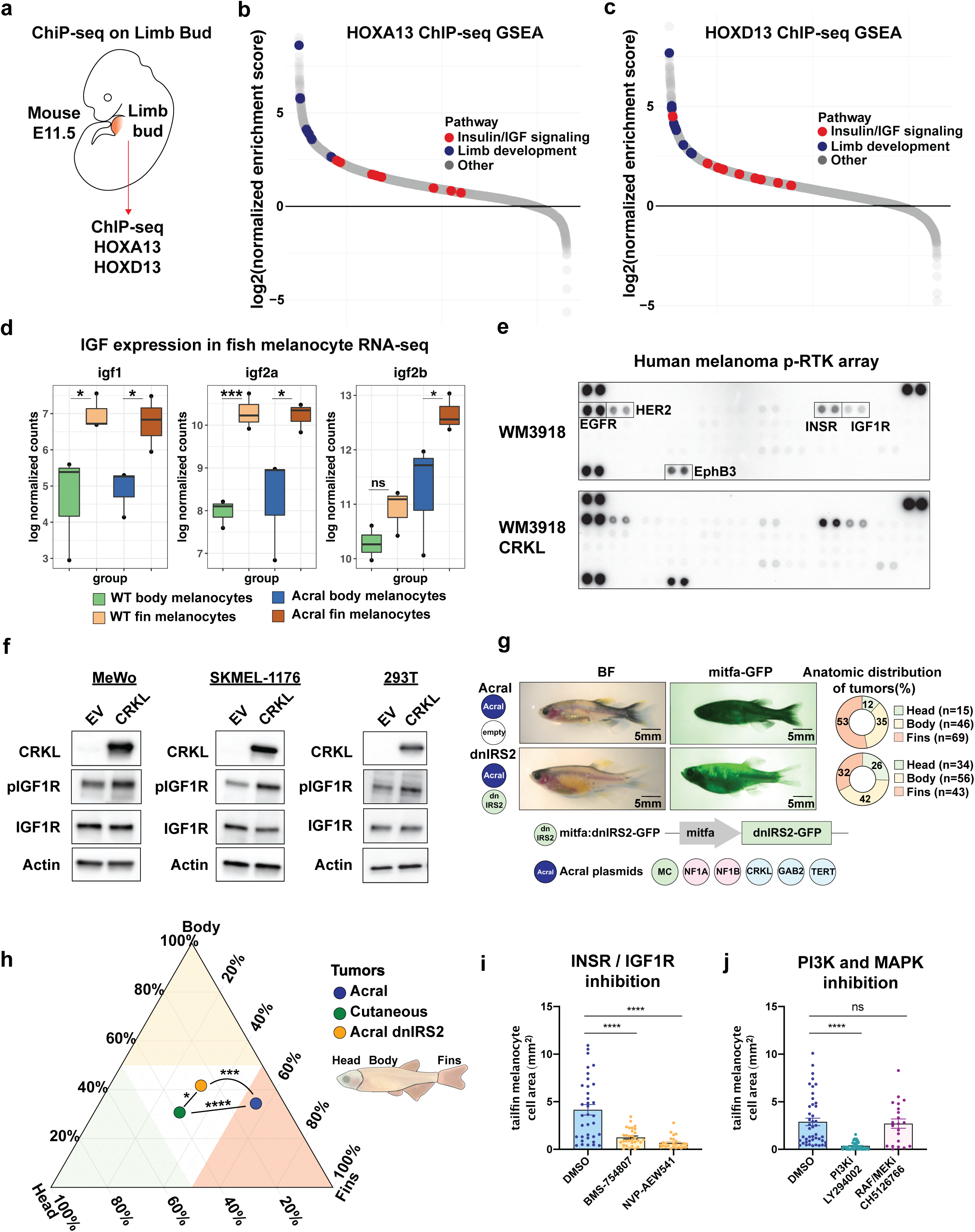
HOX13 synergizes with CRKL to drive acral melanoma through insulin/IGF signaling. HOXA13 and HOXD13 ChIP-seq performed on developing limb buds from E11.5 mouse embryos^43^. (b-c) Waterfall plot representation of 5594 and 5625 top enriched GSEA pathways regulated by HOXA13 and HOXD13 binding, highlighting limb and insulin/IGF-related pathways. Limb development pathways were identified using the search terms “limb” or “appendage.” Insulin/IGF signaling pathways were identified using the search terms “insulin” or “IGF”. Only pathways with FDR < 0.05 are highlighted. The specific pathways highlighted in the plot can be found in Extended Data Fig. 13a-b and Supplemental Table 6. (d) Boxplot showing log normalized counts of zebrafish *igf1*, i*gf2a*, and *igf2b* expression in all melanocyte samples from bulk RNA-seq. P-values are adjusted for FDR = 0.05. P-values calculated using DESeq2. (e) Phospho-RTK array performed on WM3918 human melanoma cell line with or without overexpression of CRKL. Array tests phosphorylation status of 49 RTKs. (f) MeWo, SKMEL-1176, and 293T cells were transduced and selected to overexpress CRKL or an empty vector (EV) as a negative control. Western blot for CRKL, pIGF1R, and total IGF1R was then performed to compare basal levels of IGF signaling. Actin was used as a loading control. See Extended Data Fig. 15e for quantification. (g) Schematic of dnIRS2 transgene used to block insulin/IGF signaling in zebrafish melanocytes. Representative images of acral melanoma model with or without overexpression of dnIRS2-GFP. Pie charts demonstrate the anatomic distribution of each genotype. Number of tumors is indicated. See Extended Data Fig. 18a to see tumor-free survival curve. (h) Ternary plot showing the anatomic distribution of tumors with indicated genotypes. P-values generated by Chi-squared test. See Supplemental Table 3 for a full list of fish and tumor numbers across all replicates and experimental conditions. (i-j) Acral melanoma model imaged for melanocyte tailfin area at 3-days post-fertilization after indicated pharmacologic treatment. P-value generated with two-sided student’s t-test. Error bars = SEM. (i) Insulin/IGF1 receptor antagonists BMS-754807 at 7.5μM and NVP-AEW541 at 60μM compared to 0.1% DMSO control. Data represents n=35 DMSO-treated fish, n=31 BMS-treated fish, and n=26 NVP-treated fish pooled over three biological replicates. See Extended Data Fig. 18b for representative images. (j) PI3K inhibitor LY294002 at 15μM and RAF/MEK inhibitor CH5126766 at 1μM compared to 0.1% DMSO control. Data represents n=45 DMSO-treated fish, n=43 PI3K inhibitor-treated fish, and n=21 RAF/MEK inhibitor-treated fish pooled over three biological replicates. See Extended Data Fig. 18c for representative images. * p-value < 0.05, ** p-value < 0.01, *** p-value < 0.001, **** p-value < 0.0001.

Since the mouse ChIP-seq experiment was performed on whole limb bud tissue, we also wanted to determine whether this HOX13/IGF relationship specifically existed in human melanoma cells. We first performed Western blot to identify HOX13+ patient-derived melanoma cell lines. From this analysis we identified 3 cell lines, 2 of which are confirmed to be derived from acral melanoma patients (SKMEL-1176 and SKMEL-1206) and the other of unknown anatomic origin (SKMEL-1088) (Extended Data Fig. 15a). We performed CUT&RUN using an antibody for HOXA13 on those 3 melanoma cell lines (SKMEL-1088, SKMEL-1176 and SKMEL-1206), along with an IgG negative control. HOMER motif analysis correctly found the HOX13 motif to be significantly enriched in the CUT&RUN data from all 3 cell lines (Extended Data Fig. 14a). Cistrome-GO pathway analysis identified insulin receptor substrate binding as the most enriched pathway and many pathways related to insulin/IGF signaling were also amongst the top 20 enriched pathways (Extended Data Fig. 14b). Examination of individual genes showed significant HOXA13 binding at the IGF1R promoter, along with a peak approximately 2.8kb 5’ to the IGF2 promoter as well as a peak 50kb 3’ to IGF2 (Extended Data Fig. 14c-d). Other IGF signaling related genes, such as IRS1, IRS2, and IGFBP3, demonstrated peaks near the transcription start site, indicating HOX13 regulation of multiple components of the pathway (Extended Data Fig. 14e-g).

In order to determine whether HOX13 binding peaks identified by Cut & Run functionally regulate expression of IGF1R and IGF2, we performed siRNA knockdown of HOX13 genes in HOX13^HI^ cell lines. We used siRNAs against all 4 human HOX13 genes (HOXA13, HOXB13, HOXC13, HOXD13) in the SKMEL-1176 and SKMEL-1206 acral melanoma cell lines and validated knockdown by Western blot for HOXB13 (Extended Data Fig. 15b-c). This resulted in a significant decrease in IGF2 ligand, total IGF1R, and pIGF1R after HOX13 knockdown (Extended Data Fig. 15b-c), consistent with HOX13 genes being a transcriptional activator of IGF2 and IGF1R. In order to demonstrate that HOX13 was regulating expression of functional IGF ligand, we additionally performed ELISA for secreted IGF2 from the conditioned media of SKMEL-1088 and SKMEL-1176 cells and observed significant decrease in secreted IGF2 following HOX13 knockdown (Extended Data Fig. 15d).

To further investigate this relationship, we performed Western blot for IGF1R and pIGF1R, a marker of IGF pathway activation, on human acral and cutaneous melanoma cell lines (Extended Data Fig. 15a). Supporting the CUT&RUN data, HOXB13^HI^ cells not only had higher expression of the IGF1R receptor, but also demonstrated higher levels of pIGF1R, indicating greater overall pathway activation (Extended Data Fig. 15a). In addition to observing high IGF signaling in acral melanoma cell lines, we also observed pathway activation in patient tumors, whereby 96.7% of samples from the Sloan Kettering acral melanoma TMA stained positive for pIGF1R (Extended Data Fig. 12a-b). Together this data suggests a direct role for HOX13 in regulating IGF signaling in melanoma.

### CRKL amplifies HOX13/IGF signaling

CRKL is an adapter protein that builds protein complexes that then mediate downstream signaling. The nature of those complexes is what determines what signaling pathways are most affected, which can vary across tumor types^23^. Previous work has shown that CRKL can integrate signaling through IGF1R/PI3K^49–51^, raising the hypothesis that CRKL was amplifying the HOX13/IGF axis to promote acral melanoma. To test this, we performed a phospho-RTK array of 49 different RTKs in WM3918 cells, a human NF1-null melanoma cell line with or without overexpression of CRKL (Fig. 5e). CRKL overexpression specifically increased the phosphorylation of the insulin and IGF1 receptors (INSR/IGF1R) (Fig. 5e), consistent with increased activation of these RTKs in the setting of CRKL overexpression. We confirmed this effect in 3 additional cell lines (SKMEL-1176, MeWO, and 293T) in which CRKL overexpression induced increased levels of pIGF1R seen by Western blot (Fig. 5f, Extended Data Fig. 15e). To characterize the protein complex formed by CRKL and identify its binding partners, we performed IP-mass spec (Extended Data Fig. 16). We expressed a V5-tagged version of CRKL in the WM3918 human melanoma cells used above and then pulled down with an anti-V5 antibody, followed by mass spectrometry. We found that CRKL bound multiple subunits of PI3K including p110 subunits (PIK3CB and PIK3CD) as well as p85 subunits (PIK3R2), the major downstream mediator of IGF/IGF1R signaling (Extended Data Fig. 16). In addition, we also pulled down many GTPase-activating proteins (GAPs) of Rac/Rho GTPases, such as ARAP1^52^, ASAP1^52^, ARHGAP32^53^, and guanine nucleotide exchange factors (GEFs) of Rac/Rho, such as C3G^53^, DOCK1^54, 55^, DOCK5^55^, and the DOCK co-activators, ELMO1^54^ and ELMO2^55^. DOCK1 and ELMO1 have been previously shown to activate PI3K signaling specifically through PIK3CB^54^ (Extended Data Fig. 16) and ARAP1^52^ is activated by PIP3, the product of PI3K. In contrast to this heavily enriched PI3K-centered mechanism, we found relatively few members of the MAPK pathway (SOS1/SOS2) (Extended Data Fig. 16). Given CRKL’s well known ability to mediate RTK signaling, these data imply that somatic amplification of CRKL in acral melanoma synergizes with the existing HOX13/IGF positional identify program, and this is likely mediated via direct binding to PI3K family members.

To further examine this question, we performed detailed histological analysis of our zebrafish transgenic models (Extended Data Fig. 4). We stained n=9 fish (3 wild-type, 3 acral, 3 cutaneous) using H&E as well as antibodies for GFP, CRKL, pIGF1R, pERK, and pS6 (note: available HOX13 antibodies do not work in zebrafish) (Extended Data Fig. 4). In the acral zebrafish tumors, there was a clear and striking overlap between GFP (indicating the presence of the transgenic melanocytes), CRKL, pIGF1R, and pS6 (Extended Data Fig. 4). Interestingly, the acral fish tumors also stained for pERK (Extended Data Fig. 4), yet we saw little evidence for sensitivity to MAPK inhibition (as discussed below). In contrast, the cutaneous fish tumors were negative for CRKL, faintly positive for pIGF1R (similar to control animals), strongly positive for pERK (as expected), and only 1 animal was positive for pS6 (Extended Data Fig. 4). Together this data supports a model whereby CRKL amplifies IGF signaling in fin melanocytes through its interaction with PI3K.

### Abrogation of HOX13/IGF signaling eliminates anatomic positioning of acral melanoma

This data raised the possibility that IGF signaling, downstream of the intrinsic HOX13 program, might synergize with CRKL and explain why that oncogene is enriched at acral sites. To test this, we used CRISPR to knockout both HOX13 and IGF genes in our transgenic zebrafish CRKL model. Zebrafish have 6 HOX13 genes (*hoxa13a*, *hoxa13b*, *hoxb13a*, *hoxc13a*, *hoxc13b*, *hoxd13a*) and 3 IGF genes (*igf1*, *igf2a, igf2b*), so we created a vector which allowed us to co-express 3 sgRNAs at a time, and we co-injected several of these to knock out all 6 HOX13 genes or all 3 IGF genes simultaneously in a mosaic (F0) fashion (Extended Data Fig. 17a,e). By co-injecting these plasmids with a *mitfa*:Cas9 plasmid, we were able to ensure that knockout occurred in a melanocyte specific manner. As a positive control, we first injected sgRNAs against CRKL itself, and then measured the tailfin melanocytes as we had done previously (Fig. 1g) and found that this decreased fin melanocyte expansion (Extended Data Fig. 17a-d). As a negative control, we used a sgRNA to knockout *hoxb7a* (an anterior HOX gene not expressed in the fin melanocytes), and consistent with our hypothesis, knockout of *hoxb7a* had no effect on the CRKL-induced tailfin phenotype (Extended Data Fig. 17e-g). We then tested the effect of knocking out all 6 HOX13 genes in the CRKL-overexpressing fish. Surveyor assays confirmed targeting of all 6 loci (Extended Data Fig. 17f), and fluorescent imaging showed a significant rescue of the CRKL-induced tailfin melanocyte expansion (Extended Data Fig. 17g). To directly test the role of IGF, we used sgRNAs against the fish IGF genes (*igf1*, *igf2a*, *igf2b*) and, similar to what was seen with HOX13 knockout, we found a significant decrease in the total area occupied by tailfin melanocytes (Extended Data Fig. 17a-d). To test this pathway more directly in adult tumors, we overexpressed a previously validated dominant negative IRS2 (dnIRS2) to block insulin/IGF signaling in the CRKL-driven acral melanoma model^56, 57^ (Fig. 5g). We found that the dnIRS2 modestly slowed tumor initiation (Extended Data Fig. 18a, Supplemental Table 3), but most importantly led to a robust loss of fin specificity in the acral melanoma model (Fig. 5h, Supplemental Table 3).

Finally, we also tested whether pharmacologic inhibition of the IGF/PI3K pathway would block the fin phenotype of the acral melanoma model. Treatment of the acral melanoma model with INSR/IGF1R antagonists (BMS-754807 and NVP-AEW541)^57–59^ led to a significant decrease in the total melanocyte area in the tailfin (Fig. 5i, Extended Data Fig. 18b). Along the same lines, we also tested the effects of the PI3K inhibitor LY294002^60^, and found a robust decease in the total melanocyte area in the tailfin (Fig. 5j, Extended Data Fig. 18c). In contrast, treatment with the RAF/MEK inhibitor CH5126766^61^ or MEK inhibitors trametinib^62^, pimasertib^63^, refametinib^64^, or the SOS1 inhibitor, Bl-3406^65^, only led to a very modest or insignificant rescue of the tailfin phenotype (Fig 5j, Extended Data Fig. 18c, Extended Data 19). This data is consistent with our observation that only a few MAPK members (SOS1/SOS2) were enriched in our IP-mass spec data, in contrast to the multiple PI3K family members, which suggests a greater dependence on PI3K signaling (Extended Data Fig. 16). Another possibility is that the chromatin state of acral melanocytes is not optimal for responding to activation of MAPK signaling, as we recently demonstrated^66^. Together, these data demonstrate that CRKL-mediated transformation depends on a HOX13/IGF/PI3K associated limb positional gene program (Extended Data Fig. 20).

## Discussion

What are the critical factors necessary for a cell to become a cancer? Many studies have documented phenotypically normal cells with classic oncogenic driver mutations^1, 2^, suggesting that tumorigenesis depends not just on mutations but also on a particular cell state. Here we introduce positional identity, a transcriptional program linked to a particular anatomic location^5^, as a key determinant of the transforming potential of oncogenes in cancer. Through a combination of DNA and RNA-sequencing of human patients, we identified CRKL amplification as specifically enriched in acral melanoma, a subtype defined by its anatomic position on the hands and feet^3^. Due to the evolutionary relationship between zebrafish fins and human hands and feet^32, 33^, we were able to establish the first *in vivo* model of acral melanoma. We found that fin melanocytes were more susceptible to CRKL transformation than melanocytes at other anatomic sites. Mechanistically we found that fin melanocytes have a limb positional identity defined by HOX13 genes, which drive higher levels of IGF signaling. We identified the same positional identity programs in human acral melanoma. CRKL potentiates this HOX13/IGF axis by forming a protein complex with multiple PI3K family members (PIK3CB, PIK3CD, PIK3R2), the major downstream signaling pathway enacted by IGF/IGF1R. This leads to a model in which somatic amplifications of CRKL synergize with a limb positional identity program to drive melanoma most efficiently at acral anatomic sites (Extended Data Fig. 20).

Our genomic analysis of acral and cutaneous melanoma using the targeted sequencing platform, MSK-IMPACT, is consistent with previous genomic analyses of acral melanoma^6, 7, 13–15^. Similar to previous reports, we also found a high frequency of gene amplifications for GAB2/PAK1, CCND1, TERT, MDM2, and CDK4^6, 7, 13–15^. Our analysis is the first to report amplification of CRKL as a driver of acral melanoma. There is an independent acral genomic analysis in press also finding a high frequency of CRKL amplifications^67^, thereby validating our results. The advantage of the MSK-IMPACT platform is the standardization of the pipeline^19–22^, which facilitates comparing the frequency of genetic alterations between cancer subtypes. This in combination with RNA-seq data also directly comparing acral to cutaneous melanoma from an independent cohort of patients was key in separating likely driver from passenger alterations and specifically to identify CRKL.

This study provides a conceptual framework to understand previous work that has documented spatial differences in susceptibility to tumorigenesis. On the dorsal skin of mice regional differences have been observed between follicular and interfollicular melanocytes in their ability to respond to oncogenic drivers^11, 12^. Furthermore, pigmented melanocytes in the scale region of the tail, but not the interscale region, could give rise to tumors despite both populations expressing BRAF^V600E^ and loss of PTEN^12^. While the authors attribute this to regional differences in the microenvironment^12^, it is also possible these melanocytes intrinsically have distinct transcriptional programs based on their location. While our study demonstrates how cell intrinsic positional programs can determine oncogenic specificity in tumorigenesis, we do not rule out the important role of the microenvironment in this process. Our scRNA-seq data suggests that not just melanocytes, but also microenvironmental cells, such as keratinocytes and fibroblasts, express higher levels of HOX13 genes in the fins versus the body skin of zebrafish (Extended Data Fig. 9e). Whether the positional identity of microenvironmental cells might also contribute to tumorigenesis is an area of future investigation.

Positional identity can address a longstanding question, which is why do cancers arising at different locations often possess distinct genetic and clinical characteristics? Examples include not just other skin cancers like basal and squamous cell carcinoma^2^, but also colon cancer^8^, gastric cancer^68^, cholangiocarcinoma^69^, and glioma^70^. While the HOX13-IGF-CRKL mechanism we propose may be specific to subsets of patients with acral melanoma, it is likely that oncogenes other than CRKL can interact with the HOX13 program. For example, we and others have also identified significant enrichment in genes such as PAK1, CDK4 and YAP1 in acral melanoma^13, 14^ (Fig.1a-b), and how they intersect with positional identity is an important area for future study. The concept that oncogenes interact with location-specific programs in the cell of origin may apply to a variety of cancer types, which may lead to new opportunities in the treatment of cancer. Future studies to comprehensively characterize the positional dependencies of oncogenes will be a valuable resource in elucidating the origins of cancer and treating cancer based on its positional identity.

## Supporting information

Table S1 - CAMP criteria for melanoma diagnosis in transgenic zebrafish models

Table S2 - Primer Sequences

Table S3 - Tumor Incidence and Anatomic Distribution for all Transgenic Models

Table S4 - Zebrafish Fin vs Body RNA-seq

Table S5 - Human Acral vs Cutaneous RNA-seq

Table S6 - Limb Bud HOX13 ChIP-seq Pathway Analysis

Table S7 - Human Melanoma HOXA13 Cut & Run

Table S8 - CRKL IP Mass Spec

## Acknowledgements

J.M.W. was supported by the Kirschstein-NRSA predoctoral fellowship (F30) from the NIH under award number F30CA236442 and by a predoctoral fellowship (T32) by the Cell and Developmental Biology Program at Weill Cornell Graduate School supported by the the NIH under award number T32GM008539. N.R.C was supported by the Kirschstein-NRSA predoctoral fellowship (F30) from the NIH under award number F30CA220954. J.M.W., Y.M., and N.R.C were supported by a Medical Scientist Training Program grant from the NIH under award number T32GM007739. A.B. was supported by the Swiss National Science Foundation Postdoc.mobility fellowships P2ZHP3_171967 and P400PB_180672. M.T. was supported by the Marie-Josée Kravis Women in Science Endeavor Postdoctoral Fellowship. E.D.M. was supported by the Predoctoral to Postdoctoral Fellow Transition Award (F99/K00) from the NIH under award number K00CA223016. R.A.S., G.V.L., J.F.T., G.J.M. and N.K.H. are supported by an Australian National Health and Medical Research Council Program Grant and R.A.S., N.K.H. and G.V.L. are supported by the Australian National Health and Medical Research Council Fellowship program; these authors gratefully acknowledge support from colleagues at their respective institutions and support from the Cameron Family and The Ainsworth Foundation. This research was funded in part through the NIH/NCI Cancer Center Support Grant P30 CA008748, Ludwig Collaborative and Swim Across America Laboratory, Memorial Sloan Kettering Cancer Center, New York, NY 10065, USA; Parker Institute for Cancer Immunotherapy, Memorial Sloan Kettering Cancer Center, New York, NY 10065, USA; Department of Medicine, Memorial Sloan Kettering Cancer Center, New York, NY 10065, USA; Weill Cornell Medicine, New York, NY 10065, USA. This work was supported by awards from the Melanoma Research Alliance, The Debra and Leon Black Family Foundation, NIH Research Program Grants R01CA229215 and R01CA238317, NIH Director’s New Innovator Award DP2CA186572, The Pershing Square Sohn Foundation, The Mark Foundation, The Alan and Sandra Gerry Metastasis Research Initiative at the Memorial Sloan Kettering Cancer Center, The Harry J. Lloyd Foundation, Consano and the Starr Cancer Consortium (all to R.M.W.). Thank you to Lambros T. Koufariotis for technical assistance with the alignment of human RNA-seq data. We acknowledge the use of the Integrated Genomics Operation Core, funded by the NCI Cancer Center Support Grant (CCSG, P30 CA08748), Cycle for Survival, and the Marie-Josée and Henry R. Kravis Center for Molecular Oncology.

## Author Contributions

Conceptualization, J.M.W. and R.M.W; Methodology, J.M.W., M.V.H, M.T., Y.M., S.M., T.S.V, N.R.C., F.N., T.H.H., T.X., P.V.S., A.A.J., R.K.; Software, J.M.W., M.V.H., N.R.C., F.N., R.K.; Formal Analysis, J.M.W, M.V.H., M.T., M.M., N.R.C., P.J., C.C.H., M.D., B.S.T., T.X., R.C., P.V.S., R.H., C.L., R.K., D.B.S., N.K.H.; Investigation, J.M.W., M.V.H., N.M.C., A.B., M.T., Y.M., S.M., M.M., T.S.V., E.D.M., T.H.H., P.V.H., R.H., A.A.J., C.L., R.K., L.S., Resources, R.M.W.; Data Curation, J.M.W., M.V.H, M.T., N.R.C, F.N., J.S.W., P.A.J., J.F.T., G.V.L., J.V.P., G.J.M., R.A.S., N.W., T.X., P.V.S., R.K. P.J., C.C.H., M.D., B.S.T, C.E.A., D.B.S, J.D.W., T.M., and N.K.H.; Writing – Original Draft, J.M.W. and R.M.W.; Writing – Review & Editing, J.M.W., M.V.H., S.M., C.E.A., and R.M.W.; Visualization, J.M.W; Supervision, C.E.A., D.B.S., J.D.W., T.M., N.R., and R.M.W.; Project Administration, J.M.W., C.E.A., D.B.S., J.D.W., T.M., N.R., and R.M.W.; Funding Acquisition, R.M.W.

---

**Extended Data Fig. 1:**
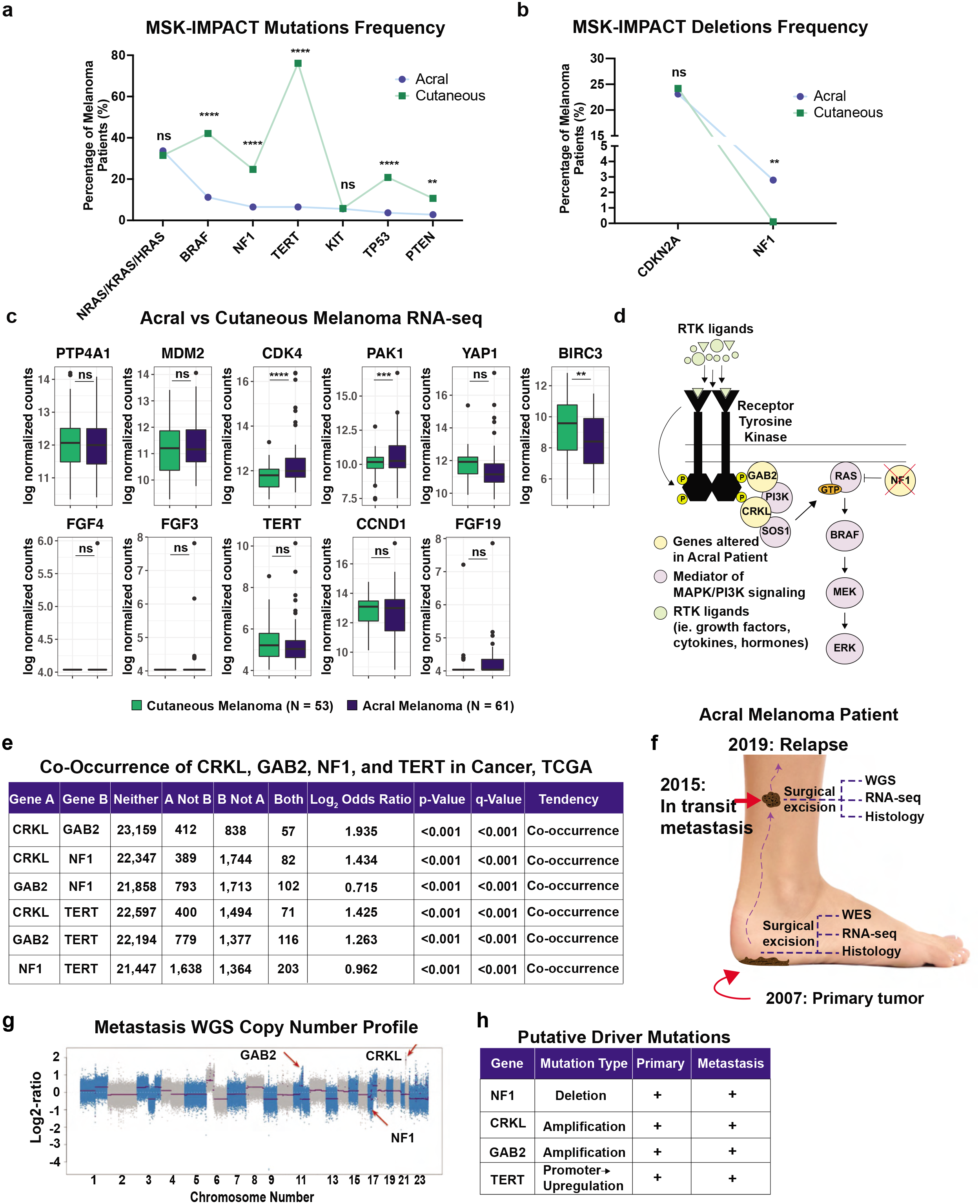
Identification of acral versus cutaneous melanoma genes. Related to Fig. 1. (a-b) MSK-IMPACT of acral and cutaneous melanoma. Fisher’s exact test was used to compare the frequency of the most recurrently mutated and deleted genes by melanoma subtype. Both coding and promoter mutations were counted for TERT. (c) RNA-seq of 61 acral and 53 cutaneous melanoma patient samples. Boxplots compare indicated genes by subtype. Error bars = SEM. P-values calculated using DESeq2. (d) Schematic detailing predicted synergistic interaction between putative driver genes in acral melanoma. (e) TCGA pan-cancer analysis shows significant co-occurrence of CRKL, GAB2, NF1, and TERT alterations. (f) Clinical course of an acral melanoma patient. (g) WGS copy number profile showing copy number changes in GAB2, CRKL, and NF1. (h) Putative drivers of patient’s acral melanoma. * p-value < 0.05, ** p-value < 0.01, *** p-value < 0.001, **** p-value < 0.0001.

**Extended Data Fig. 2:**
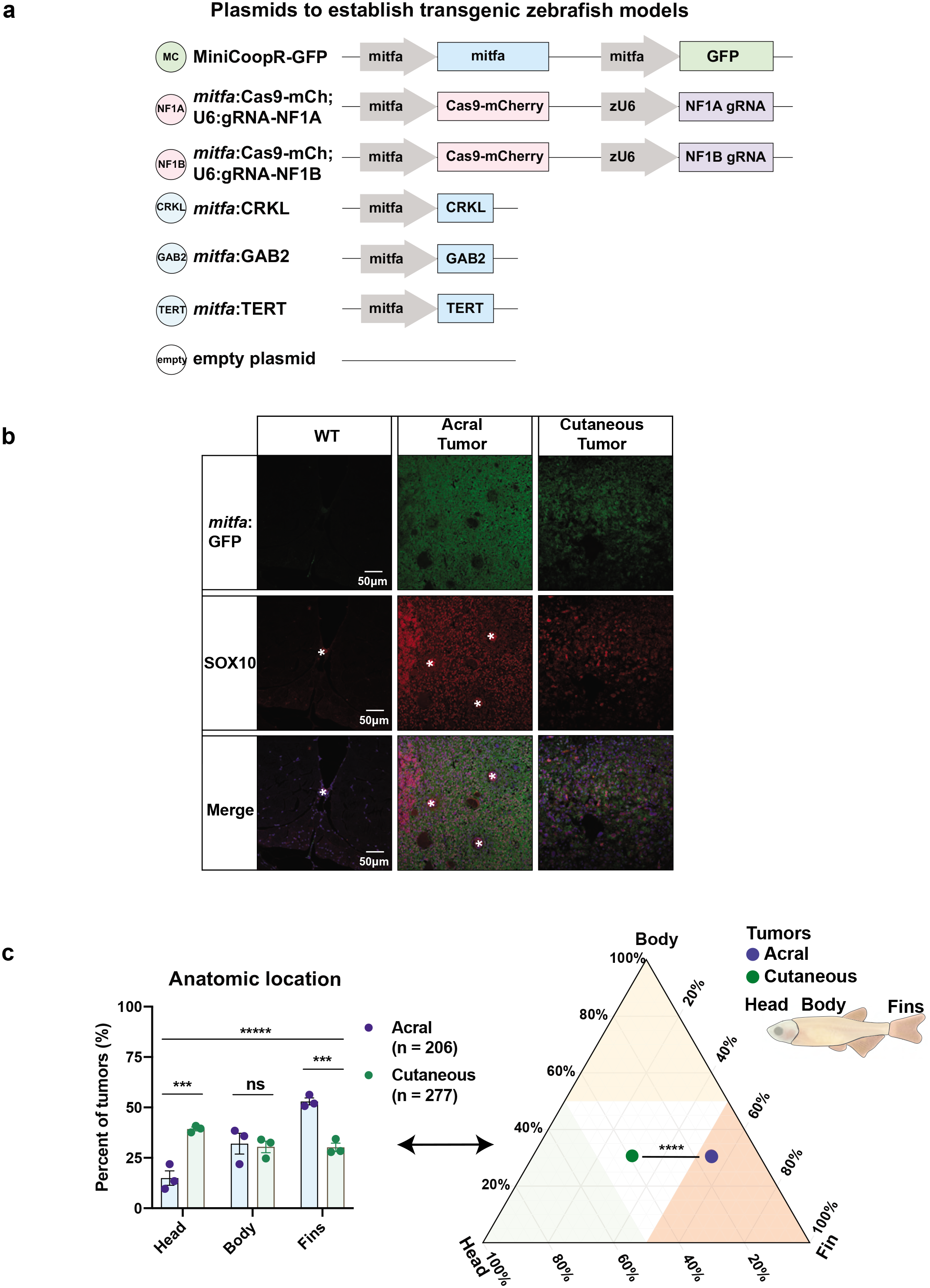
A transgenic zebrafish model of acral melanoma. Related to Fig. 1. (A) Plasmids used to create the WT melanocyte model, acral melanoma model, and cutaneous melanoma models. (b) Immunofluorescence on transverse sections of WT melanocyte model, tumor-bearing acral melanoma model, and tumor-bearing cutaneous melanoma model for GFP and SOX10. GFP labels all tumor cells and SOX10 is used as a melanocyte lineage marker. Asterisks indicate blood vessels with autofluorescence. (c) Histogram representation of the ternary plot portrayed in Fig. 1e showing the percentage of tumors forming in the head, body, or fins of acral fish and cutaneous fish melanoma models. Data represents three biological replicates, which range from n=43 to n=141 fish per replicate. See Supplemental Table 3 showing the exact number of fish and corresponding percentages for each replicate. To compare overall anatomic distribution between the two genotypes, a Chi-squared test was performed. To compare the frequency of tumors at each anatomic location, a student’s two-sided t-test was performed. Error bars = SEM. Ternary plot represents the same data presented in the histogram. * p-value < 0.05, ** p-value < 0.01, *** p-value < 0.001, **** p-value < 0.0001.

**Extended Data Fig. 3:**
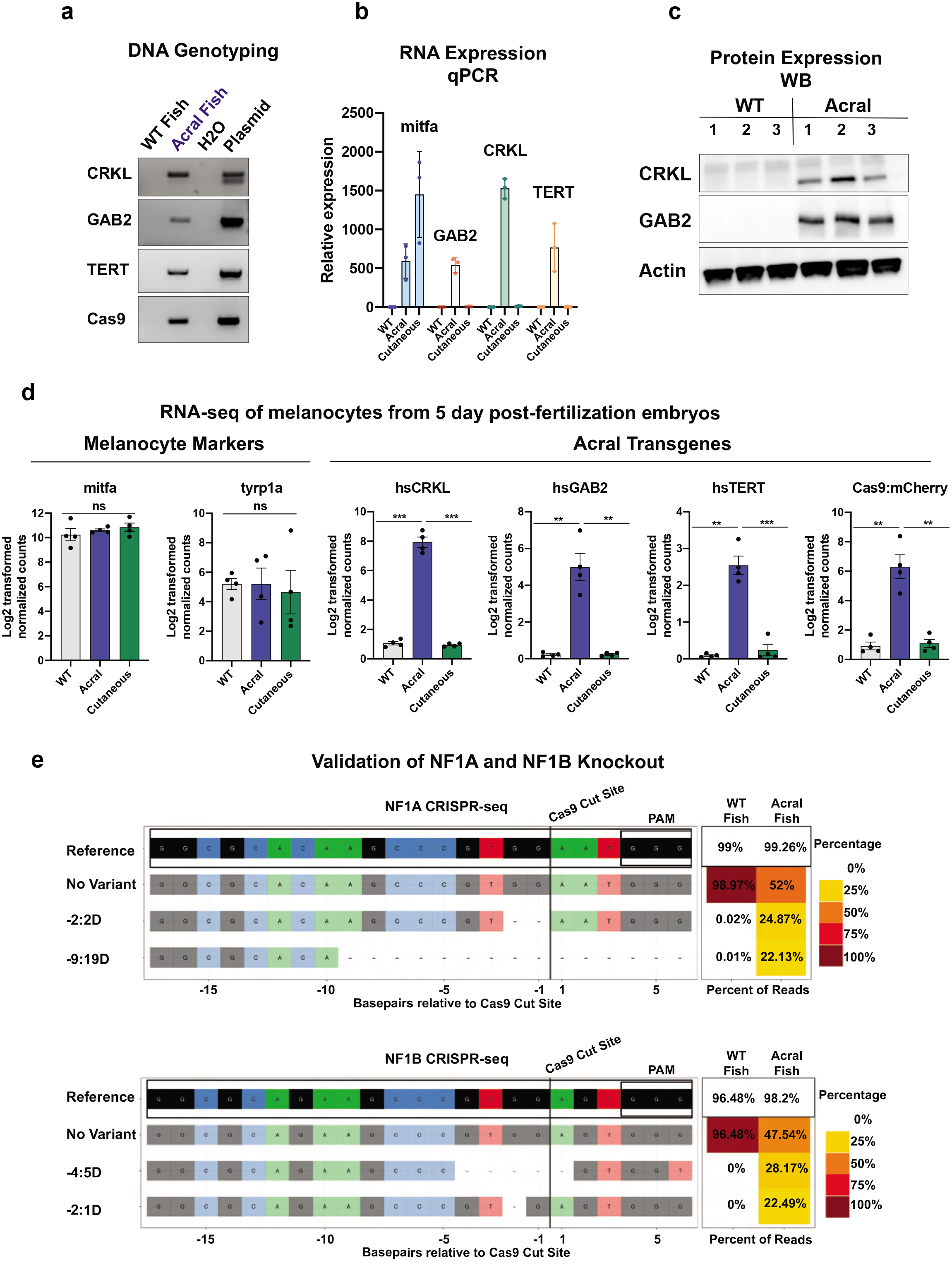
Acral melanoma model demonstrates overexpression and knockout of multiple driver genes. Related to Fig. 1. (a) Genotyping PCR to confirm integration of plasmids into zebrafish genome. (b) qPCR to validate RNA expression of transgenes. (c) Western blot was performed on WT and acral melanoma fish for human CRKL and GAB2 to validate transgene expression. Three biological replicates are shown. (d) RNA-seq on embryos at 5 days post-fertilization. Log normalized expression of melanocyte markers and acral transgenes are indicated. P-values calculated using DESeq2. (e) CRISPR-seq was performed on the predicted sgRNA cut locus of zebrafish *nf1a* and *nf1b* to sensitively detect Cas9-mediated editing. Reference genome as well as the two most commonly altered reads are shown. The right panel is a heatmap with the frequency of reference and edited sequences in WT melanocyte and acral melanoma models. Variants displayed for both *nf1a* and *nf1b* are frameshift mutations in exon 1 leading to a predicted loss of function. * p-value < 0.05, ** p-value < 0.01, *** p-value < 0.001, **** p-value < 0.0001.

**Extended Data Fig. 4:**
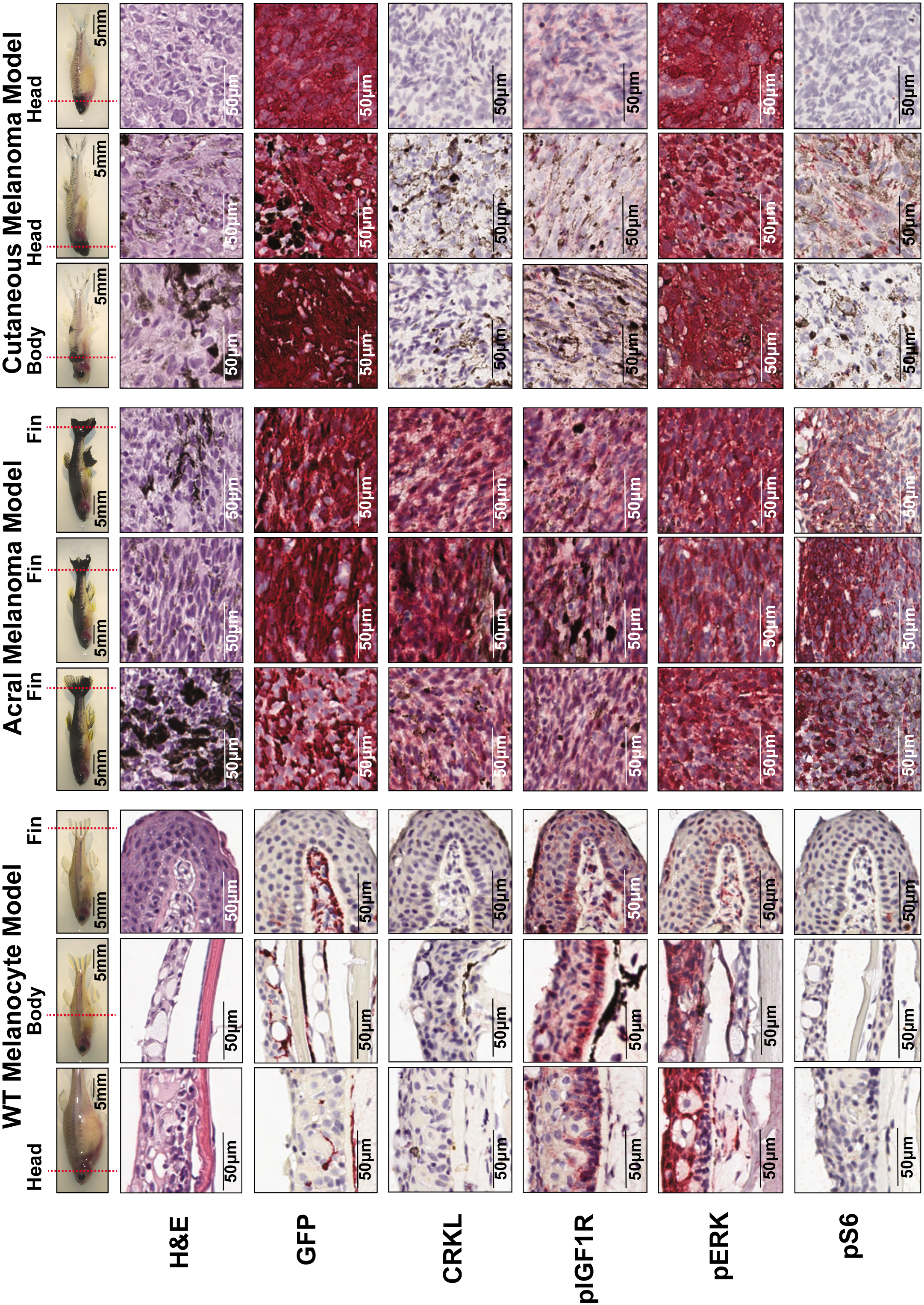
Histological profiling of zebrafish model of acral melanoma. Related to Fig. 1. n=3 WT, acral, and cutaneous melanoma transgenic fish were used for histological profiling. Images of the fish used for histology are shown on top with a red line indicating anatomic region used for sectioning and profiling. For WT fish, head, body, and fin regions were used as a negative control. For acral fish, profiled tumors were located on the fins. For cutaneous fish, profiled tumors were located on the body and head. The markers used include H&E, GFP (marking melanocytes in all 3 models), CRKL (expressed only in the acral model), pIGF1R, pERK, and pS6. See methods for details regarding antibody, concentration, and staining procedure.

**Extended Data Fig. 5:**
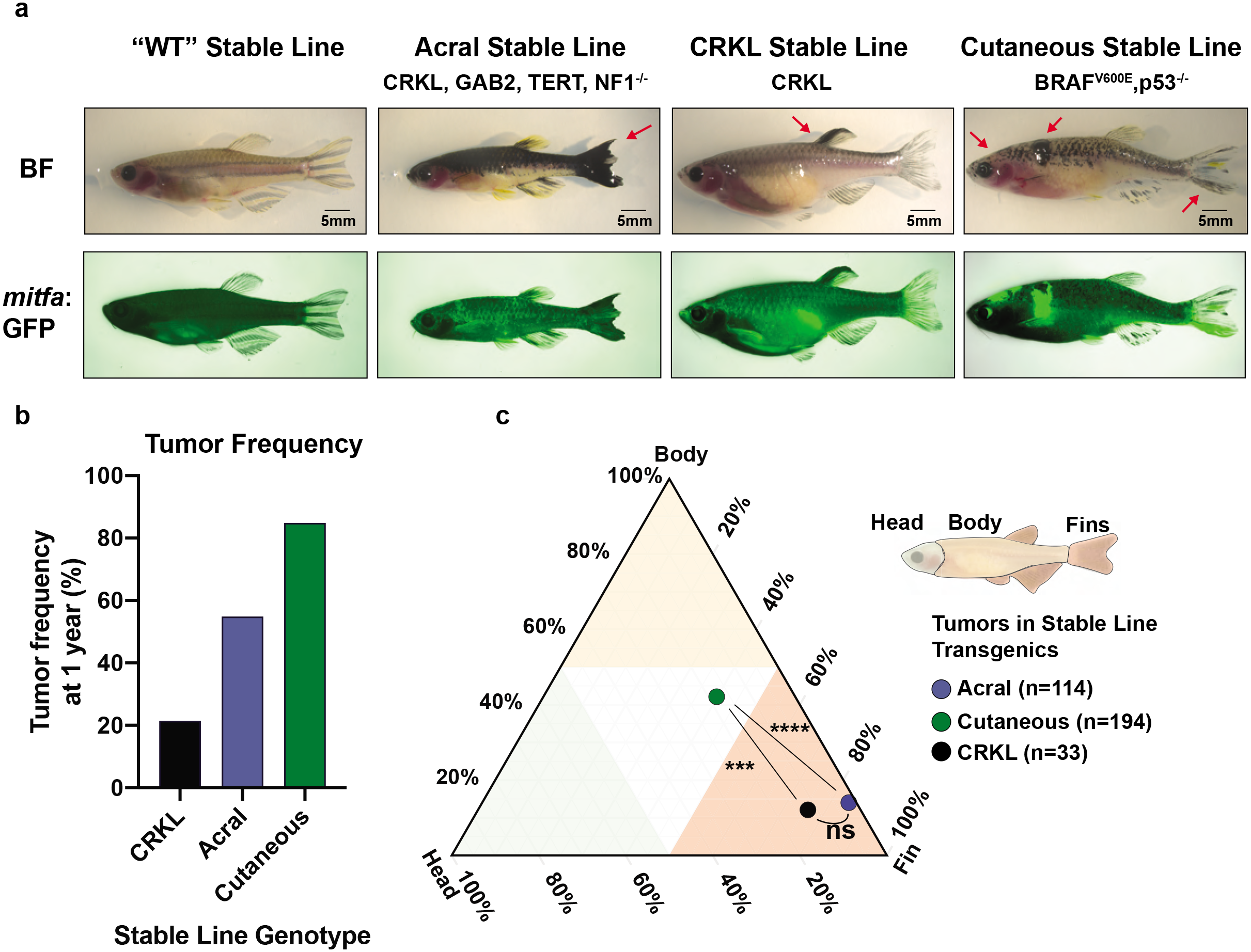
Stable (germline) transgenic acral lines show enhanced anatomic specificity. Related to Fig. 1. (a) F0 transgenic fish shown in Fig 1d and Fig 2b were outcrossed to generate stable germline transgenics. Representative images of each transgenic line are shown with arrows to indicate the location of tumors. (b) Tumor frequency of each stable line at 1 year post fertilization. (c) Ternary plot showing the anatomic distribution of tumors for each stable transgenic line. P-values generated by Chi-squared test. See Supplemental Table 3 for a full list of fish and tumor numbers across all replicates and experimental conditions.

**Extended Data Fig. 6:**
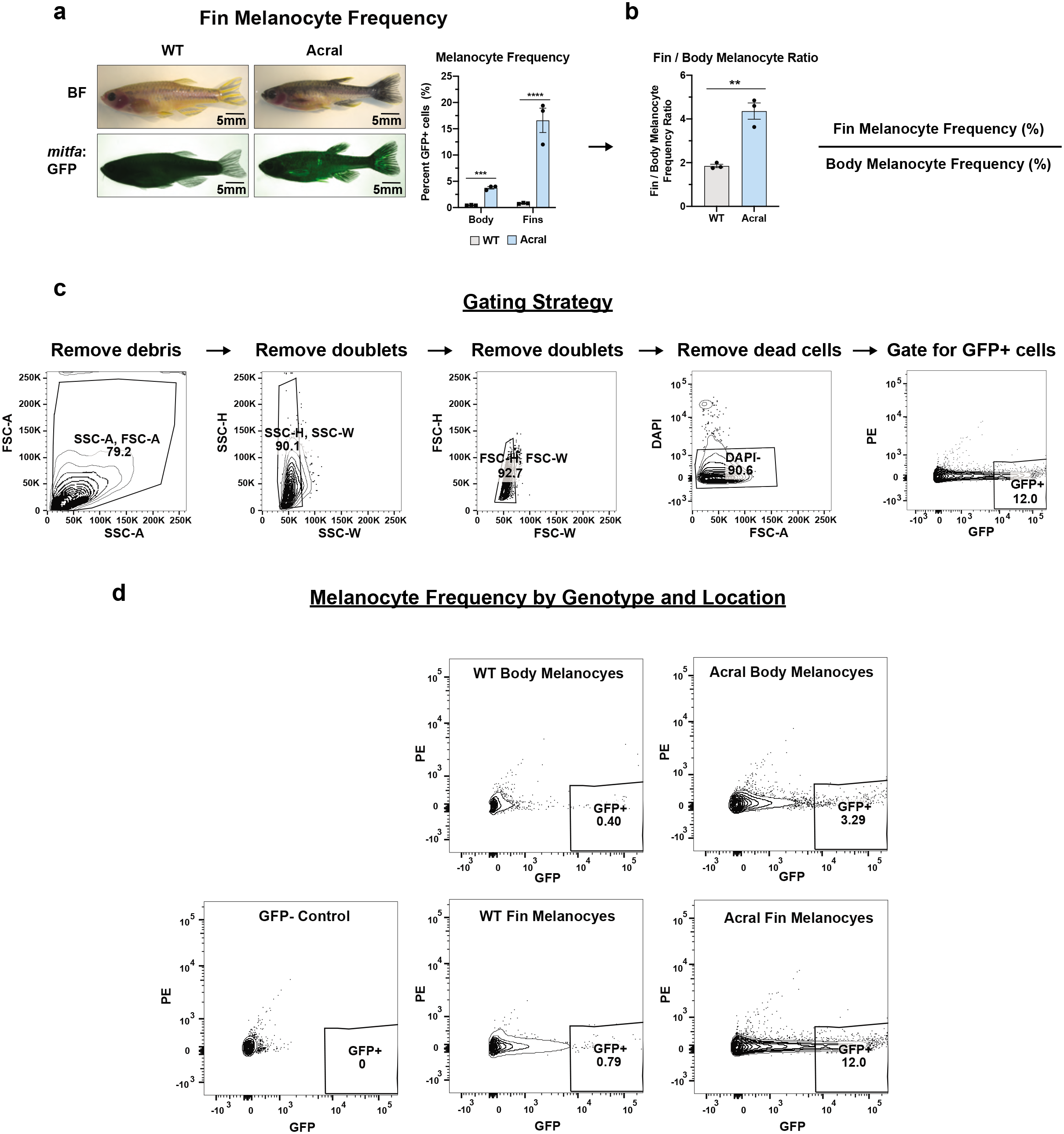
Acral melanoma model demonstrates melanocyte enrichment in the fins. Related to Fig. 1. (a) Adult WT melanocyte and acral melanoma models were dissected to isolate body skin and fins and then analyzed via flow cytometry. The images show representative differences in pigment patterning between the two models. The histogram shows percentage of total cells that were GFP+ grouped by anatomic location and genotype. Data represents three biological replicates. Each replicate represents the pooling of n=2 male and n=2 female fish. P-values generated by student’s two-sided t-test. Error bars = SEM. (b) The ratio of fin melanocytes to body melanocytes was compared between the two models using the data from (a). Data represents three biological replicates. P-values generated by student’s two-sided t-test. Error bars = SEM. (c) Gating strategy of the flow cytometry data. (d) Representative contour plots of melanocyte frequency by genotype and location. * p-value < 0.05, ** p-value < 0.01, *** p-value < 0.001, **** p-value < 0.0001.

**Extended Data Fig. 7:**
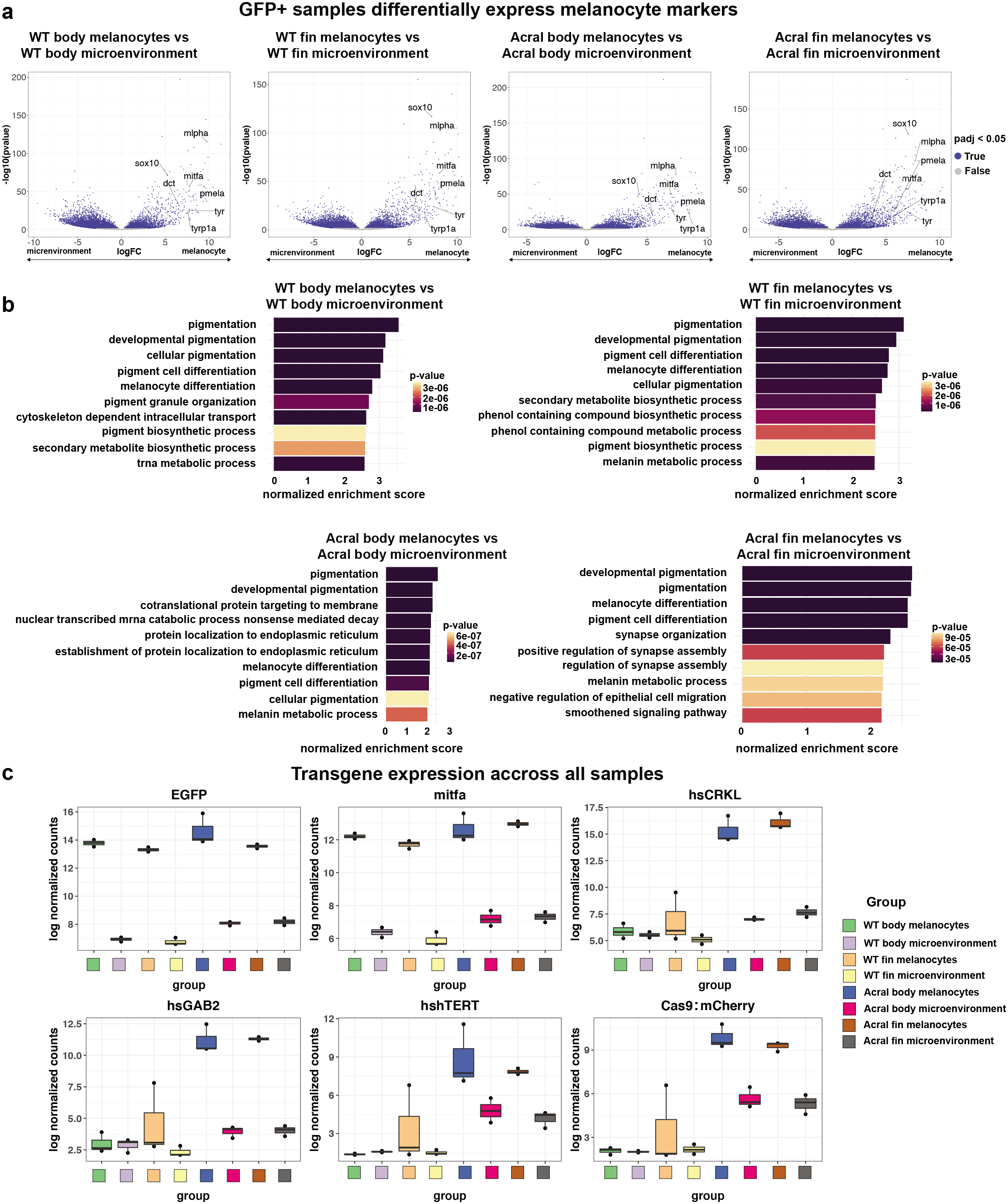
Validation of zebrafish melanocyte RNA-seq. Related to Fig. 3. (a) Volcano plots comparing melanocytes (GFP+ sample) to their microenvironment (GFP-sample) across locations and transgenic models. Melanocyte markers are labeled. Genes with FDR-adjusted p-value < 0.05 indicated in blue. P-values calculated using DESeq2. (b) GSEA showing the list of the top pathways enriched in melanocytes (GFP+ sample) compared to their microenvironment (GFP-sample). Colors indicate p-value adjusted for FDR = 0.05. (c) Log normalized counts for the expression of transgenes across all samples. EGFP and *mitfa* expression are high in all melanocyte samples and CRKL, GAB2, TERT, and Cas9-mCherry expression is high only in acral melanoma model melanocytes. Error bars = SEM.

**Extended Data Fig. 8:**
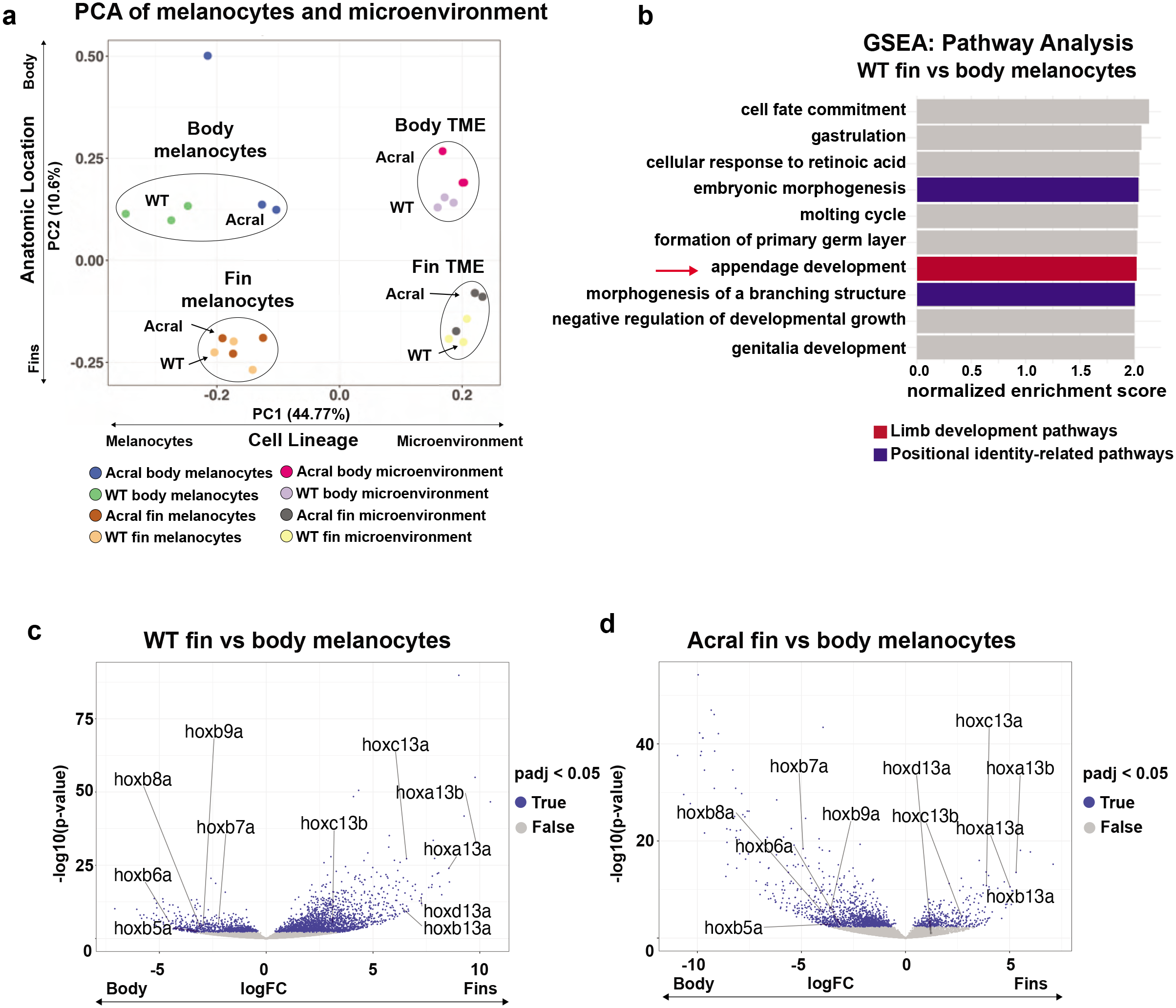
Fin melanocytes have a limb positional identity. Related to Fig. 3. (a) Principal component analysis (PCA) for all samples showing principal components 1 (PC1) and 2 (PC2). The percent of transcriptional variation captured by each principal component is indicated. (b) GSEA pathway analysis comparing fin vs body melanocytes from the WT melanocyte model listing the top enriched pathways in fin melanocytes. Limb development and positional identity-related pathways are highlighted. See Supplemental Table 4 for a full list of pathways. (c-d) Volcano plots comparing fin melanocytes vs body melanocytes from the (c) WT melanocyte model and (d) acral melanoma model. Genes with FDR-adjusted p-value < 0.05 indicated in blue. P-values calculated using DESeq2.

**Extended Data Fig. 9:**
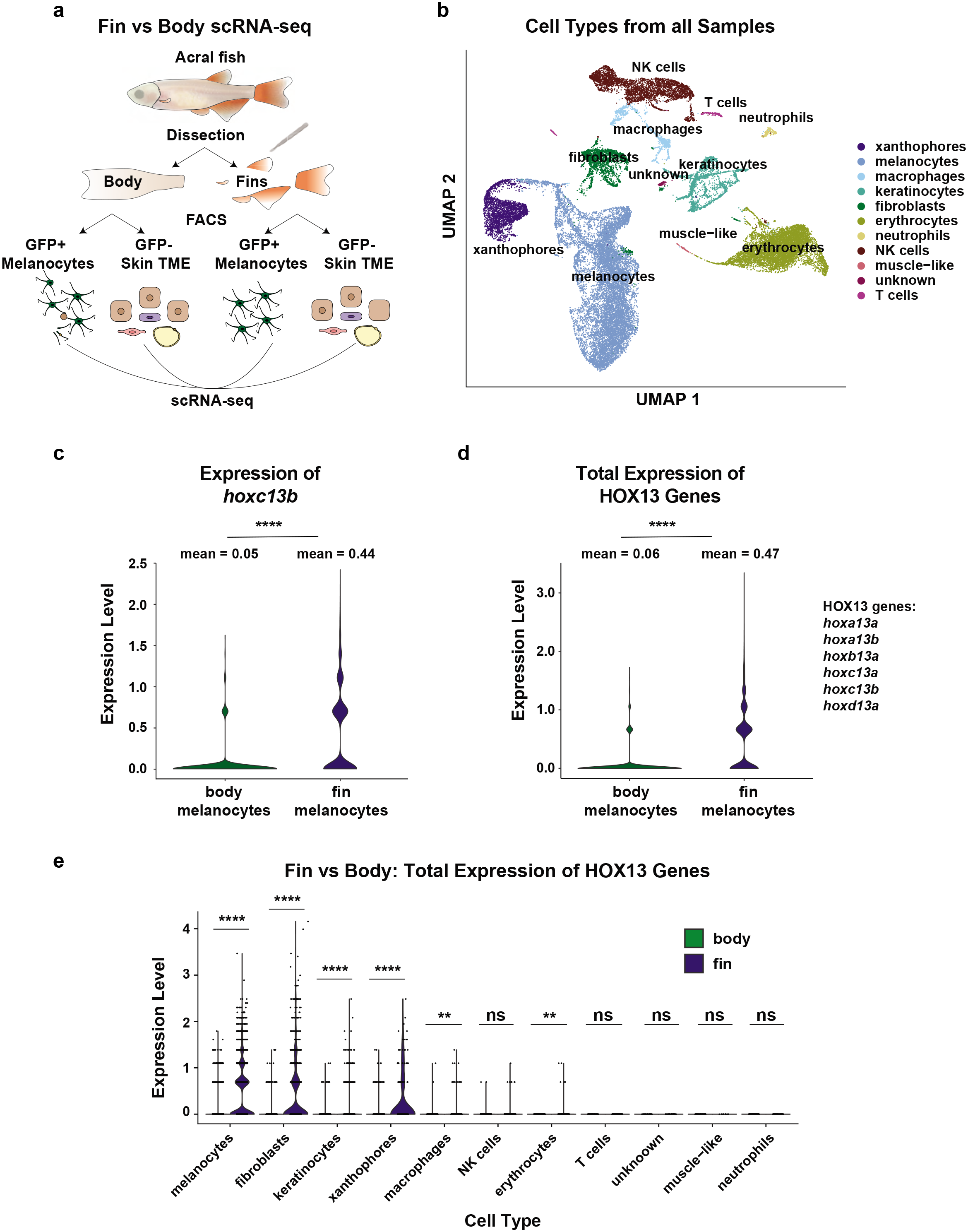
Zebrafish fin melanocytes express higher levels of HOX13 genes. Related to Fig. 3. (a) Schematic illustrating the fin versus body scRNA-sequencing experiment. Similar to the bulk RNA-seq experiment in Fig. 3a, body skin and fins were isolated from acral melanoma fish by dissection and then FACS sorted for GFP+ melanocytes and GFP-microenvironmental cells. This generated n=4 samples (body melanocytes, body TME, fin melanocytes, fin TME) that each underwent scRNA-seq. (b) UMAP pooled for all 4 samples highlighting the various cell types captured by scRNA-seq. (c) Violin plot comparing *hoxc13b* expression in body and fin melanocytes. Data represents log normalized counts and was compared by Wilcoxon rank sum test. (d) Violin plot comparing total expression of all HOX13 genes (*hoxa13a*, *hoxa13b*, *hoxb13a*, *hoxc13a*, *hoxc13b*, *hoxd13a*). To avoid single-cell drop out of more lowly expressed HOX13 genes, log normalized counts were summed together to calculate total expression per cell. Data was analyzed by Wilcoxon rank sum test. (e) Violin plot comparing the total expression of all HOX13 genes across all fin and body cell types. Data was analyzed by Wilcoxon rank sum test. * p-value < 0.05, ** p-value < 0.01, *** p-value < 0.001, **** p-value < 0.0001.

**Extended Data Fig. 10:**
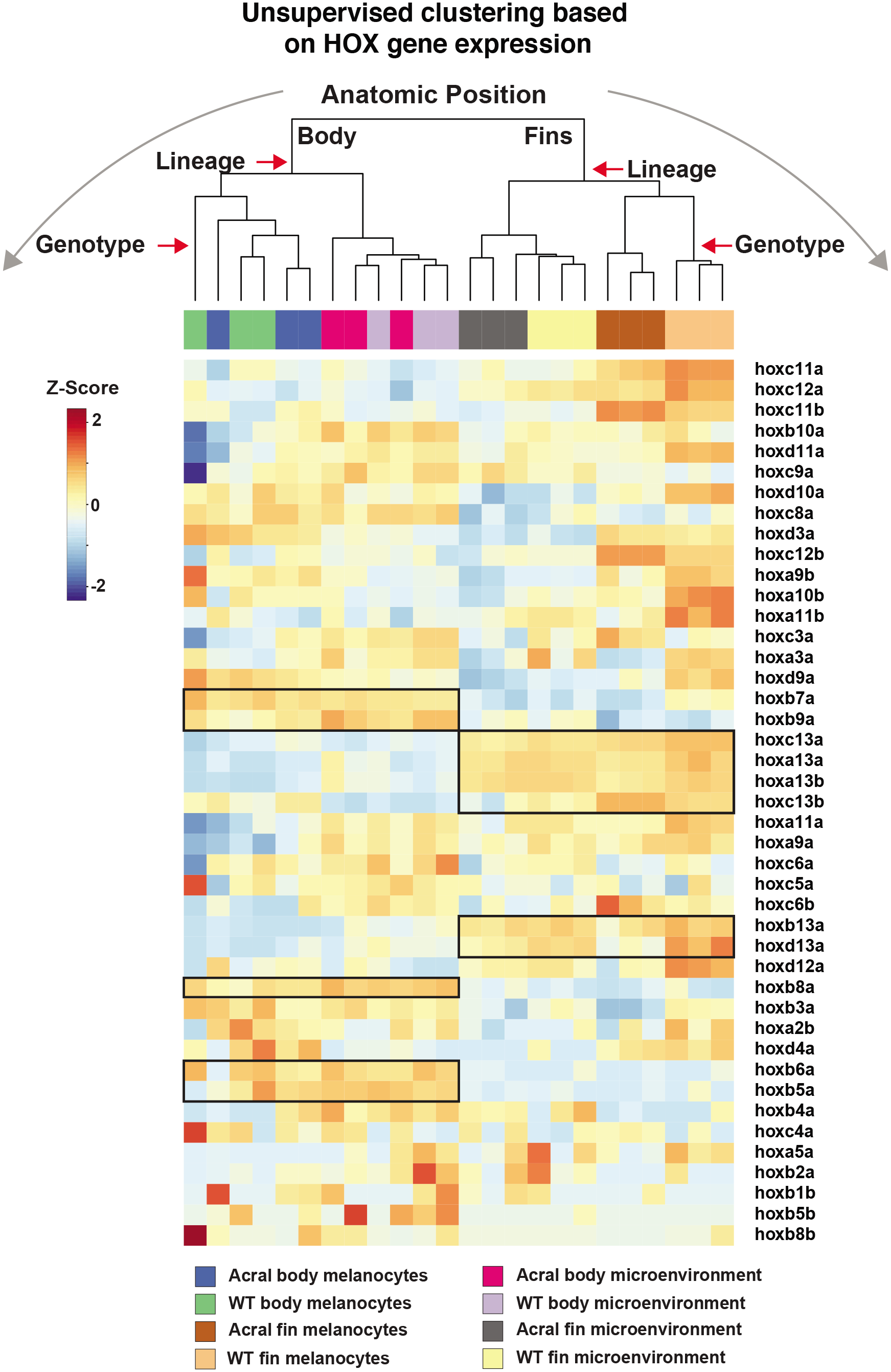
Zebrafish melanoma has a positional identity defined by *hox* genes. Related to Fig. 3. Zebrafish *hox* genes detected by bulk RNA-seq were used to perform unsupervised clustering and visualized with a heatmap. Samples clustered by anatomic location, then lineage, and then genotype in that order, indicating the *hox* genes predominantly associate with anatomic position. Particular *hox* genes that are differentially expressed across all body and fin samples are indicated with a black box.

**Extended Data Fig. 11:**
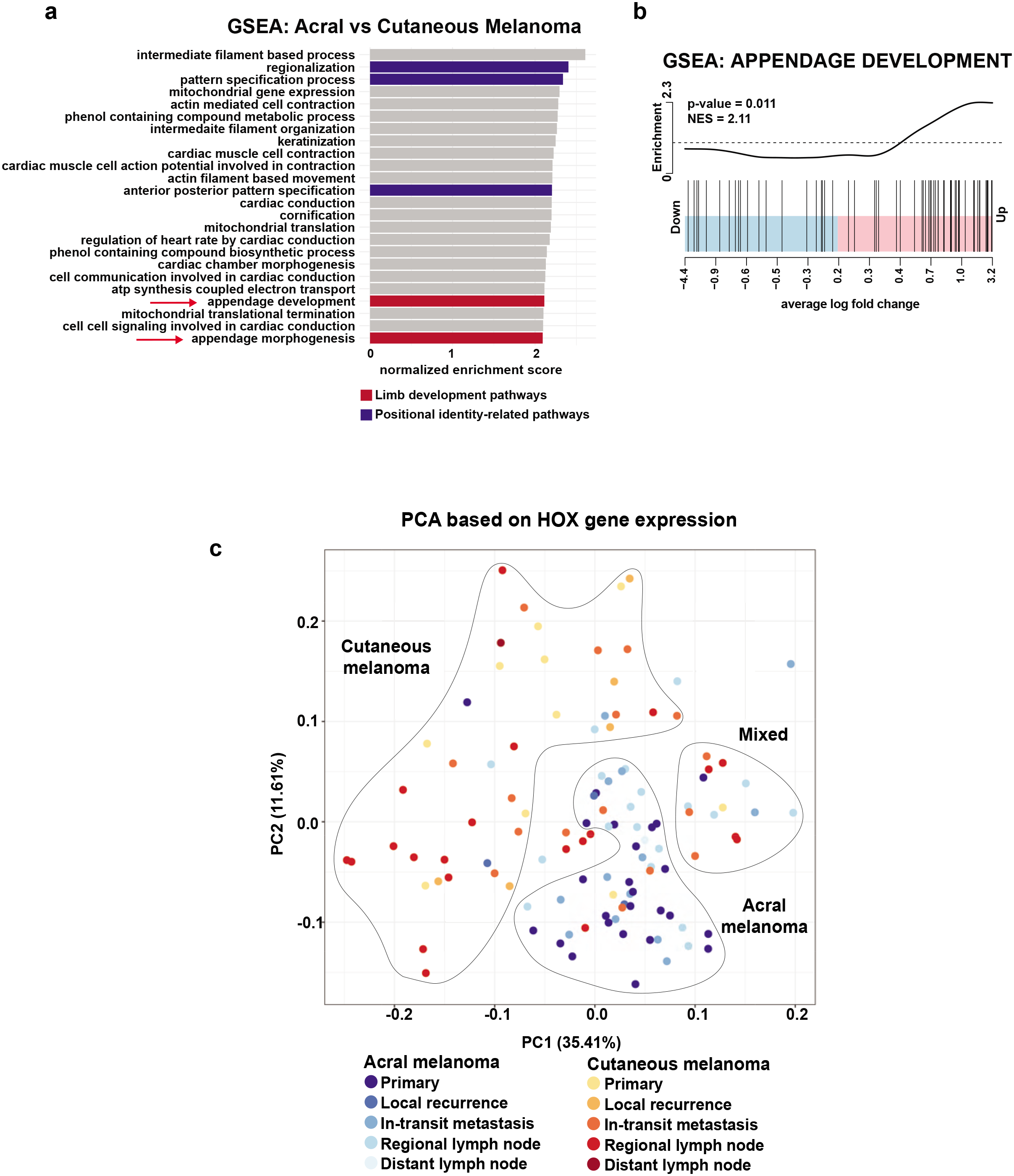
Human melanoma has a positional identity defined by HOX genes. Related to Fig. 4. (a) GSEA pathway analysis for top enriched pathways in human acral melanoma and is controlled for disease stage. Limb development pathways and positional identity-related pathways are highlighted. See Supplemental Table 5 for a full list of pathways. (b) GSEA barcode plot showing enrichment for the GO: Appendage Development pathway in acral melanoma. NES and FDR are indicated. (c) PCA of all samples based on the expression of just HOX genes. Color indicates the combination of melanoma subtype and disease stage.

**Extended Data Fig. 12:**
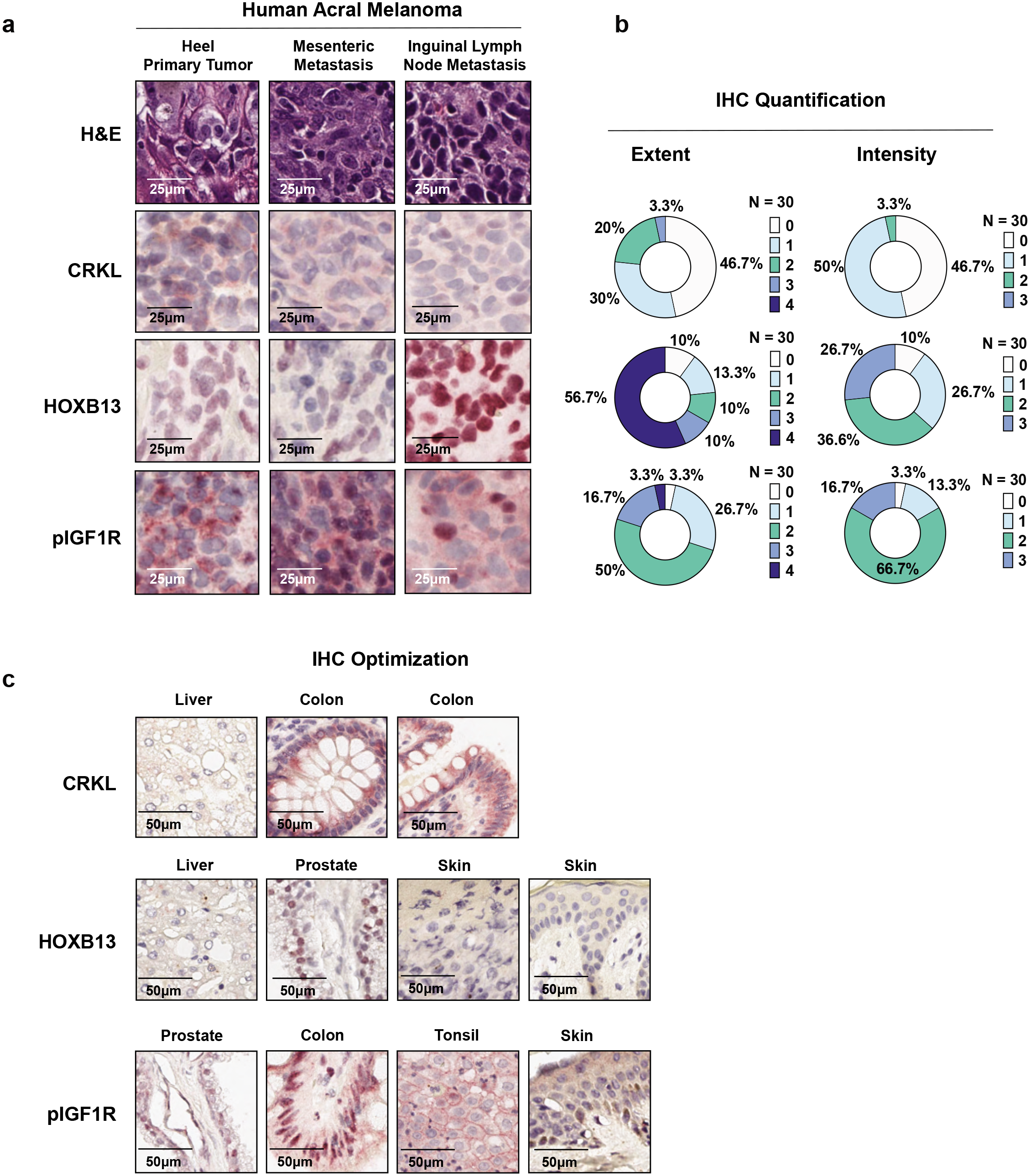
Histological profiling of human acral melanoma. Related to Fig. 4. (a) An acral melanoma tissue microarray of n=32 samples from Sloan Kettering was used for staining for H&E, CRKL, HOXB13, and pIGF1R. (b) Extent and intensity of each sample was independently scored by a Sloan Kettering dermatopathologist and is represented in a pie chart. 2 samples were removed from the analysis due to heavy pigmentation, resulting in n=30 samples total used for IHC quantification analysis. Extent scores range 0-4 and intensity scores range 0-3. (c) CRKL, HOXB13, and pIGF1R antibodies were optimized on human tissues chosen for their expression characterized by the Human Protein Atlas. CRKL is a ubiquitously expressed protein with relatively high expression in the colonic epithelia and lower expression in liver. HOXB13 is expressed in prostate epithelia with low expression in liver and skin (anatomic source unknown). pIGF1R demonstrates higher levels in prostate epithelia, colon epithelia, tonsil, and lower levels in skin. See methods for details regarding antibody, concentration, and staining procedure.

**Extended Data Fig. 13:**
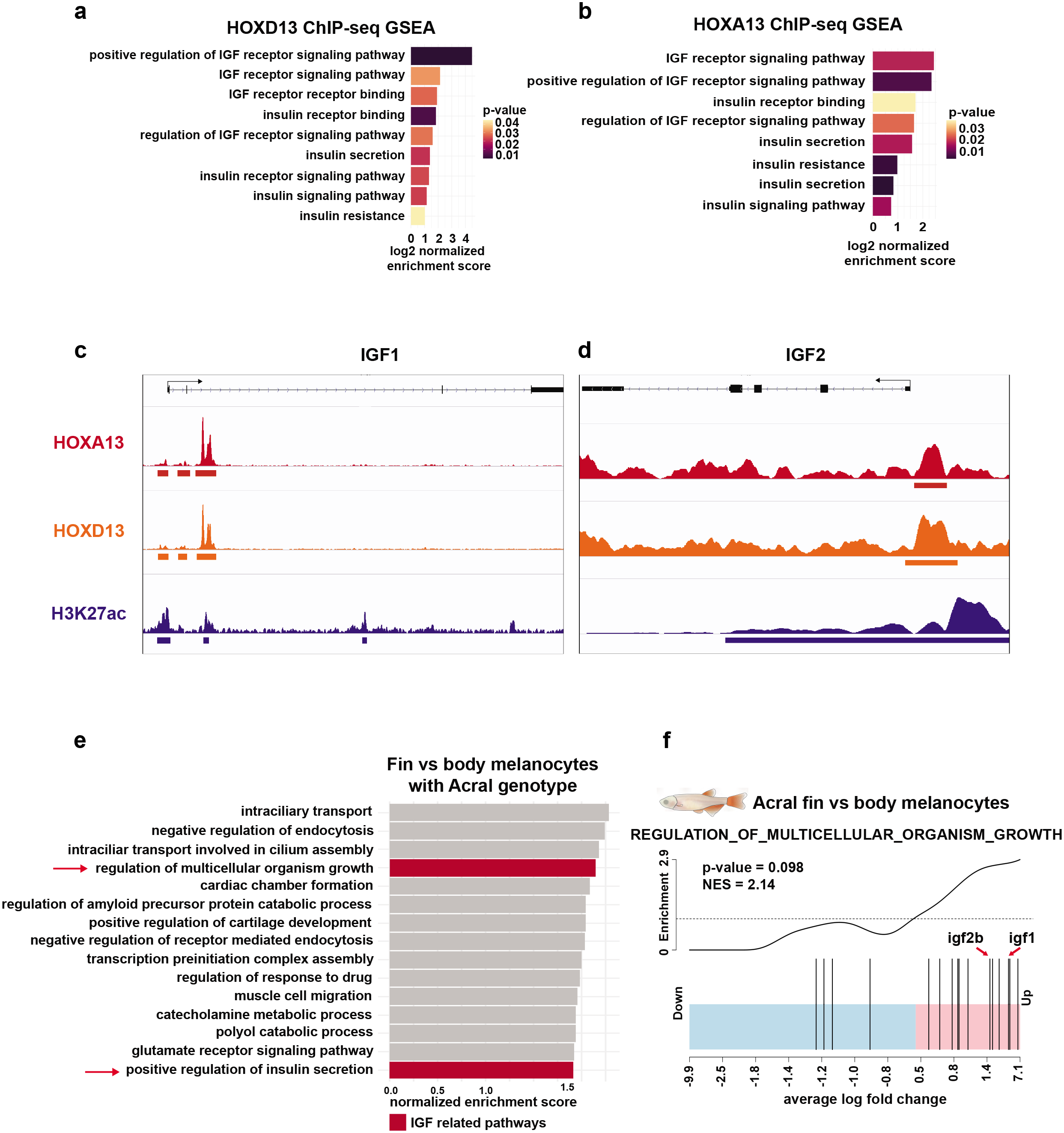
HOX13 regulates insulin/IGF signaling. Related to Fig. 5. (a-d) Analysis of HOXA13, HOXD13, H3K27ac ChIP-seq data analyzed from Sheth et al., 2016^43^ performed on developing limb buds from E11.5 mouse embryos. (a-b) Histogram showing the significantly enriched insulin/IGF pathways regulated by HOXA13 and HOXD13. NES and p-value are indicated. (c-d) Integrated genome browser tracks for HOXA13 and HOXD13 ChIP-seq binding and H3K27 acetylation near transcription start site of IGF1 and IGF2. A full list of pathways can be found in Supplemental Table 6. (e) GSEA pathway analysis for the top pathways enriched in fin melanocytes versus body melanocytes from the zebrafish acral melanoma model. IGF-related pathways are indicated in red. A full list of pathways can be found in Supplemental Table 4. (f) GSEA barcode plot comparing acral model fin vs body melanocytes showing enrichment of genes in the GO: Regulation of Multicellular Organisms Growth pathway. NES and FDR are indicated.

**Extended Data Fig. 14:**
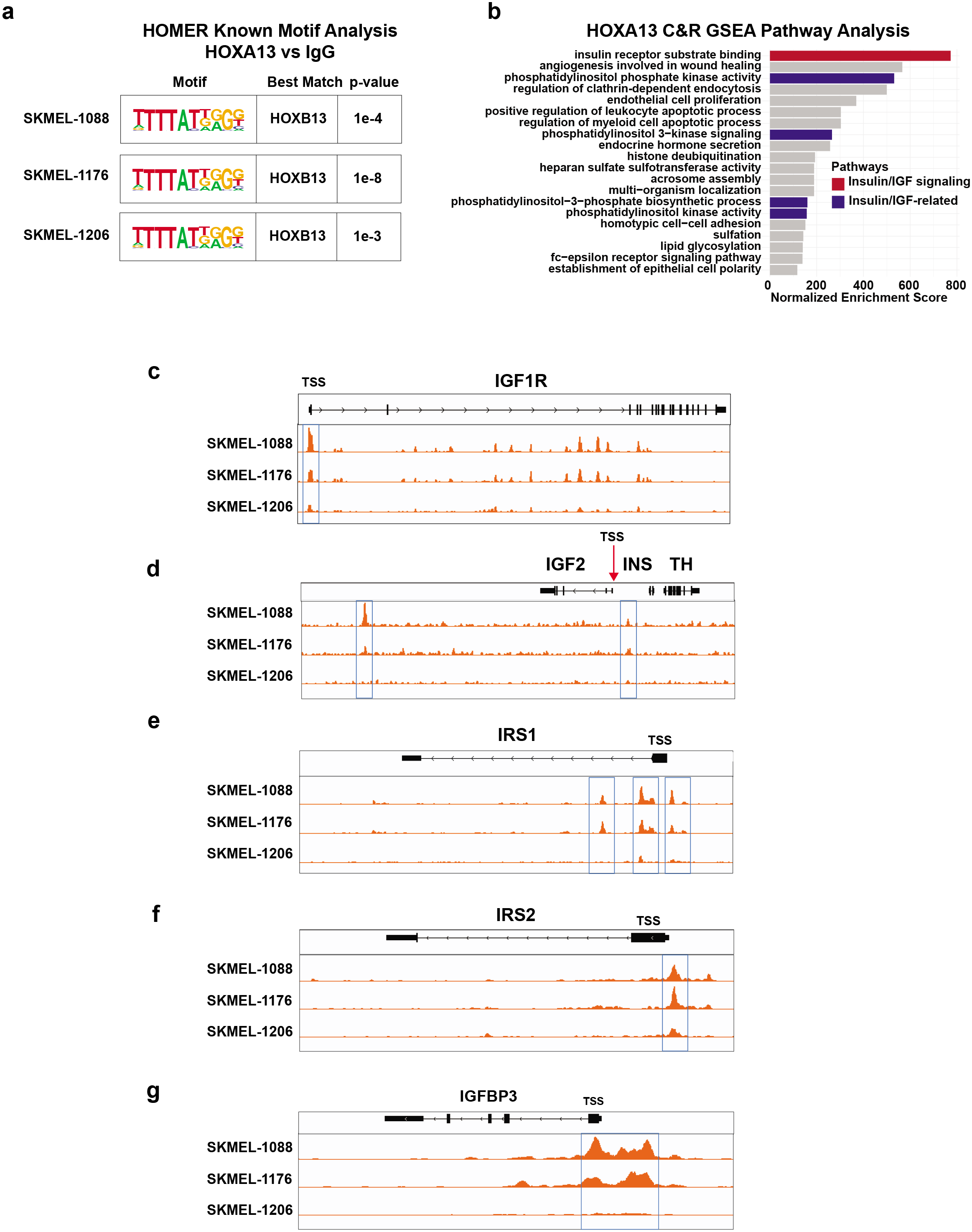
HOX13 regulates insulin/IGF signaling in human melanoma cells. Related to Fig. 5. (a-g) Cut & Run was performed on HOX13 expressing human melanoma cell lines, SKMEL-1088, SKMEL-1176, SKMEL-1206, using an antibody against HOXA13 and IgG as a negative control. (a) HOMER known motif analysis comparing HOXA13 vs IgG peaks identified significant enrichment for HOXB13 in all 3 melanoma cell lines. (b) GSEA pathway analysis using Cistrome-GO was performed, and the top enriched pathways are listed highlighting pathways related to insulin/IGF signaling. Insulin/IGF signaling pathways were identified using the search terms “insulin” and “IGF.” Insulin/IGF-related pathways were identified based on literature and specify pathways operating directly downstream of insulin/IGF signaling. A full list of pathways can be found in Supplemental Table 7. (c-g) IGV plot showing peaks along IGF1R, IGF2, IRS1, IRS2, and IGFBP3 for all 3 cell lines. The transcription start site (TSS) is indicated.

**Extended Data Fig. 15:**
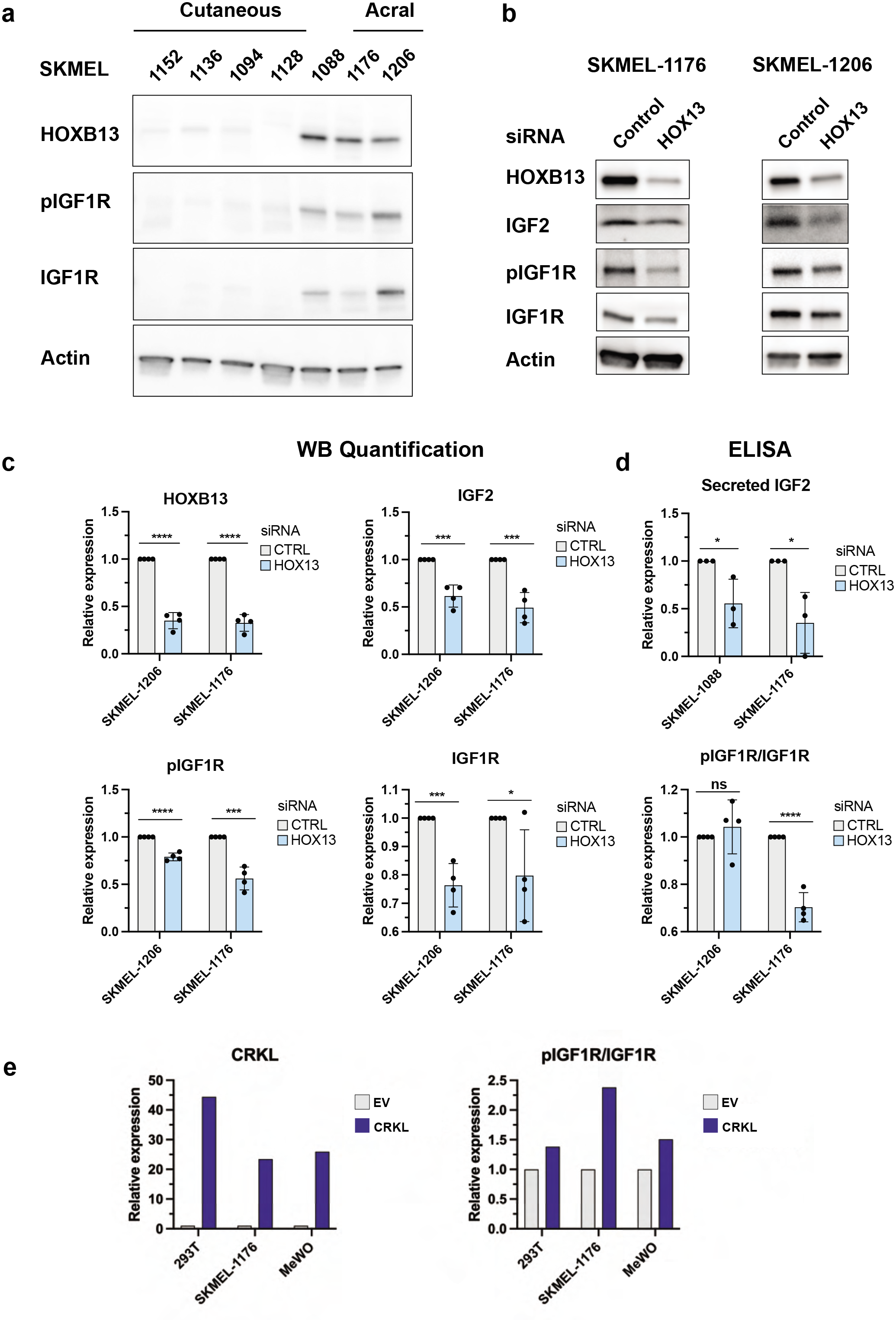
HOX13 and CRKL increase IGF signaling in human melanoma cells. Related to Fig. 5. (a) Patient derived melanoma cell lines were analyzed for HOXB13, pIGF1R, and total IGF1R expression by Western blot. Actin was used as a loading control. Cell lines derived from acral and cutaneous tumors are labelled. SKMEL-1088 is a melanoma cell line of unknown anatomic origin. (b) SKMEL-1176 and SKMEL-1206 acral melanoma cell lines were transfected with siRNAs targeting all 4 human HOX13 genes (HOXA13, HOXB13, HOXC13, HOXD13) or a non-targeting control. Cells were grown in standard media conditions. 96 hours post-transfection, cells were collected and analyzed by Western blot. HOXB13 was used to validate siRNA knockdown and then also probed for IGF2 ligand, pIGF1R, and IGF1R. Actin was used as a loading control. Blots are representative of n=4 biological replicates. (c) Western blot quantification of the n=4 biological replicates from (b). Data was analyzed using a two-sided student’s t-test. (d) SKMEL-1088 and SKMEL-1176 were transfected with siRNA’s targeting all 4 human HOX13 genes (HOXA13, HOXB13, HOXC13, HOXD13) or a non-targeting control. 48 hours post-transfection cells were changed to serum-free media. 96 hours post-transfection, conditioned media was collected and used for ELISA to measure secreted IGF2. N = 3 biological replicates were performed. Data was analyzed using a two-sided student’s t-test. (e) Quantification of Western blots shown in Fig. 5f. 293T, SKMEL-1176, and MeWo human cell lines were transduced with CRKL overexpression vector or an empty vector control. Cells underwent blasticidin selection and then were collected for Western blot analysis for CRKL, pIGF1R, and IGF1R. Actin was used as a loading control. * p-value < 0.05, ** p-value < 0.01, *** p-value < 0.001, **** p-value < 0.0001.

**Extended Data Fig. 16:**
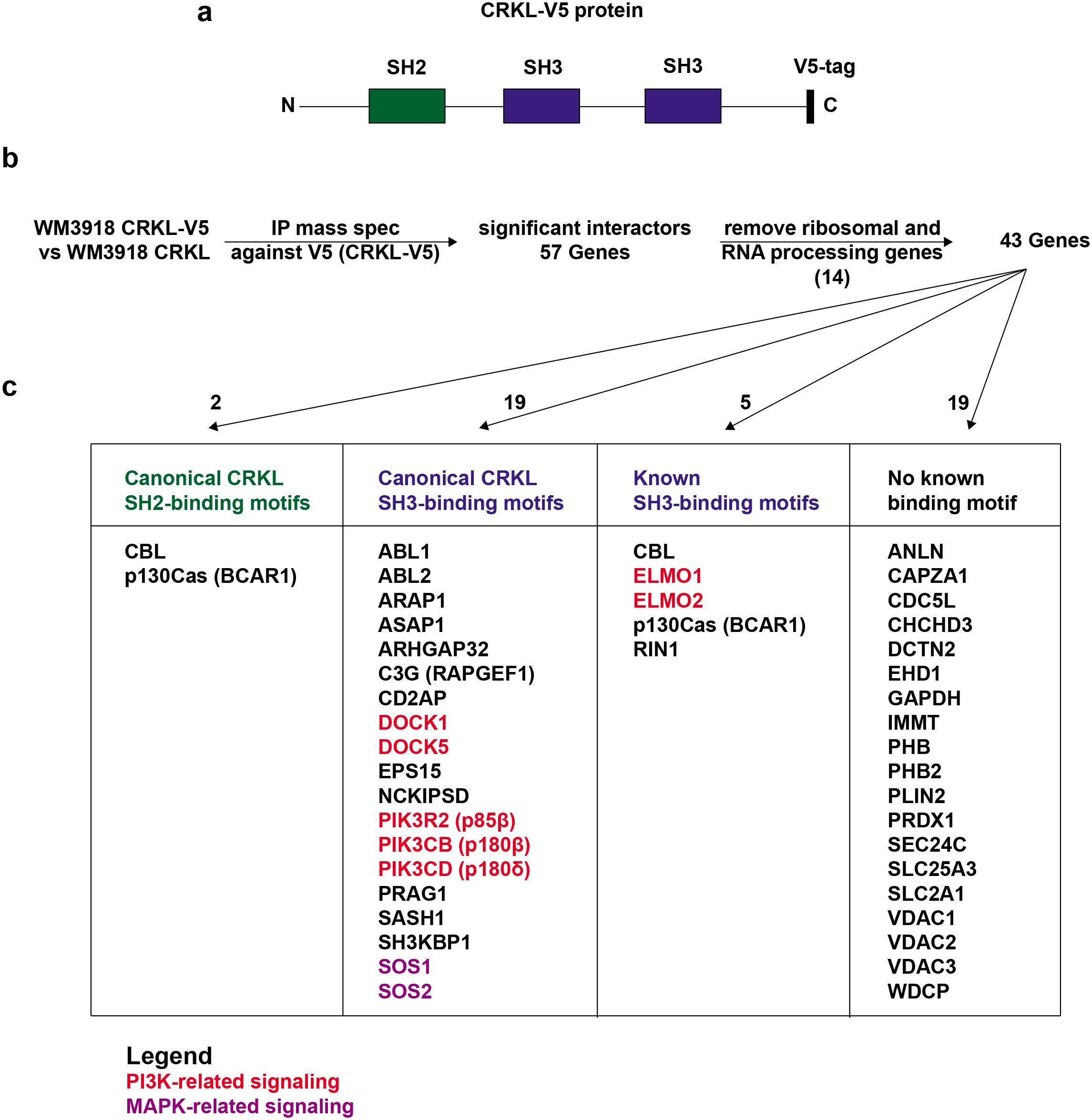
CRKL interaction demonstrates direct binding to PI3K. Related to Fig. 5. (a) Schematic representation of CRKL with a V5 tag. CRKL is composed of a SH2 (Src homology) domain that bind proteins with phospho-tyrosines and two SH3 domains that bind to proline-rich proteins. (b) Description of the experimental design and analysis. Briefly, human melanoma cell line WM3918 overexpressing CRKL with a V5 tag was compared to CRKL overexpressing cells without the V5 tag. N = 4 replicates were used for each condition. Lysates were immunoprecipitated using a V5 antibody and then underwent mass spectrometry, yielding 56 significant interactors as defined by logFC > 2 and corrected p-value < 0.05. Significant hits were filtered for ribosomal and RNA processing genes, leaving 43 significant interactors with CRKL. (c) Significant interactors with CRKL were organized based on whether they contain a canonical CRKL SH2-binding domain (pY-x-x-P), canonical CRKL SH3 binding motif (Ψ-P-Ψ-L/V/P/A/I-P-Ψ-K), known SH3 binding motif (proline-rich sequence), or no identified binding motif. Genes are color coded by their contribution to PI3K and MAPK signaling. Raw data can be found in Supplemental Table 8.

**Extended Data Fig. 17:**
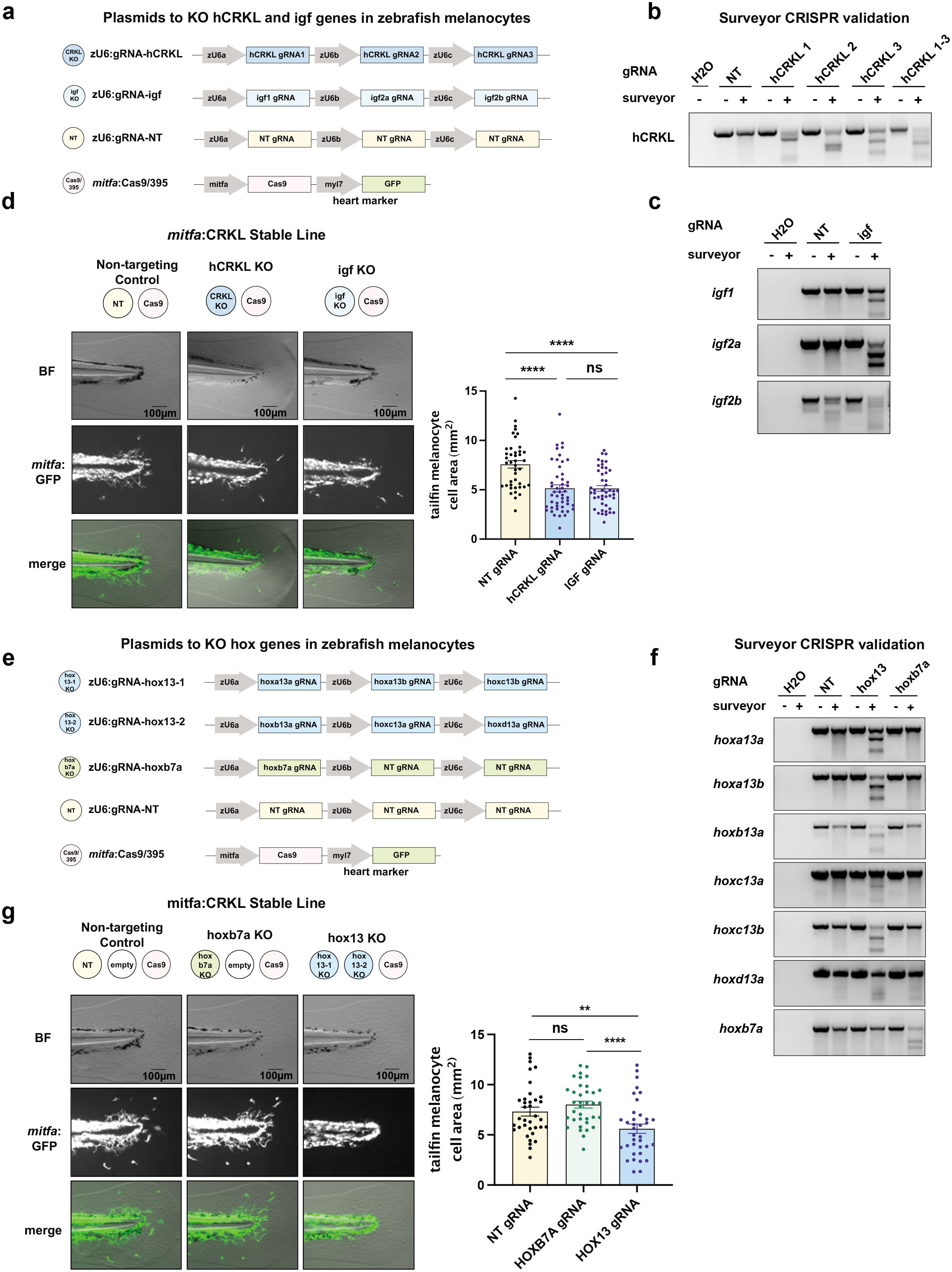
CRKL-mediated melanocyte tailfin expansion depends on a HOX13/IGF positional program. Related to Fig. 5. (a) Plasmids used to knockout the human CRKL transgene and zebrafish IGF ligands (*igf1, igf2a*, *igf2b*) in a mosaic manner in CRKL stable line zebrafish. *mitfa*:Cas9 ensures melanocyte specificity in CRISPR editing. This plasmid also contains a fluorescent heart marker *myl17*:GFP, which was used to determine which fish had successful plasmid integration to be used in further downstream image analysis. (b) Surveyor validation demonstrating targeted editing of the human CRKL transgene using 3 different sgRNAs. (c) Surveyor validation demonstrating targeted editing of all zebrafish IGF ligands using 3 different sgRNAs. (d) *mitfa*:CRKL stable line fish were injected with indicated plasmids at the one-cell stage. At 3-days post-fertilization, embryos with GFP+ hearts were selected, and their tailfins were imaged for melanocyte area. Representative images and quantification are provided. Data represents n=42 fish receiving a non-targeting (NT) sgRNA, n=49 receiving sgRNAs against the human CRKL transgene, and n=46 receiving sgRNAs against all zebrafish IGF genes pooled together over three biological replicates. Each point represents the tailfin melanocyte area of a different animal. Data was analyzed using a two-sided student’s t-test. (e) Plasmids used to knockout the zebrafish HOX13 genes (*hoxa13a*, *hoxa13b*, *hoxb13a*, *hoxc13a, hoxc13b*, *hoxd13a*) and orthologue of HOXB7, *hoxb7a*, in a mosaic manner in CRKL stable line zebrafish. *mitfa*:Cas9 ensures melanocyte specificity in CRISPR editing. This plasmid also contains a fluorescent heart marker *myl17*:GFP, which was used to determine which fish had successful plasmid integration to be used in further downstream image analysis. (f) Surveyor validation demonstrating targeted editing zebrafish HOX genes using 6 different sgRNAs. Note, HOX13-targeting plasmids only target HOX13 genes (not *hoxb7a*) and *hoxb7a*-targeting plasmid only targets *hoxb7a*. (g) *mitfa*:CRKL stable line fish were injected with indicated plasmids at the one-cell stage. At 3-days post-fertilization, embryos with GFP+ hearts were selected, and their tailfins were imaged for melanocyte area. Representative images and quantification are provided. Data represents n=36 fish receiving a non-targeting (NT) sgRNA, n=36 receiving a sgRNA against *hoxb7a*, and n=36 receiving sgRNAs against all zebrafish HOX13 genes pooled together over three biological replicates. Each point represents the tailfin melanocyte area of a different animal. Data was analyzed using a two-sided student’s t-test. * p-value < 0.05, ** p-value < 0.01, *** p-value < 0.001, **** p-value < 0.0001.

**Extended Data Fig. 18:**
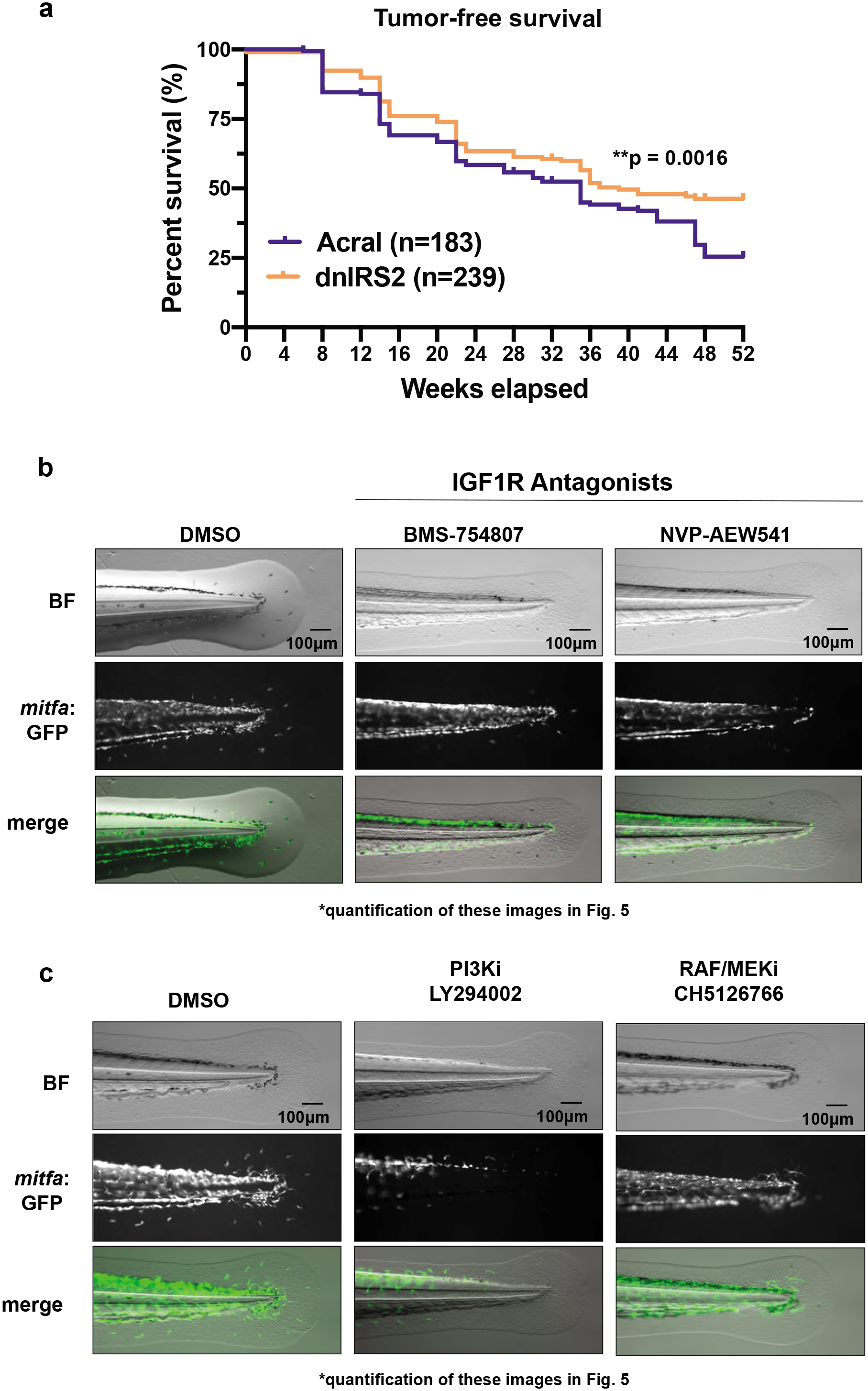
HOX13 synergizes with CRKL to drive acral melanoma through IGF/PI3K signaling. Related to Fig. 5. (a) Tumor-free survival comparing acral to acral dnIRS2 genotypes. Number of fish for each genotype indicated in figure key. P-values generated by log-rank Mantel-Cox test. (b-c) Representative images that correspond to histograms in Fig. 5i-j. Acral melanoma model imaged for melanocyte tailfin area at 3-days post-fertilization after indicated treatment with (b) INSR/IGF1R antagonists BMS-754807 at 7.5μM, NVP-AEW541 at 60μM, (c) PI3K inhibitor LY294002 at 15μM, RAF/MEK inhibitor CH5126766 at 1μM, or 0.1% DMSO control. * p-value < 0.05, ** p-value < 0.01, *** p-value < 0.001, **** p-value < 0.0001.

**Extended Data Fig. 19:**
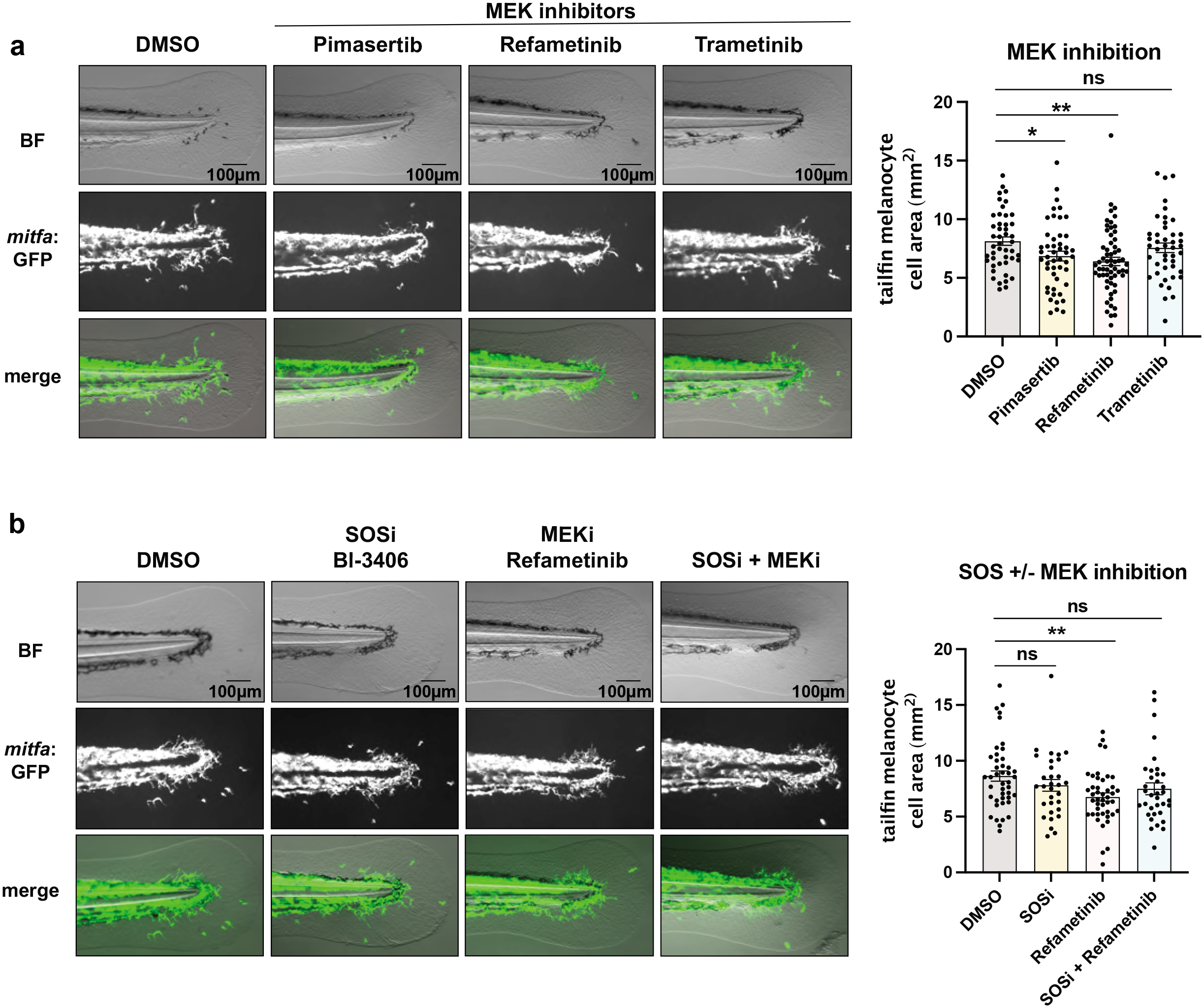
CRKL-mediated tailfin melanocyte expansion has a modest dependence on MAPK signaling. Related to Fig. 5. (a-b) *mitfa*:CRKL stable line imaged for melanocyte tailfin area at 3-days post-fertilization after indicated pharmacologic treatment. P-value generated with two-sided student’s t-test. Error bars = SEM. Each point represents the tailfin melanocyte area of a different animal. MEK inhibitors pimerasertib at 1μM, refametinib at 1μM, and trametinib at 200nM compared to 0.1% DMSO control. Data represents n=46 DMSO-treated fish, n=48 pimasertib-treated fish, and n=61 refametinib-treated fish, and n=44 trametinib-treated fish pooled over four biological replicates. (b) SOS1 inhibitor, Bl-3406, at 1μM, MEK inhibitor, refametinib, at 1μM, or combined Bl-3406 (1μM) + refametinib (1μM) treatment, compared to 0.1% DMSO control. Data represents n=43 DMSO-treated fish, n=31 SOS1 inhibitor-treated fish, n=44 MEK inhibitor-treated fish, and n=35 combined SOS1/MEK inhibitor-treated fish pooled over three biological replicates. * p-value < 0.05, ** p-value < 0.01, *** p-value < 0.001, **** p-value < 0.0001.

**Extended Data Fig. 20:**
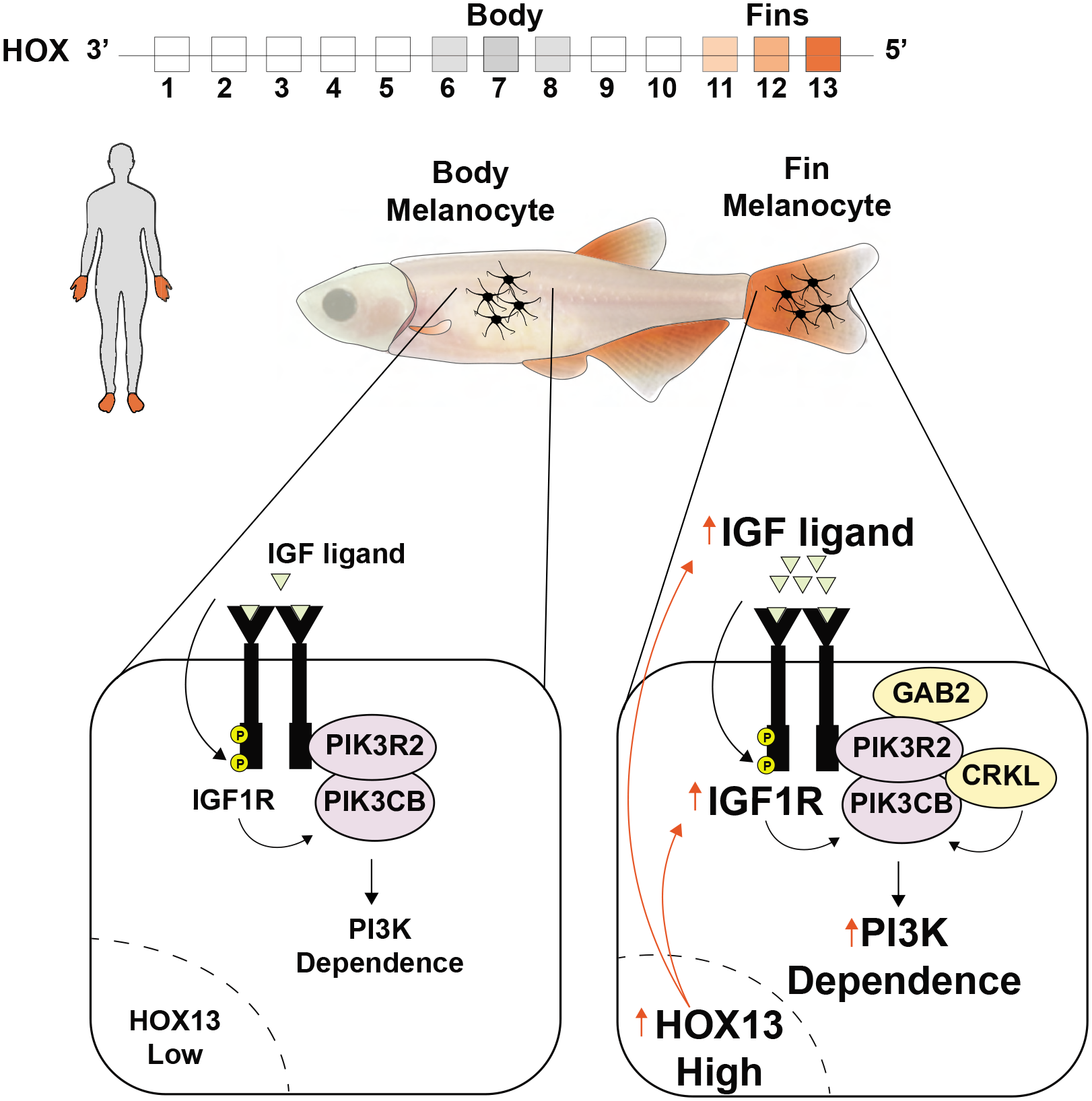
Anatomic position determines oncogenic specificity in melanoma. Related to Figure 5. Melanocytes at different anatomic locations have different positional identities determined by HOX genes. Fin/limb melanocytes have higher expression of posterior HOX13 genes. HOX13 drives higher expression of IGF ligands and IGF1R, resulting in greater IGF signaling and increases the vulnerability of fin melanocytes to CRKL-mediated transformation. CRKL synergizes with a HOX13 program in fin melanocytes by forming a complex with PI3K, the primary downstream mediator of IGF1R, thereby amplifying IGF signaling. GAB2, another commonly amplified gene in acral melanoma, does not directly bind to CRKL, but participates in the same complex as CRKL through their shared binding and amplification of PI3K signaling. This results in tumor phenotypes with sensitivity to IGF1R or PI3K inhibition, but only modest effects from MAPK inhibition.

**Supplemental Table 1: CAMP criteria for diagnosis in zebrafish models of melanoma**

Related to Methods.

**Supplemental Table 2: Primer Sequences**

Related to Methods.

**Supplemental Table 3: Tumor Incidence Anatomic Distribution for all Transgenic Models**

Related to Fig. 1, Fig. 2, and Fig. 5.

**Supplemental Table 4: Zebrafish Fin vs Body RNA-seq**

Related to Fig. 3 and Fig. 5. Differential expression and pathway analysis provided.

**Supplemental Table 5: Human Acral vs Cutaneous RNA-seq**

Related to Fig. 1 and Fig. 4. Differential expression and pathway analysis provided.

**Supplemental Table 6: HOX13 ChIP-seq pathway analysis**

Related to Fig. 5. Pathway analysis results provided.

**Supplemental Table 7: Human Melanoma HOXA13 Cut&Run**

Related to Fig. 5. MACS2 normalized peaks (HOXA13 normalized to IgG control) and pathway analysis results provided.

**Supplemental Table 8: CRKL IP Mass Spec**

Related to Fig. 5. Raw data and data on significant interactors provided.

## Methods

### Resource Availability

#### Lead Contact

Further information and requests for resources and reagents should be directed to and will be fulfilled by Richard M. White.

#### Materials Availability

- Plasmids generated in this study will be deposited to Addgene [MiniCoopR-GFP, *mitfa*:hsCRKL, *mitfa*:hsGAB2, *mitfa*:hsTERT, *mitfa*:Cas9-mCherry;zU6:gRNA-nf1a, *mitfa*:Cas9-mCherry;zU6:gRNA-nf1b, *mitfa*:dnIRS2-GFP, empty-vector/394, empty-vector/395, *mitfa*:Cas9/395, zU6:gRNA-hCRKL, zU6:gRNA-hox13-1, zU6:gRNA-hox13-2, zU6:gRNA-hoxb7a, zU6:gRNA-igf, zU6:gRNA-NT catalog number and unique identifiers pending] and will also be available by request.
- Zebrafish lines generated in this study will be made available through Zebrafish International Resource Center (ZIRC) or by request.
- Cell lines generated in this study are available by request.

#### Data and Code Availability

- All RNA-seq data generated and utilized in this study will be made publicly available.
- Human RNAseq data that support the findings of this study have been deposited in the European Genome-phenome Archive (EGA) and are available under study accession EGAS00001001552 (https://ega-archive.org/studies/EGAS00001001552) with dataset accession EGAD00001006439 (https://www.ebi.ac.uk/ega/datasets/EGAD00001006439). Patient sample information and differential expression tables are made available in the Supporting Information (Supplemental Table 5). Data is also available through Mendeley Data: http://dx.doi.org/10.17632/5f5mf5gw26.1.
- The raw data from the zebrafish fin versus body bulk RNA-seq experiment will be made available via the NCBI GEO repository under identifier code GSE158538, with bulk RNA-seq counts and differential expression tables in the Supporting Information (Supplemental Table 4). Data is also available through Mendeley Data: http://dx.doi.org/10.17632/5f5mf5gw26.1. Samples are annotated as follows:

- MBG -> WT body melanocyte
- MBN -> WT body skin microenvironment
- MFG -> WT fin melanocyte
- MFN -> WT fin microenvironment
- XBG -> Acral body melanocyte
- XBN -> Acral body skin microenvironment
- XFG -> Acral fin melanocyte
- XFN -> Acral fin microenvironment
- The raw data from the zebrafish fin versus body single cell RNA-seq experiment will be made available via the NCBI GEO repository under the identifier code GSE181748. Samples are annotated as follows:

- XBG -> Acral body melanocyte
- XBN -> Acral body skin microenvironment
- XFG -> Acral fin melanocyte
- XFN -> Acral fin microenvironment
- Original source data for ChIP-seq analysis is from ^43^ and can be found at GEO GSE81358. The pathway analysis is made available in the Supporting Information (Supplemental Table 6). Data is also available through Mendeley Data: http://dx.doi.org/10.17632/5f5mf5gw26.1.
- The human melanoma HOXA13 Cut & Run data will be made available via the NCBI GEO repository under identifier code GSE181768. MACS2 peak scores and pathway analysis is made available in the Supporting Information (Supplemental Table 7).
- The CRKL IP mass-spec data will be made available Raw data are available via ProteomeXchange (http://www.proteomexchange.org/) with identifier PXD027968 and is also available in the Supporting Information (Supplemental Table 8).
- Matlab script for tailfin image analysis is available upon request.
- R scripts for RNA-seq, ChIP-seq pathway analysis, and C&R analysis is available upon request.

### Experimental Model and Subject Details

#### Zebrafish

##### Zebrafish Husbandry

Fish stocks were kept under standard conditions at 28.5°*C* under 14:10 light:dark cycles, pH (7.4), and salinity-controlled conditions. Animals were fed standard zebrafish diet consisting of brine shrimp followed by Zeigler pellets. The animal protocols described in this manuscript are approved from the Memorial Sloan Kettering Cancer Center (MSKCC) Institutional Animal Care and Use Committee (IACUC), protocol number 12-05-008. All anesthesia was performed using Tricaine-S (MS-222, Syndel USA, 712 Ferndale, WA) with a 4g/L, pH 7.0 stock. Both male and female zebrafish were utilized in equal proportions for all experiments utilizing adult fish. Sex determination in embryos is not possible at 3 days post fertilization (dpf). Embryos were collected from natural mating and incubated in E3 buffer (5mM NaCl, 0.17 mM KCl, 0.33 mM CaCl2, 0.33 mM MgSO4) at 28.5 °*C*.

##### Generating Zebrafish Transgenic Lines

One-cell-stage embryos of indicated genotype were injected with indicated plasmid (see below) along with 20pg of tol2 mRNA. The total amount of plasmid per injection did not exceed 40pg. Embryos were grown to adulthood at 2 months post-fertilization (2mpf) and then screened for melanocyte rescue. To create stable lines, these fish were outcrossed with casper (mitfa^-/-^, mpv17^-/-^)^71^ or BRAF/p53 caspers fish for two generations (F1 and F2) before in-crossing for F3 stable line generation. The following stable lines were generated in this study: “WT” stable line (MiniCoopR-GFP), acral melanoma stable line (*mitfa*:CRKL, *mitfa*:GAB2, *mitfa*:TERT, *mitfa*:Cas9-mCh;zU6:gRNA-nf1a, *mitfa*:Cas9-mCh;zU6:gRNA-nf1b, MiniCoopR-GFP), CRKL stable line (*mitfa*:CRKL, MiniCoopR-GFP), cutaneous melanoma stable line (*mitfa*:BRAF^V600E^;p53^-/-^;MiniCoopR-GFP, *crestin*:tdTomato). Monitoring for tumor-free survival began at 2mpf and ended at 1 year. For imaging of tailfin melanocytes, embryos at 3dpf were utilized. Fish utilized for FACS and RNA-seq were 6mpf.

##### Zebrafish Transgenic Lines

Zebrafish strains used in these studies included casper (mitfa^-/-^,mpv17^-/-^)^71^ and casper p53^-/-^ with *mitfa*:hBRAF^V600E^ (“BRAF/p53 casper”). The following transgenic lines were all generated using one-cell injection of 20pg of tol2 mRNA and indicated plasmids. The WT melanocyte model was generated by injecting casper fish with 5pg of MiniCoopR-eGFP and 25pg of an empty vector/394b plasmid, which is a plasmid backbone with no transgene. This was used as filler to ensure all conditions were injected with the same total amount of DNA. The cutaneous melanoma model was generated by injecting BRAF/p53 casper fish with 5pg of MiniCoopR-eGFP and 25pg of empty vector/394 plasmid. The acral melanoma model was generated by injecting casper fish with 5pg of each of the following plasmids: MiniCoopR-eGFP, *mitfa*:hsCRKL, *mitfa*:hsGAB2, *mitfa*:hsTERT *mitfa*:Cas9-mCherry;zU6:*nf1a*-gRNA, of mitfa:Cas9-mCherry;zU6:*nf1b*-gRNA. For transgenic models evaluating the role of a single transgene (CRKL, GAB2, and TERT), 5pg of MiniCoopR-eGFP and 25pg of the indicated transgene was used. The NF1 KO melanoma model was generated by injecting 5pg of MiniCoopR-eGFP and 12.5pg of *mitfa*:Cas9-mCherry;zU6-*NF1a*-gRNA and 12.5pg of mitfa:Cas9-mCherry;zU6-*NF1b*-gRNA. For transgenic models evaluating the removal of a given transgene from acral melanoma model, 5ng was used for all indicated plasmids. For transgenic models evaluating the effect of dnIRS2 on the acral melanoma model, 5pg was used for all indicated plasmids. *crestin*:tdTomato reporter was also utilized to aid in identifying tumors to monitor tumor-free survival^72, 73^. For experiments to knockout hCRKL, IGF, and HOX genes in the *mitfa*:CRKL stable line, 10pg of *mitfa*:Cas9/395 along with 10pg along with the following indicated sgRNA plasmids: zU6:gRNA-hCRKL, zU6:gRNA-igf, zU6:hox13-1, zU6:hox13-2, zU6:hox7ba, or zU6:gRNA-NT. Genotypes were regularly monitored by PCR.

#### Humans

##### MSK-IMPACT DNA Sequencing

Genomic analysis of human acral and cutaneous melanoma patients came from a broad institutional effort to sequence cancer patients that arrive at MSK and make available for the clinical and scientific community^20, 22^. Patients analyzed were clearly indicated as having either acral or cutaneous melanoma. Melanoma patients without subtype designations or with other subtypes (i.e. uveal or mucosal) were not included. Patients of all sexes, ethnicities, and ages were included for analysis.

##### WES and WGS of an Individual Acral Melanoma Patient

The female patient was enrolled at Memorial Sloan-Kettering Cancer Center (MSKCC) and consented on a protocol approved by MSKCC and analyzed as part of a previous acral melanoma study^15^. Samples were obtained in accordance with standard biopsy or surgical procedures.

##### RNA-sequencing of Human Melanoma Patients

For human RNA-seq sequencing analysis, fresh-frozen tissue samples were obtained from the biospecimen bank of Melanoma Institute Australia (MIA) and all samples were accrued prospectively with written informed patient consent. The protocol for the study was approved by the Sydney Local Health District Ethics Committee (Protocol No X15-0454 (prev X11-0289) & HREC/11/RPAH/444 and Protocol No X17-0312 (prev X11-0023) & HREC/11/RPAH/32) and cases were also approved by institutional ethics committees of Melanoma Institute of Australia and QIMR Berghofer Medical Research Institute (HREC approval P452 & P2274). Details of patient samples can be found in Supplemental Table 5. The acral RNA-seq samples have been previously published in^13^.

##### Acral melanoma Tissue Microarray

The human acral melanoma tissue microarray was made from patients consented to IRB protocol 06-107 at Memorial Sloan Kettering Cancer Center. Punch biopsies were taken from viable tumor areas and embedded in paraffin.

#### Cell Lines and Virus Preparation

WM3918 melanoma cell line was grown with Dulbecco’s Modified Eagle Medium (Gibco #11965) supplemented with 10% FBS (Seradigm), 1X penicillin/streptomycin/glutamine (1X PSG) (Gibco #10378016). SKMEL-1152, SKMEL-1136, SKMEL-1094, SKMEL-1128, SKMEL-1088, SKMEL-1176, and SKMEL-1206 are patient-derived human melanoma cell lines established at Memorial Sloan Kettering Cancer Center. SKMEL-1152, SKMEL-1136, SKMEL-1094, and SKMEL-1128 are derived from cutaneous melanoma patients and SKMEL-1176 and SKMEL-1206 are derived from acral melanoma patients. SKMEL-1088 is derived from a human melanoma of unknown subtype. MSK-IMPACT sequencing was performed to identify their putative genetic drivers. These patient derived cell lines were grown in RPMI 1640 Medium (Invitrogen #11875093) supplemented with 20% FBS (Seradigm), 1X penicillin/streptomycin/glutamine (Gibco #10378016). All cell lines were kept in a sterile 37℃, 5% CO_2_ incubator.

To make CRKL-overexpressing cell line, CRKL retrovirus was produced by transfecting 293T packaging cells with CRKL-pWzl, VSVg and pCL-Ampho plasmids. WM3918 human melanoma cell line was transduced with CRKL retroviral particles plus polybrene and then selected with 10 μg/mL blasticidin. To make CRKL-V5-overexpressing cell lines and empty vector (EV) control cell lines, CRKL-V5 and EV lentivirus was produced by transfecting 293T packaging cells with pLenti6.3-CRKL-V5 or pLenti6.3-EV and Ready-to-Use Lentiviral Packaging Plasmid Mix (Cellecta #CPCP-K2A). WM3918, SKMEL-1176, MeWo, and 293T cell lines were transduced with CRKL lentivirus particles plus polybrene and then selected with 10μg/mL blasticidin. CRKL-pWzl, and pLenti6.3-CRKL-V5 was a kind gift from professor William Hahn.

### Method Details

#### Acral Enrichment Score Analysis of MSK-IMPACT DNA Sequencing

Only melanoma patients clearly indicated as having either acral or cutaneous melanoma were analyzed. We selected the 14 most frequently observed copy number alterations and 7 most frequently mutated genes in acral melanoma and compare to the frequency observed in cutaneous melanoma. We also included well described melanoma drivers, such as deletion of NF1^74^ and mutations in PTEN^75^. To calculate the acral enrichment score we used the following equation log2(acral frequency/cutaneous frequency). Statistical differences in the frequency of copy number alterations and mutations were determined using a Fisher’s exact test.

#### WES/WGS Analysis of an Acral Melanoma Patient

Tissue was selected by the pathologist to limit the amount of necrotic tissue and adjacent normal tissue was collected for DNA extraction of germline DNA. An H&E slide was made to confirm normal or malignant tissue and <50% necrosis, which was reviewed by the pathologist. The patient had both a primary tumor sample, which was formalin-fixed and paraffin embedded (FFPE) and an in-transit metastasis, which was fresh frozen by placing in a vial and submerging into a liquid nitrogen container. Paired tumor/normal whole-exome and whole-genome sequencing (WES and WGS, respectively) libraries were constructed and sequenced on the Illumina HiSeq using V3 reagents. WES and WGS data was aligned to the genome with BWA ^76, 77^ and mutations called using MuTect^78^. Coverage-based copy-number analysis was performed using custom scripts for WGS and using the allele-specific method FACETS for the WES data^79^.

#### RNA-seq Analysis of Human Melanoma Patients

Fresh-frozen tumor RNA was extracted using the AllPrep® DNA/RNA/miRNA Universal kit (Qiagen #80224) and were quantified using the Qubit® RNA HS Assay (Q32852, Life Technologies). The TruSeq RNA library prep kit (Illumina, San Diego, California, USA) was used to prepare libraries from RNA and these were sequenced with 100bp paired-end reads using Illumina HiSeq2000 or HiSeq2500 platforms. RNA-seq reads were trimmed for adapter sequences using Cutadapt (version 1.9) and aligned with STAR (version 2.5.2a)^80^ to the GRCh37 assembly using the gene, transcript, and exon features of Ensembl (release 70) gene model. Gene expression counts were estimated using RSEM (version 1.2.30)^81^. Differential expression was calculated with DESeq2^82^ using the output of the quantMode and GeneCounts feature of STAR. The vst function was used to generate log2 transformed normalized counts. Differential expression for Figure 5D-F, including HOMER analysis used comparison of all acral vs cutaneous melanoma samples. Differential expression used for pathway analysis for Figure 5B-C used comparison of all acral vs cutaneous melanoma samples normalized by specimen type (i.e. primary tumor or lymph node metastasis). Pathway and Gene Ontology (GO) analysis were performed with GSEA using FGSEA-multilevel^83^. Known motif analysis was performed with the HOMER ^40^ function findMotifs.pl, using the human genome (GRCh37) and searching for motifs of lengths 8, 10, and 16 within ± 500bp of the TSS of differentially expressed genes. Motifs were annotated using JASPAR ^84^.

#### Co-Occurrence of CRKL, GAB2, NF1, and TERT in Cancer

The frequency of alterations occurring across multiple human cancer types was analyzed in cBioPortal^19, 21^ looking at the following TCGA studies: Firehose Legacy ^85^, PanCancer Atlas ^86^, Cell 2017^87–90^, Nature 2014^91–93^, Nature 2012^94–96^, Cell 2015^74, 97, 98^, Nature 2008^99^, Nature 2015^100^, Nature 2013^101, 102^, Cancer Cell 2014^103^, NEJM 2013^104^, Nature 2011^105^, Cell 2013^106^, Cell 2014^107^. The odds ratio for co-occurrence of alterations was calculated and used to generate a p-value adjusted for FDR = 0.05.

#### HOXA13/HOXD13 ChIP-seq Pathway Analysis

ChIP-seq analysis was performed from raw reads of publicly available GEO datasets from Sheth et al, 2016 (GSE81358). Paired end reads from the following samples were processed: GSM2151011 (H3K27ac), GSM2151013 (HOXA13), GSM2151014 (HOXD13), GSM2151016 (H3K27ac input DNA), GSM2151017 (HOXA13 input DNA), GSM2151018 (HOXD13 input DNA). SRA files were converted to FASTQ format using the SRA toolkit. Raw reads were checked for sequence quality, adapter content, overrepresented sequences and Kmer content using FASTQC (Babraham Bioinformatics). Adapters and low quality sequences were filtered out using Trimmomatic^108^ and filtered reads were mapped to the mm9 genome using Bowtie2^109^. Mapped reads were analyzed using Samtools^110^ and the sorted bam files were processed using Deeptools^111^ to generate input normalized bigwig files. MACS v1.4^112^ was used to perform peak calling using input DNA for each antibody as control. Pathway analysis of enriched peaks was performed using Cistrome-GO ^44^.

#### HOXA13 Cut&Run of Human Melanoma Cell Lines

##### Sample Preparation

We performed Cut&Run for the melanoma cell lines SKMEL-1088, SKMEL-1766, and SKME-1206. For Cut&Run we used 100’000 cells per condition and it was performed as described in^113^. We used antibodies against HOXA13 (Invitrogen, #PA5-76440, 1:100), H3K27ac (Active Motiv, #39034, 1:100), IgG (abcam, #ab6709, 1:100) and we added a “no antibody” condition as additional negative control. In brief, 100’000 melanoma cells were collected per condition. Cells were harvested and bound to concanavalin A-coated magnetic beads after 8min incubation at RT on a rotator. Cell membranes were permeabilized with digitonin and the different antibodies were incubated overnight at 4°C on a rotator. Beads were washed and incubated with pA-MN. Ca2+-induced digestion occurred on ice for 30min and stopped by chelation. DNA was isolated using an extraction method with phenol and chloroform as described in Skene et al.^113^

##### Sequencing

Immunoprecipitated DNA was quantified by PicoGreen and the size was evaluated by Agilent BioAnalyzer. When possible, fragments between 100 and 600 bp were size selected using aMPure XP beads (Beckman Coulter catalog # A63882) and Illumina libraries were prepared using the KAPA HTP Library Preparation Kit (Kapa Biosystems KK8234) according to the manufacturer’s instructions with up to 1.5ng input DNA and 12-16 cycles of PCR. Barcoded libraries were run on the NovaSeq 6000 in a PE100 run, using the NovaSeq 6000 SP Reagent Kit (200 cycles) (Illumina). An average of 13 million paired reads were generated per sample.

##### Analysis

Reads were trimmed and filtered for quality (q=15) and adapter content using version 0.4.5 of TrimGalore (https://www.bioinformatics.babraham.ac.uk/projects/trim_galore). Reads were aligned to human assembly hg38 with version 2.3.4.1 of bowtie2 (http://bowtie-bio.sourceforge.net/bowtie2/index.shtml) and duplicates were collapsed to one read using MarkDuplicates in version 2.16.0 of Picard Tools. CUT&RUN target enrichment was assessed using MACS2 (https://github.com/taoliu/MACS) with FDR=0.1 and fold change of 2 over the matched IgG control background. Depth-normalized read density profiles were created using the BEDTools suite (http://bedtools.readthedocs.io). A global peak atlas was created by first removing blacklisted regions (http://mitra.stanford.edu/kundaje/akundaje/release/blacklists/hg38-human/hg38.blacklist.bed.gz) then merging all overlapping peaks. Peak-gene associations were created by assigning peaks using linear genomic distance to transcription start sites. Motif signatures were obtained using Homer v4.5 (http://homer.ucsd.edu). Pathway analysis of enriched peaks (from MACS2) was performed using Cistrome-GO with the following parameters: peak number to use: all, half-decay distance: automatic, FDR cutoff: 0.2.

#### CRKL IP-mass spec

##### Sample preparation

Two cell lines were used for IP-MS/MS: Human melanoma cell line WM3918 with V5-tagged CRKL overexpression and a control cell line with wildtype (no V5-tag) CRKL overexpression. For each cell line IP was performed in six replicates. 100 x 10^6^ cells were thawed at 37C for 30 sec lysed in 2ml of ice cold 50mM EPPS pH 7.5, 150mM NaCl, 1% triton, cOmplete™, EDTA-free Protease Inhibitor Cocktail (1 tablet per 20mL of lysis buffer), 1:100 of of sigma phosphatase inhibitors 2 and 3 cocktails and 250U/μL of benzonase. Lysates were incubated on ice for 5 min to allow DNA digestion, centrifuged at 20K g for 5 min, to remove insoluble material and filtered through acroprep 1.0um glass filter plate at 2000g for 1 minute. The concentration of protein was then estimated by BCA and immunoprecipitation was performed in 2ml deep well plates with 1mg of protein material, 5.75ug of V5 antibody (Invitrogen #R960-25) bound to 5.75μL of Protein G Sepharose (17-0618-02). The final volume of IP reaction was 300μL. The incubation was performed at 4C shaking at 1100 rpm for 1 hour. The beads were then transferred to OF 1100 filter plate (Orochem) and washed 5 times with ice cold 50mM EPPS pH 7.5, 150mM NaCl using vacuum manifold. 18μL of 10mM EPPS pH 8.5 with 20ng/μL trypsin, 10ng/μL LysC was added to the beads in each well and digestion was performed for 2 hours at 37C at 2000rpm. The partial digest was then collected into 96 well PCR plate and left overnight at room temperature to complete digestion. 4μL s of 22g/L 11 plex TMT tags were added to each sample (6 replicates of IP for V5-tagged CRKL overexpression sample and 5 replicates of IP from wildtype CRKL overexpression cell line was combined in the TMT 11 plex experiment). The samples were then pulled and 20μL of the combined sample was set aside, while the rest was fractionated into 8 fractions using High pH Reversed-Phase Peptide Fractionation Kit, as suggested by manufacturer. The fractions were concatenated into 4 fractions (1st and 5th fractions, 2nd and 6th and so on) and evaporated in speed vac (1μL of DMSO was added to each sample to prevent complete evaporation) and resuspended in 20 μL 0.1%TFA Data acquisition

5μL of unfractionated sample and every fraction was analyzed by EASY-nLC 1200 System with 2mm particle size, 75mm x 500 mm easyspray column in direct injection mode. The samples were separated using the following gradient at 300nl/min of buffer A (0.1% formic acid in water) and buffer B (0.1%formic acid in acetonitrile): 0%–5% in 10 min, 5%–25% in 92 min, 25%–50% in 18min, the column was then washed with 95% B for 10 min at 400nl/min. Eluting peptides were analyzed on Orbitrap Fusion Lumos mass spectrometer using MS3 SPS with the settings recommended by the instrument manufacturer for TMT11 plex analysis. Cycle time was set at 3 sec and exclusion time was set at 45 sec. Data analysis Data were analyzed in Proteome Discoverer 3.1 software. A database search was performed with Sequest HT search engine using Human UniProt database containing only reviewed entries and canonical isoforms (retrieved on 06/14/2019). Oxidation (M) was set as a variable modification, while TMT6plex was set as fixed modification. A maximum of two missed cleavages were permitted. The precursor and fragment mass tolerances were 10 ppm and 0.6 Da, respectively. PSMs were validated by percolator with a 0.01 posterior error probability (PEP) threshold. Only PSMs with isolation interference < 25% and at least 5 MS2 fragments matched to peptide sequence among the 10 selected for MS3 were considered. The quantification results of PSMs were combined into protein-level quantitation using MSstatsTMT R package^114^. Only proteins with at least 3 peptides were reported. Significant interactors were defined by having an adjusted p-value < 0.05 and fold change > 2 compared to control. Proteins were then analyzed for previously described SH2 and SH3 binding motifs^1^^15,1^^16^.

#### Plasmid Construction

The following plasmids were constructed using the Gateway Tol2kit.

*mitfa*:hsCRKL/394
*mitfa*:hsGAB2/394
*mitfa*:hsTERT/394
*mitfa*:Cas9-mCherry;zU6:*NF1a*:gRNA/394
*mitfa*:Cas9-mCherry;zU6:*NF1b*:gRNA/394
*mitfa*:dnIRS2-GFP/394
empty-vector/394
empty-vector/395
*mitfa*:Cas9/395
zU6:gRNA-hCRKL/394
zU6:gRNA-hox13-1/394
zU6:gRNA-hox13-2/394
zU6:gRNA-hoxb7a/394
zU6:gRNA-igf/394
zU6:gRNA-NT/394

cDNAs in tol2-compatible pENRT223.1 plasmids were ordered through Horizon Discovery for the following genes.

hsCRKL (Clone ID: 100000145)
hsGAB2 (Clone ID: 56810)
hsTERT (Clone ID: 100061944)

sgRNA sequences

*nf1a*: GGCGCACAAGCCCGTGGAAT
*nf1b*: GGCGCAGAAGCCCGTGGAGT
*hoxa13a*: GGGCAATCACAACCAGTGGA
*hoxa13b*: GGATGATATGAGCAAAAACA
*hoxb13a*: GCGAGGATTCAGGACCAGGG
*hoxc13a*: CCGTGATATGACGACTTCGC
*hoxd13a*: GGCTCTGGCTCCTTCACGTT
CRKL-1: CGCGGACGAGGAACATACCG
CRKL-2: CAACCGCCGTTTTAAGATCG
CRKL-3: GTCGGTGTCCGAGAACTCGC
*igf1*: TCTAGCGGTCATTTCTTCCA
*igf2a*: TGCATCTTGCCGAAAAACGG
*igf2b*: GAAACTGTCTGTTCTCGAGC
Non-target (NT): AACCTACGGGCTACGATACGCGG

Zebrafish dnIRS2-GFP was a generous gift from ^56^ and then further cloned into pENTR/D-TOPO vector.

#### Zebrafish Genotyping by PCR

Tail clips from adult zebrafish were placed in microcentrifuge tubes or thermal cycler plates containing 50 ml of 50 mM NaOH. Samples were boiled at 95°C for 30min, then cooled down with 5 ml of 1M Tris-HCL (pH=8.0). A 1:10 dilution of the supernatant was used in PCR. DNA was PCR amplified with Promega GoTag green mastermix (Promega #M7123). PCR amplicon sizes for CRKL, GAB2, TERT, and Cas9 are 221bp, 125bp, 296bp, and 207bp. All transgenes have the same forward primer. GTTGAACGCAAGTTTGTACA going off the *mitfa* promoter Primer sequences are listed below.

CRKL: TGATGTAGTGGGAGACCCGC
GAB2: AGTATTCCAGAACATCTGGG
TERT: AGGCAGGACACCTGGCGGAA
Cas9-mCherry: CATGTGCACCTTGAAGCGCA

#### CRISPR-seq

Genomic DNA of transgenic zebrafish was isolated via tail clip genotyping described above. DNA was PCR amplified with Phusion polymerase (NEB #M0531S), run on an agarose gel, and then gel purified using NucleoSpin® Gel & PCR Clean-up Midi (Takara # 740986.20) for deep sequencing using the CRISPR-seq platform^117^. Primers used are listed below. Sequencing data was aligned to the zebrafish genome (GRCz10) and analyzed with CrispRVariantsLite version 1.2 ^118^.

*nf1a*: TCGGGATCGCAAAAGTGATT CACCAAGCTCACATCTTCAA
*nf1b:* CACCATCTTCATCATCCTCCT ACTACTCTCTGTCCCGTGTC

#### Surveyor for sgRNA Validation

Recombinant Cas9 (IDT #1081059) and sgRNA were complexed and injected into Casper embryos at the one-cell stage. Briefly, 100 μM crRNA and tracrRNA (IDT #1075928) were mixed at 1:1 ratio and incubated at 95C for 5 minutes and then cooled to room temperature. 1.11μL of sgRNA duplex, 0.9μL (9μg) of recombinant Cas9, and 2.99μL of duplex buffer (IDT #11-05-01-12) were incubated at 37C for 10minutes. After incubation 2μL of 100ng/μL tol2 mRNA, 1μL of 100ng/μL empty-vector/395 and 2μL of phenol red (Sigma-Aldrich #P0290) were added to sgRNA:Cas9 complex. This mix was then used to inject 1pL into Casper embryos at the one-cell stage, resulting in delivery of 0.9pg of Cas9 protein, 10pg of empty-vector/395, and 20pg of tol2 mRNA. At 24hpf, embryos were sorted for the GFP+ heart marker from successful 395-empty incorporation. 10 embryos were used for genomic DNA isolation using the Qiagen DNAeasy Blood & Tissue kit (Qiagen #69504). DNA was PCR amplified with Promega GoTag green mastermix (Promega #M7123). Surveyor was performed using IDT Surveyor kit (IDT #706020) following manufacturer’s instructions. Primers used for each gene are listed below:

*hoxa13a:* GGGTGATTCTGGAAAGCAAT CTCCATGGGATACTGACTCT
*hoxa13b:* ATCCCATTGTGCAATGGAAT CGCCAAAATATCCATAGGGC
*hoxb13a:* TTGACATTCTTCACCCAAGG AGCCCTGGTAGGATATTCTT
*hoxc13a:* GCCAGTAGTTGTTTAAAGGG GTCTCGGCATATTTTTCTGC
*hoxc13b*: TAGTGAAAGACGTTTGCGTT ACAACTGGGACGTCCAAATA
*hoxd13a:* ATGCACTGAGGAATATGGAC CTAATGAAGAGAGGCGAGGA
*hoxb7a:* ATATATCATCACGTGCTGCC CTCTACATACACAGACGCAC
*CRKL-1:* TTCGAATAAACATGTCGTCTGC CGCCTATCTTGAATCTCTTGCT
*CRKL-2:* TCCACTTGTCCTGGTGATTATG TTCTATCAGGGTGGTCGTATCC
*CRKL-3*: CTCCACTAGCGAGAAGCTGATT TCTTGAAAGGAAGGTCTTCAGC
*igf1:* GGATTTTCTCTCCAAATCCG CTTAATCATGTCGACTCAGC
*igf2a:* CAAAAGAACCACTCGTTCAC CACACGAACTGCAGTGTATC
*igf2b:* CTCTCAGAGAACTTTTGCCT GCGTATCCAGAACGTAATGT

#### RT-PCR Validation of Transgene Expression

Wildtype melanocyte skin, acral melanoma model tumor, and cutaneous melanoma model tumor tissue were dissected from zebrafish and RNA was isolated with the Zymo Quick RNA Miniprep kit (#R1050) using kit protocol. cDNA was synthesized using SuperScript III First Strand Synthesis System (Life Technologies #18080-400) following the kit protocol. RT-PCR was performed using iQ Sybr Green Supermix (Biorad #1708882). RT-PCR primer sequences are listed below:

*mitfa*: GCCCTATGGCCCTTCTCAC CATCCATGAACCCAAGAATGTCA
hsGAB2: GCGGCGACGTGGTGT CTTCCAGGCATAGCGCCTC
hsCRKL: Sino Biological qPCR primer pairs (HP101145)
hsTERT: GGAGCAAGTTGCAAAGCATTG TCCCACGACGTAGTCCATGTT
*beta-actin*: GCCAACAGAGAGAAGATGACAC CAGAGAGAGCACAGCCTGG

#### Western Blot

Zebrafish lysates were prepared by sonication in RIPA buffer (Thermo #89901) with 1X Halt Protease and Phosphatase Inhibitor Cocktail (Thermo #78441) followed by centrifugation (14,000rpm for 10min at 4°C) and collection of the supernatant. Protein concentration was quantified by Bradford (Sigma B6916-500mL) according to manufacturer’s protocol. Samples were mixed with 6X reducing loading buffer (Boston BioProducts #BP-788 111R) and denatured at 95°C for 10 minutes. Samples were run on a Mini PROTEAN TGX gel (BioRad) and transferred using Turbo Mini Nitrocellulose Transfer Pack (Bio-Rad, catalog #1704158). Membranes were blocked with 5% nonfat dry milk in TBST (1X TBS + 0.1% Tween 20) for 1 hour before incubation with primary antibody in PBS overnight at 4°C. Membranes were washed with TBST and incubated with secondary antibody in 5% nonfat dry milk for 1 hour at room temperature. Membranes were washed with TBST and developed with ECL (Amersham, RPN2109) using an Amersham Imager 600 (GE) or chemiluminescence film.

Cell lines were cultured in standard media conditions (RPMI 1640 Medium (Invitrogen #11875093) supplemented with 20% FBS (Seradigm), 1X penicillin/streptomycin/glutamine (Gibco #10378016)) prior to collection. Cells were washed with cold PBS and lysed in RIPA buffer (Pierce #89901) plus phosphatase and protease inhibitors (Thermo Scientific #1861277, #1861278). Lysates were cleared by centrifugation at 14000rpm at 4°C and quantified using BCA method (Pierce #23224). Samples were prepared using LDS+Reducing agent Novex buffers (Invitrogen #NP0008, #NP0009). 10 to 20μg of lysates were loaded and run on NuPageTM 4-12% Bis-Tris gels (ThermoFisher #NP0321BOX) followed by transfer to nitrocellulose membranes (Biorad #1620233). Membranes were incubated over night with the indicated antibodies, washed and incubated again for 45 minutes with anti-rabbit or anti-mouse secondary antibodies. Detection was performed using Immobilion Western (Millipore #WBKLS0500).

Primary antibodies are: CRKL (32H4) #3182S (Cell Signaling Technology) (used for fish lysates), CRKL (B-1) sc-365092 Lot#B0819 (Santa Cruz Biotechnology) (used for human cell lines), GAB2 (26B6) #3239 (Cell Signaling Technology), IGF-1R beta (D23H3) #9750S (Cell Signaling Technology), phospho-IGF-1R beta Y1135 (DA7A8) #3918S (Cell Signaling Technology), HOXB13 (D7N8O) #90944 (Cell Signaling Technology), IGF2 #ab170304 (Abcam), and β-Actin (AC-74) #A2228 (Sigma-Aldrich). All the primary antibodies were used at 1:1000 dilution except for β-Actin that was used at 1:10,000 dilution. All secondary antibodies were used at 1:10,000. Secondary antibodies include goat anti-rabbit IgG #ab97051 (Abca), rabbit anti-mouse IgG #ab97046, rabbit anti-goat IgG #HAF017 (R&D Systems).

#### Phospho-RTK array

Human Phospho-RTK arrays (R&D Systems, ARY001B) were used to detect activated RTKs according to manufacturer’s instructions. WM3918 parental and WM3918-CRKL cell lines were serum starved for 24 hours and then stimulated with 20% FBS (Seradigm) for 10 minutes before collection. Cells were washed with cold PBS and lysed using the provided lysis buffer plus phosphatase and protease inhibitors. Lysates were cleared by centrifugation at 14000rpm at 4°C and quantified using BCA method (Pierce #23224). 200 μg lysates were incubated on membranes overnight. Membranes were subsequently washed and exposed to chemiluminescent reagent (Millipore #WBKLS0500).

#### HOX13 siRNA Knockdown

Three human acral melanoma cell lines including SKMEL-1088, SKMEL-1176 and SKMEL-1206 were used for knockdown of four HOX13 genes, respectively. Non-targeting Control siRNA (D-001810-01-05), HOXA13 siRNA SMARTPool (#L-011052-00-0005), HOXB13 siRNA SMARTPool (#L-012226-00-0005), HOXC13 siRNA SMARTPool (#L-017600-00-0005) and HOXD13 siRNA SMARTPool (#L-011053-00-0005) were purchased from Horizon Discovery. Briefly, 4 × 10^5^ cells were seeded in 6-well plates and cultured with RPMI 1640 medium (ThermoFisher # 11875093) containing 10% FBS (Seradigm), 1X PSG (Gibco #10378016) for 24-hr. Media was then replaced with 1ml of fresh medium before transfection. 100nM of total siRNA (25nM of each HOX siRNA) were added into 200μL of Gibco Opti-MEM (ThermoFisher #<u>31985070</u>) and mixed with 12μL l DharmaFECT Duo Transfection Reagent (Horizon Discovery #T-2010-03) followed by a 10-sec agitation. After 10-min incubating at room temperature, transfection mixtures were evenly dropped into each well. Media was aspirated 6-hr later and replaced with 2ml fresh RPMI medium containing 10% FBS, 1X PSG. After 48 hours post-transfection, cells were changed to 3mL of RPMI 20% FBS 1X PSG for Western blot or 1.5mL of serum-free RPMI 1X PSG for ELISA. Samples were collected 96 hours post-transfection for both Western blot and ELISA.

#### IGF2 ELISA

SKMEL-1088 and SKMEL-1176 were prepared as described above. At 96 hours post-transfection, cell supernatants were collected and spun at 1000G for 5min to remove cell debris. IGF2 concentration was determined by using the Human IGF2 Quantikine ELISA Kit (R&D Systems #DG200) following manufacturer’s instructions.

#### Zebrafish Imaging Tumor-free Survival Curves

Zebrafish embryos were injected at the one-cell stage with indicated plasmids. *crestin*:tdTomato reporter was also utilized to aid in identifying tumors to monitor tumor-free survival^72, 73^. All injected embryos were grown to adulthood and then screened for melanocyte rescue at 2mpf based on a combination of GFP fluorescence and pigmentation. Zebrafish were regularly monitored every 2-4 weeks for the development of new tumors. Tumors were called based on the CAMP criteria outlined in Table S1. Kaplan Meir curves were generated and analyzed in Prism version 8. Statistical differences in tumor-free survival were determined by log-rank Mantel-Cox test.

#### Analysis of Anatomic Distribution of Tumors

During monitoring of tumor-free survival, the anatomic location (head vs body vs fins) of the tumor is noted. If a fish has either >2 tumors or a tumor that encroaches on the border of multiple anatomic sites, all tumors and anatomic sites are counted. To compare the anatomic distribution of tumors, a Chi-squared test was performed using Prism software version 8. Ternary diagrams were generated using ternaryplot.com.

#### Histology of Zebrafish and Human Samples

Zebrafish were sacrificed using ice-cold water. The head and tail were separated from the body separated via dissection with a clean razor. The sample of interest was then placed in 4%PFA (Santa Cruz #30525-89-4) for 72 hours on a shaker at 4°C and then immediately embedded in paraffin blocks to generate 5-micron sections. Human acral melanoma tissue TMA was generated as described above. Human and zebrafish samples were processed and stained using the same protocols described below. All H&E and IHC staining was performed at Histowiz. Tissue was put through a heat-induced epitope retrieval process (100°C), followed by a proprietary automatic staining assay developed for the BOND RX automated stainer (Leica Biosystems Division of Leica Microsystems Inc, Buffalo Grove, Illinois). Information about the antibodies used and staining conditions is listed below. The staining process included the use of the BOND Polymer Refine Red Detection Kit (Leica Biosystems) in accordance with the manufacturer’s protocols, utilizing alkaline phosphatase and Fast Red chromogen. Endogenous alkaline phosphatase activity was blocked using a levamisole solution (Vector Laboratories). After staining, sections were dehydrated and film coverslipped using a Tissue-Tek Film automated Coverslipper (Sakura Finitek USA Inc, Torrance, California). Whole slide scanning (×40 objective) was performed on an Aperio AT2 digital whole slide scanner (Leica Biosystems). Human acral melanoma TMA stains for CRKL, HOXB13, and pIGF1R were analyzed and scored by a dermatopathologist at Memorial Sloan Kettering Cancer Center. Samples were scored for extent based on the following criteria: 0 = negative, 1= 1-25% of tumor cells are positive, 2 = 26-50% of tumor cells are positive, 3 = 51-75% of tumor cells are positive, and 4 = 76-100% of tumor cells are positive. Intensity was scored, 0 = negative, 1 = weak staining, 2 = intermediate staining, 3 = strong staining. N = 2/32 samples were removed from the analysis due to heavy pigmentation obscuring detection of staining signal, leading to an analysis of n = 30 samples total.

| Antibody | Vendor | Catalog # | Host | Clonality | Dilution | Antigen Retrieval |
| --- | --- | --- | --- | --- | --- | --- |
| CRKL | Sigma | HPA001100 | Rabbit | Polyclonal | 1:50 for fish<br>1:25 for human | pH9<br>20mins |
| HOXB13 | Cell Signaling | CST90944 | Rabbit | Monoclonal [D7N8O] | 1:50 for fish<br>1:50 for human | pH9<br>20mins |
| p-IGF1R | Abcam | ab39398 | Rabbit | Polyclonal | 1:100 for fish<br>1:100 for human | pH9<br>20mins |
| GFP | Abcam | ab183734 | Rabbit | Monoclonal [EPR14104] | 1:100 for fish | pH6<br>20mins |
| p-ERK | Cell Signaling | CST4370 | Rabbit | Monoclonal [D13.14.4E] | 1:100 for fish | pH6<br>20mins |
| p-S6 | Cell Signaling | CST4858 | Rabbit | Monoclonal [D57.2.2E] | 1:200 for fish | pH6<br>20mins |

#### IF of Zebrafish Melanoma Samples

Unstained histology slides of zebrafish samples were deparaffinized using 2×10 minutes xylene, 4 minutes 100% ethanol, 1 minute 95% ethanol, 1 minute 70% ethanol, 1 minute 50% ethanol, and then rinsed with water. Antigen retrieval was achieved by placing slides in 10mM sodium citrate pH = 6.2 by heating in the microwave for approximately 2 minutes until the solution begins to bubble. The slides then sat for 5 minutes and then were microwaved again for 40 seconds. The slides cooled down to room temperature for 30 minutes in the same buffer. The slides were blocked in a blocking solution consisting of 5% donkey serum (Millipore Sigma #S30-M), 1% BSA, 0.4% Triton X-100 (Fisher # PI85111) in PBS. 50uL of blocking buffer was added per section. The blocking solution was incubated at room temperature for 1 hour. Primary antibodies were diluted at 1:100 in blocking solution and incubated overnight at 4°C. Slides were washed 3 x 5 minutes in PBS and then applied with secondary antibody solution 1:250 in blocking solution. Hoescht (Fisher # H3570) was added to be 1:1000 dilution. The slides were washed again 3 x 5 minutes in PBS. Coverslips were mounted onto the slides using Vectashield Vibrance Antifade Mounting Medium (Vector Laboratories #H-1800-2). Slides were then imaged on a confocal microscope. Primary antibodies used were Sox10 (GeneTex #GTX128374) and GFP (Abcam #5450). Secondary antibodies used were Alexa-anti Rabbit 594 (Cell Signaling Technology #8889) and Alexa-anti Goat 488 (ThermoFisher #A32814).

#### Pharmacologic Treatment of Zebrafish Embryos

Zebrafish embryos were collected at 1dpf and placed in a 40-micron cell strainer (Thermo Fisher # 08-771-2) in a 6-well dish (Fisher #08-772-1B) in 6mL of E3 water. 20 embryos were used per well. Zebrafish embryos were treated with the indicated compounds: insulin/IGF1 receptor antagonists BMS-754807 (Sigma-Aldrich #BM0003-5MG) at 7.5μM and NVP-AEW541(Selleckchem # S1034) at 60μM, PI3K inhibitor LY294002 (Sigma-Aldrich #L9908-1MG) at 15μM, RAF/MEK inhibitor CH5126766 (Selleckchem # S7170) at 1μM, MEK inhibitor Pimasertib (Selleckchem #S1475) at 1μM, MEK inhibitor Refametinib (Selleckchem #S1089) at 1μM, MEK inhibitor Trametinib (Selleckchem #S2673) at 200nM, and SOS1 inhibitor Bl-3406 at 1μM (MedChemExpress #HY-125817). Compound stocks were kept at 1000X in DMSO. Treatment started at 1dpf and reapplied at 2dpf. Zebrafish were imaged for tailfin area at 3dpf.

#### Measurement of Zebrafish Melanocyte Cell Area in Tailfin

The extent of melanocyte area in the tailfin was calculated as the area of the melanophore covering the tailfin mesenchyme. This was quantified by using MATLAB to perform background subtraction of autofluorescence and then used FIJI to threshold on GFP (from MiniCoopR-eGFP) intensity to highlight the pixels that represent melanocytes in each image. All zebrafish were imaged at 3dpf. If treated pharmacologically, treatment started at 1dpf at the indicated concentration for 48 hours and imaged at 3dpf. For experiments involving genetic perturbation using mitfa:Cas9/395, the GFP heart marker in this plasmid was used to screen for zebrafish with successful plasmid integration and expression.

#### Flow *Cytometry of Zebrafish Melanocytes*

##### Zebrafish Embryos

Approximately 250 5dpf embryos were euthanized via tricaine, transferred to an Eppendorf tube (Eppendorf # 022431021), and spun at 500G x 3 minutes at room temperature to remove E3 supernatant. 400μL of trypsin (Invitrogen # 25200-114) was added and incubated for 20 minutes at 28°C. Every 5 minutes an RNase-free disposable pellet pestle (Fisher # 12-141-364) was used to mash the embryos for 2-3 minutes into a single cell solution. After digestion was complete, 500μL of phenol-free Dulbecco’s Modified Eagle Medium (ThermoFisher # 21063029) with 10% FBS (Seradigm) (DMEM10) was added to each sample. The samples were centrifuged 500G x 5 minutes at room temperature and supernatant was gently aspirated. Calcein red stock (Cayman Chemicals # 20632) at 1mM was diluted 1:4000 to desired concentration of 250nM in phenol-free DMEM10 and 500μL was added to embryo pellet and incubated for 30 minutes at 28°C to be used to identify viable cells. Samples were centrifuged again 500G x 5 minutes, supernatant was removed, and pellet was resuspended in phenol-free DMEM with 2% FBS (Seradigm) (DMEM2). Samples were filtered through 40-micron cell-strainer (Thermo Fisher # 08-771-2) two times and then placed in a flow cytometry tube (Fisher # 08-771-23). 1mL of DMEM2 and 1μL of 1000X DAPI (Sigma-Aldrich #D9542-10MG) was added and analyzed on the FACS sorter (BD FACSAria). No color and single-color controls were used to gate for GFP+/Calcein Red+ double positive cells to calculate the frequency of viable melanocytes from the bulk cell suspension. The reporter frequency of viable melanocytes in Figure S3B were calculated on the FACS sorter.

##### Zebrafish Adults

Zebrafish were euthanized via placement in ice-cold water and then dissected to separate body skin and all fins. A clean razor was used to dice the sample into small pieces that can fit through a wide bore p1000 tip (ThermoFisher #2069G). Samples were placed into a 15mL Falcon tube (Fisher 14-959-49D) with 3mL of 1X PBS (Invitrogen 14190-250) and 187.5μL of 2.5mg/mL liberase (Millipore Sigma # 05401020001) and incubated at room temperature for 30 minutes on a shaker to gently keep tissue in suspension. At 15 minutes, a wide bore p1000 tip was used to pipette up and down gently for 3 minutes to dissociate the tissue. After the 30 minutes incubation in liberase at room temperature, 250μL of FBS (Seradigm) was added and then another 3 minutes of pipetting up and down using a wide bore p1000 tip was performed. Cells were then filtered through a 40-micron cell strainer (Thermo Fisher # 08-771-2) into a 50mL conical. Samples were spun at 500G x 5 minutes at 4 °*C* and the supernatant was carefully aspirated using a Pasteur pipette (Fisher #13-678-20D) with low vacuum suction. The pellet was then resuspended in 500μL of PBS with 5% FBS (Seradigm) to be used as flow buffer. Cells were drawn up cells in a p1000 tip with a regular bore size and then fit on 40um filter onto the tip and place in a flow cytometry tube (Fisher # 08-771-23). 0.5μL of 1000X DAPI (Sigma-Aldrich #D9542-10MG) was added and samples were placed on ice. Samples were then FACS sorted (BD FACSAria) for GFP-positive signal and gated based on a GFP-negative control. Each biological replicate represents the pooling of 2 males and 2 female adult zebrafish of the indicated genotype. The reporter frequency of viable melanocytes in Figure 2H were calculated on the FACS sorter.

#### Preparation of Zebrafish Sample for Bulk RNA-seq

##### Zebrafish Embryos

Samples were FACS sorted as described above into 200μL of Trizol (Invitrogen #15596026) in RNase-free LoBind Eppendorf tubes (Eppendorf # 022431021) and snap-frozen by placing on dry ice. Samples were shipped to Genewiz (South Plainfield, NJ), where the RNA isolation and RNA-sequencing was performed.

##### Zebrafish Adult Body Skin and Fins

4 WT stable line fish (2 males and 2 females) and 4 acral melanoma stable line fish (2 males and 2 females) were pooled for each biological replicate. Fish were approximately 6 months post-fertilization and acral melanoma fish had early-stage fin tumors. Samples were processed and FACS sorted as described above into 750μL of Trizol LS (Invitrogen # 10296010) in RNase-free LoBind Eppendorf tubes (Eppendorf # 022431021) and snap-frozen by placing on dry ice. RNA isolation was performed by following Trizol LS protocol. Glycogen (Millipore Sigma #10901393001 was used to co-precipitate the RNA. RNA quality and quantity was measured by Bioanalyzer (Agilent). RNA samples were shipped to Genewiz (South Plainfield, NJ), where RNA-sequencing was performed.

##### Analysis of Zebrafish Bulk RNA-seq

RNA-sequencing was performed using SMART-seq v4 Ultra Low Input RNA Kit (Clonetech). Libraries were constructed using Illumina Nexterna XT kit and were analyzed for concentration by Qubit and for size distribution by Agilent Bioanalyzer. Paired-end sequencing was performed on Illumina HiSeq 2500. After quality control with FASTQC (Babraham Bioinformatics) and trimming with TRIMMOMATIC ^108^ when necessary, reads were aligned to GRCz11 (Ensembl version 96) with transgenes added using STAR ^80^, with quality control via SeQC ^119^. Differential expression was calculated with DESeq2 ^82^ using the output of the --quantMode GeneCounts feature of STAR. The rlog function was used to generate log2 transformed normalized counts Pathway and Gene Ontology (GO) analysis were performed with GSEA using FGSEA-multilevel^83^. Ortholog mapping between zebrafish and human was performed with DIOPT^120^.

Only orthologs with a DIOPT score greater than 6 were used for GSEA and heatmap generation. In cases of more than one zebrafish ortholog of a given human gene, the zebrafish gene with the highest average expression was selected. GSEA comparing the gene expression of GFP+ and GFP-samples was used to validate the melanocyte identity of GFP+ samples (Extended Data Fig. 7b).

#### Zebrafish Single-cell RNA-seq

##### Sample preparation

4 acral melanoma stable line fish (2 males and 2 females) approximately 6 months post-fertilization were utilized and had early-stage fin tumors. All 4 fish were pooled together and dissected into a fin and body skin sample, which were then digested and FACS sorted as described above. The fin and body skin samples were FACS sorted for GFP+ and GFP-populations, sorting 200,000 cells for each group, creating n=4 samples total for scRNA-seq. Cells were FACS sorted into a LoBind Eppendorf (Eppendorf # 022431021) tube containing 600μL DMEM (ThermoFisher # 21063029) supplemented with 20% FBS and 1X penicillin/streptomycin/glutamine and placed on ice. Samples were then centrifuged 300g for 5 minutes in a bucket centrifuge at 4°C. The supernatant was aspirated, and the pellet was resuspended in 40μL of DMEM supplemented with 10% FBS (Seradigm) and 1X penicillin/streptomycin/glutamine to ensure a cell concentration of at least 1000cells/μL. The viability of cells was above 80%, as confirmed with 0.2% (w/v) Trypan Blue staining (Countess II).

##### Cell encapsulation, library preparation, and sequencing

Droplet-based scRNA-seq was performed using the Chromium Single Cell 3’ Library and Gel Bead Kit v3 (10X Genomics) and Chromium Single Cell 3’ Chip G (10X Genomics). Approximately 10,000 cells were encapsulated per each of the four reactions. GEM generation and library preparation was performed according to kit instructions. Libraries were sequenced on a NovaSeq S4 flow cell. Sequencing parameters were: Read1 28 cycles, i5 10 cycles, i7 10 cycles, Read2 90 cycles. Sequencing depth was approximately 40,000 reads per cell. Sequencing data was aligned to our reference zebrafish genome using CellRanger version 5.0.1 (10X Genomics).

##### Analysis

Data was processed using R version 4.0.4 and Seurat version 4.0.3^121^. Each of the four reactions were processed separately before merging into a single object. Cells with fewer than 200 unique genes were filtered out. Expression data was normalized with SCTransform^122^. Principal component analysis^123^ and UMAP dimensionality reduction^124^ were performed using default parameters, with 15 principal components used for UMAP calculations. Clustering was done using the Seurat function FindMarkers with a resolution of 0.2. Clusters were annotated based on expression of zebrafish cell-type specific marker genes as done previously^125, 126^. Expression of *hox13* genes was calculated by summing the normalized expression per cell for each of *hoxa13a, hoxa13b, hoxb13a, hoxc13a, hoxc13b, hoxd13a*.

### Quantification and Statistical Analysis

Statistical comparisons were performed with the aid of GraphPad PRISM 8.4.3 and the statistical details including sample size can be found in the figure legends. A p value of >0.05 is not considered statistically significant. * indicates p<0.05, ** indicates p<0.01, *** indicates p<0.001 and **** indicates p<0.0001.

RNA-seq and ChIP-seq analysis was analyzed in R version 4.0.2 and R Studio version 1.2.5033.

ChIP-seq data was visualized in IGB (BioViz)^127^.

Cunt & Run data was visualized in IGV 2.10.2.

Image analysis performed in Matlab version Update 4 (9.6.0.1150989) and FIJI version 2.0.0.

