## Supplementary material for "Anatomic position determines oncogenic specificity in melanoma": Table S1 - CAMP criteria for melanoma diagnosis in transgenic zebrafish models

**Supplemental Table 1: CAMP criteria for diagnosis in zebrafish models of melanoma** Related to Methods. CAMP criteria: Crestin positivity, Anatomic localization, *Mitfa* expression, Pigmentation. Schematic shows the decision tree used in calling lesions a melanoma vs not a melanoma in both acral and cutaneous melanoma models. These criteria were validated by performing histology on fish from both acral and cutaneous melanoma models.

**Supplemental Table 1: CAMP Criteria for Diagnosis in Zebrafish Models of Melanoma**
(**C**restin positivity, **A**natomic localization, **M**ITF expression, **P**igmentation)

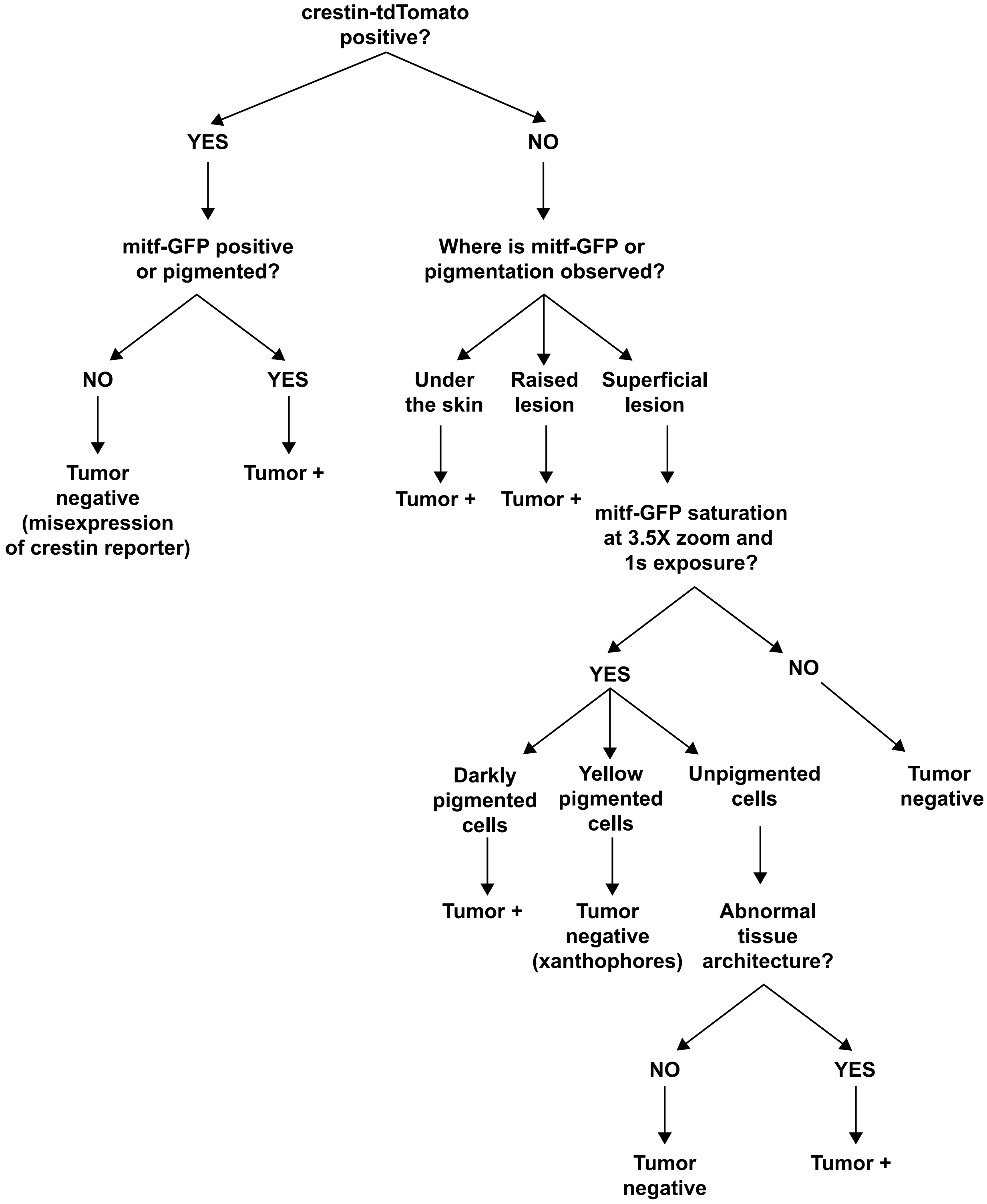
