## Supplementary material for "Anatomic position determines oncogenic specificity in melanoma": Table S2 - Primer Sequences

### Table S2. Related to Methods: primer sequences

| Primer | Sequence (5’-3’) |
|---|---|
| Acral genotype F | GTTGAACGCAAGTTTGTACA |
| hsCRKL genotype R | TGATGTAGTGGGAGACCCGC |
| hsGAB2 genotype R | AGTATTCCAGAACATCTGGG |
| hsTERT genotype R | AGGCAGGACACCTGGCGGAA |
| Cas9-mCherry | CATGTGCACCTTGAAGCGCA |
| zebrafish nf1a CRISPR-seq F | TCGGGATCGCAAAAGTGATT |
| zebrafish nf1a CRISPR-seq R | CACCAAGCTCACATCTTCAA |
| zebrafish nf1b CRISPR-seq F | CACCATCTTCATCATCCTCCT |
| zebrafish nf1b CRISPR-seq F | ACTACTCTCTGTCCCGTGTC |
| zebrafish mitfa RT-PCR F | GCCCTATGGCCCTTCTCAC |
| zebrafish mitfa RT-PCR R | CATCCATGAACCCAAGAATGTCA |
| hsCRKL RT-PCR F&R | Sino Biological cat #HP101145 |
| hsGAB2 RT-PCR F | GCGGCGACGTGGTGT |
| hsGAB2 RT-PCR R | CTTCCAGGCATAGCGCCTC |
| hsTERT RT-PCR F | GGAGCAAGTTGCAAAGCATTG |
| hsTERT RT-PCR R | TCCCACGACGTAGTCCATGTT |
| zebrafish beta-actin RT-PCR F | GCCAACAGAGAGAAGATGACAC |
| zebrafish beta-actin RT-PCR R | CAGAGAGAGCACAGCCTGG |
| zebrafish hoxa13a Surveyor F | GGGTGATTCTGGAAAGCAAT |
| zebrafish hoxa13a Surveyor R | CTCCATGGGATACTGACTCT |
| zebrafish hoxa13b Surveyor F | ATCCCATTGTGCAATGGAAT |
| zebrafish hoxa13b Surveyor R | CGCCAAAATATCCATAGGGC |
| zebrafish hoxb13a Surveyor F | TTGACATTCTTCACCCAAGG |
| zebrafish hoxb13a Surveyor R | AGCCCTGGTAGGATATTCTT |
| zebrafish hoxc13a Surveyor F | GCCAGTAGTTGTTTAAAGGG |
| zebrafish hoxc13a Surveyor R | GTCTCGGCATATTTTTCTGC |
| zebrafish hoxc13b Surveyor F | TAGTGAAAGACGTTTGCGTT |
| zebrafish hoxc13b Surveyor R | ACAACTGGGACGTCCAAATA |
| zebrafish hoxd13a Surveyor F | ATGCACTGAGGAATATGGAC |
| zebrafish hoxd13a Surveyor R | CTAATGAAGAGAGGCGAGGA |
| zebrafish hoxb7a Surveyor F | ATATATCATCACGTGCTGCC |
| zebrafish hoxb7a Surveyor R | CTCTACATACACAGACGCAC |
| hsCRKL gRNA-1 Surveyor F | TTCGAATAAACATGTCGTCTGC |
| hsCRKL gRNA-1 Surveyor R | CGCCTATCTTGAATCTCTTGCT |
| hsCRKL gRNA-2 Surveyor F | TCCACTTGTCCTGGTGATTATG |
| hsCRKL gRNA-2 Surveyor R | TTCTATCAGGGTGGTCGTATCC |
| hsCRKL gRNA-3 Surveyor F | CTCCACTAGCGAGAAGCTGATT |
| hsCRKL gRNA-3 Surveyor R | TCTTGAAAGGAAGGTCTTCAGC |
| zebrafish igf1 Surveyor F | GGATTTTCTCTCCAAATCCG |
| zebrafish igf1 Surveyor R | CTTAATCATGTCGACTCAGC |
| zebrafish igf2a Surveyor F | CAAAAGAACCACTCGTTCAC |
| zebrafish igf2a Surveyor R | CACACGAACTGCAGTGTATC |
| zebrafish igf2b Surveyor F | CTCTCAGAGAACTTTTGCCT |
| zebrafish igf2b Surveyor R | GCGTATCCAGAACGTAATGT |
